# Quantifying genetic effects on disease mediated by assayed gene expression levels

**DOI:** 10.1101/730549

**Authors:** Douglas W. Yao, Luke J. O’Connor, Alkes L. Price, Alexander Gusev

## Abstract

Disease variants identified by genome-wide association studies (GWAS) tend to overlap with expression quantitative trait loci (eQTLs). However, it remains unclear whether this overlap is driven by mediation of genetic effects on disease by expression levels, or whether it primarily reflects pleiotropic relationships instead. Here we introduce a new method, mediated expression score regression (MESC), to estimate disease heritability mediated by the cis-genetic component of assayed steady-state gene expression levels, using summary association statistics from GWAS and eQTL studies. We show that MESC produces robust estimates of expression-mediated heritability across a wide range of simulations. We applied MESC to GWAS summary statistics for 42 diseases and complex traits (average *N* = 323K) and cis-eQTL data across 48 tissues from the GTEx consortium. We determined that a statistically significant but low proportion of disease heritability (mean estimate 11% with S.E. 2%) is mediated by the cis-genetic component of assayed gene expression levels, with substantial variation across diseases (point estimates from 0% to 38%). We further partitioned expression-mediated heritability across various gene sets. We observed an inverse relationship between cis-heritability of expression and disease heritability mediated by expression, suggesting that genes with weaker eQTLs have larger causal effects on disease. Moreover, we observed broad patterns of expression-mediated heritability enrichment across functional gene sets that implicate specific gene sets in disease, including loss-of-function intolerant genes and FDA-approved drug targets. Our results demonstrate that eQTLs estimated from steady-state expression levels in bulk tissues are informative of regulatory disease mechanisms, but that such eQTLs are insufficient to explain the majority of disease heritability. Instead, additional assays are necessary to more fully capture the regulatory effects of GWAS variants.

## Introduction

In the past decade, genome-wide association studies (GWAS) have shown that most disease-associated variants lie in noncoding regions of the genome^1–3^, leading to the hypothesis that regulation of gene expression levels is the primary biological mechanism through which genetic variants affect complex traits, and motivating large scale expression quantitative trait loci (eQTL) studies^4, 5^. Many statistical methods have been developed to integrate eQTL data with GWAS data to gain functional insight into the genetic architecture of disease. These methods fall into two classes. The first class operates on individual genes and includes colocalization tests^6–10^, which identify genes with shared causal variants for their expression levels and disease, and transcriptome-wide association studies^11–16^, which identify genes with significant cis-genetic correlations between their expression and disease. The second class of methods operates on the entire genome and partitions disease heritability by SNP categories defined by eQTL status (i.e. whether or not SNPs are eQTLs)^17–20^. Application of these methods to eQTL and GWAS data has shown that many genes have eQTLs that colocalize with GWAS loci^6–10^ and/or exhibit significant cis-genetic correlations between their expression and trait^11–16, 21–28^, while also showing that eQTLs as a whole are significantly enriched for disease heritability^17–20^.

However, the true causal relationships between SNPs, gene expression levels, and disease remain uncertain for the vast majority of genes^9, 29–31^. This uncertainty arises from the fact that several different causal scenarios can result in similar patterns of enrichment/overlap between GWAS loci and eQTLs, summarized as follows: (1) mediation, in which a SNP affects gene expression levels, which then affect disease risk; (2) pleiotropy, in which a SNP affects gene expression levels and independently affects disease risk through an alternative mechanism; and (3) linkage, in which a SNP that affects gene expression levels is in linkage disequilibrium (LD) with a different, independent disease SNP (Figure 1a). These scenarios carry very different biological interpretations. In particular, only scenario (1) is informative of the SNP’s mechanism of action on disease, so it is vital to distinguish scenarios (2) and (3) from scenario (1). However, existing methods are unable to do so consistently. Among the individual-gene methods mentioned above, colocalization tests can sometimes rule out linkage as an explanation for overlap between eQTLs and disease SNPs, but cannot rule out pleiotropy^13, 32^, while transcriptome-wide association studies cannot rule out either pleiotropy or linkage^13, 29^, especially for genes with only one strong eQTL^15^. Among the genome-wide methods, some aim to rule out linkage through fine-mapping of eQTLs^19^, but none aim to rule out pleiotropy. Thus, it remains unclear whether enrichment/overlap between eQTLs and disease SNPs usually reflects mediation, or whether it more commonly reflects pleiotropy and/or linkage^9, 29^. For example, in the case of autoimmune diseases, most instances of overlap between significant disease loci and immune cell eQTLs are driven by linkage^9^, suggesting that linkage may be more prevalent than mediation^31^.

**Figure 1:**
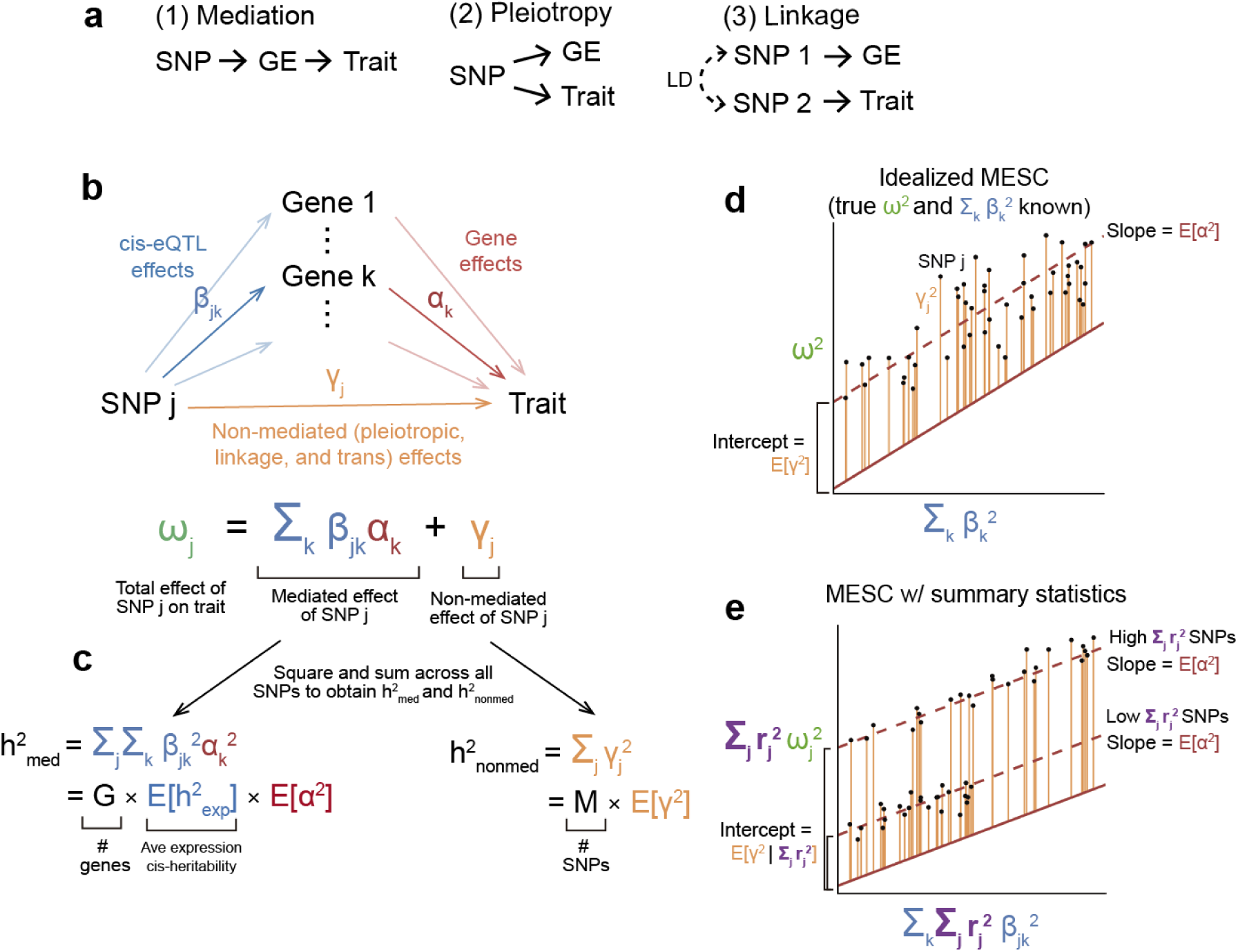
Schematic of MESC. (**a**) Different models of causality between SNP, gene expression levels (GE), and trait. (**b**) SNP effect sizes are modeled as the sum of a mediated component (defined as cis-eQTL effect sizes *β* multiplied by gene-trait effect sizes *α*) and a non-mediated component *γ*. (**c**) Heritability mediated by the cis-genetic component of gene expression levels 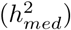 is defined as the squared mediated component of SNP effect sizes summed across all SNPs (assuming that genotypes and phenotypes are standardized). 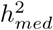 can be rewritten as the product of the number of genes *G*, the average expression cis-heritability 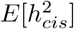, and the average gene-trait effect size *E*[*α*^2^] (**d**) The basic premise behind MESC is to regress squared GWAS effect sizes on squared eQTL effect sizes, in which case the slope will equal *E*[*α*^2^] given appropriate effect size independence assumptions (see Methods). (**e**) In practice, MESC involves regressing squared GWAS summary statistics on squared eQTL summary statistics. Differences in the level of LD between SNPs are captured by an LD score covariate. In the figure, we show a simplified LD architecture with two discrete levels of LD.

In this study, we aim to quantify the proportion of disease heritability that is specifically mediated in cis by the assayed expression levels of the set of all genes, and of genes in specific functional categories (scenario 1 from above). We first define expression-mediated heritability under a generative model featuring both mediated and non-mediated (including pleiotropic and linkage) effects of SNPs on the trait. We introduce a method, mediated expression score regression (MESC), to estimate expression-mediated heritability from GWAS summary statistics, ancestry-matched LD scores, and eQTL effect sizes obtained from external expression panels. Intuitively, MESC distinguishes mediated from non-mediated effects in a set of genes via the idea that mediation (unlike pleiotropy and linkage) induces a linear relationship between the magnitude of eQTL effect sizes and disease effect sizes.

## Results

### Definition of expression-mediated heritability

We first define heritability mediated by the cis-genetic component of gene expression levels 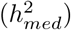, which is the quantity that our method aims to estimate. We model individual SNP effect sizes on a disease as the sum of a mediated component, defined as causal cis-eQTL effect sizes (where “cis” includes SNPs located within 1 Mb of the gene) multiplied by gene-trait effect sizes, and a non-mediated component, which includes pleiotropic, linkage, and trans-eQTL-mediated effects (Figure 1b). Under our model, 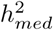 is defined as the squared mediated component of each SNP effect size summed across all SNPs (Figure 1c). We consider additional causality scenarios, such as reverse mediation, cis-by-trans mediation, and mediation by unobserved intermediaries, and we justify that these scenarios do not compromise our definition of 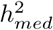 (see “Modes of expression causality” in Supplementary Note).

In the above definition of 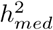, which we call 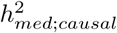, we assume that cis-eQTL effect sizes are taken in the *causal cell types/contexts for the disease*, which is the natural setting in which we conceptualize a model of mediation via gene expression levels. However, we typically cannot measure expression in causal cell types/contexts for a given disease and thus cannot directly estimate 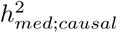. Instead, we rely on *assayed* expression from an external expression panel such as GTEx^5^ to act as a proxy to expression in causal cell types/contexts. We define the heritability mediated by *assayed* expression levels in a set of tissues *T* 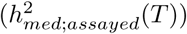 as follows:

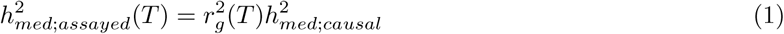

where 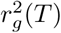 is the average squared genetic correlation between expression in *T* and expression in the causal cell types/contexts for the disease. 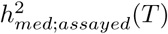 represents the final quantity that we estimate in practice, and it reflects two quantities: (1) the amount of mediation occurring in true causal cell types/contexts for the disease, as captured by 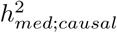, and (2) the extent to which eQTL effect sizes in *T* are correlated with eQTL effect sizes in causal cell types/contexts for the disease, as captured by 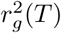. For brevity, we refer to 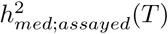 as 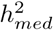 for the remainder of the manuscript, where the set of tissues *T* is implicit.

In addition to the heritability mediated by the expression levels of *all* genes, we define a quantity 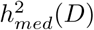 with respect to *gene category D*, where *D* can be arbitrarily defined over any set of genes (e.g. genes in a specific molecular pathway or interaction network). 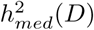 corresponds to the heritability mediated by the expression levels of only genes in *D* (Methods).

### Estimating expression-mediated heritability using MESC

In order to estimate 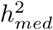, we first decompose 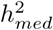 into the product of three quantities: the number of genes, the average cis-heritability of expression across all genes, and the average squared gene-trait effect size across all genes (Fig 1c). The average cis-heritability of expression across genes can be readily estimated with existing methods^33, 34^. In order to estimate the average squared gene-trait effect size across all genes, we propose an approach that involves regressing squared SNP-trait effect sizes on squared cis-eQTL effect sizes summed across genes (Figure 1d). We make two extensions to this simplified regression model. First, we model tagging effects due to LD between SNPs, which allows us to perform this regression using summary statistics from GWAS and eQTL studies (Figure 1e). Second, to avoid bias (see below), we stratify the regression across both gene categories *D* and SNP categories *C* (Methods). The regression equation used to estimate 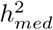 is

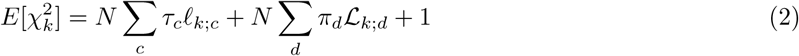

where 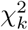 is the GWAS *χ*^2^-statistic of SNP *k, N* is the number of samples, *τ*_*c*_ is the per-SNP contribution to non-mediated heritability of SNPs in SNP category *C, ℓ*_*k*;*c*_ is the *LD score*^2, 34^ of SNP *k* with respect to SNP category *C* (defined as 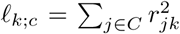), *π*_*d*_ is the per-gene contribution to 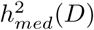, and ℒ_*k*;*d*_ is the *expression score* of SNP *k* with respect to gene category *D* (defined as 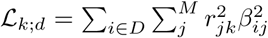). Note that MESC can accommodate both overlapping gene categories and overlapping SNP categories. ℒ_*k*;*d*_ can be conceptualized as the total expression cis-heritability of genes in *D* that is tagged by SNP *k*.

Equation (2) allows us to estimate *π*_*d*_ and *τ*_*c*_ via computationally efficient multiple regression of GWAS chi-square statistics against LD scores and expression scores. In order for equation (2) to provide unbiased estimates of 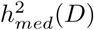, the following key assumptions must be met:

1. Within each gene category *D*, the magnitude of eQTL effect sizes is uncorrelated with the magnitude of gene-trait effect sizes
2. Within each SNP category *C*, the magnitude of eQTL effect sizes is uncorrelated with the magnitude of non-mediated effect sizes

Note that if we estimate total 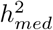 without stratifying genes and SNPs, these two assumptions are likely violated in practice, leading to biased estimates of 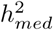. The first assumption is violated in the scenario that genes under selection systematically have low expression heritability and large effect on disease^5, 35^, resulting in a negative correlation between eQTL and gene effect size magnitude and thus causing 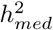 to be underestimated. We can ameliorate this bias by stratifying genes in a manner that captures the dependence between eQTL effect sizes and gene-trait effect sizes across the genome (e.g. stratifying genes by the magnitude of their expression cis-heritability). The second assumption from above is violated in the scenario that the magnitude of non-mediated SNP effect sizes and eQTL effect sizes tend to both be larger in biologically active regions of the genome^36–39^ (e.g. promoters, enhancers), resulting in a positive correlation between eQTL and non-mediated effect size magnitude and thus causing 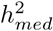 to be overestimated. We can ameliorate this bias by stratifying SNPs in a manner that captures the dependence between eQTL effect sizes and non-mediated effect sizes across the genome (e.g. stratifying SNPs according to whether they lie in promoters, enhancers, etc.). In summary, stratifying the regression over strategically defined SNP and gene categories is necessary for us to obtain unbiased estimates of 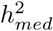 in practice due to violations to the two above assumptions (see below for simulation results and “Model assumptions” in Methods for additional discussion).

In practice, we estimate LD scores from an external reference panel. Although expression scores can be directly estimated from cis-eQTL summary statistics, we obtain less noisy estimates from individual-level genotypes and gene expression measurements (Supplementary Note); thus, we estimate expression scores from individual-level data when it is available^11^. We do not exclude genes from our analyses based on any significance threshold, in contrast to previous approaches^9–13, 19^. When applicable, we meta-analyze expression scores across tissues (Methods).

Throughout this study, we present estimates of three quantities that are a function of 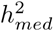 and/or 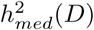. These quantities include the proportion of heritability mediated by expression (defined as 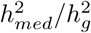), the proportion of expression-mediated heritability for gene category *D* (defined as 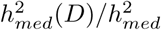), and the enrichment of expression-mediated heritability for *D* (defined as the proportion of expression-mediated heritability in *D* divided by the proportion of genes in *D*). We estimate standard errors and p-values for all quantities by block jackknife^2, 34^. When applicable, we perform random effects meta-analysis of these quantities across diseases and complex traits (Methods). We have released open source software implementing our method (URLs).

### Simulations assessing calibration and bias

We performed simulations to assess the calibration and bias of MESC in estimating 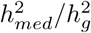 and its standard error from simulated complex trait and expression data under a variety of genetic architectures (Methods). We performed all simulations using real genotypes from UK Biobank^40^ (*N*_*GW AS*_ = 10,000 samples; *M* = 98,499 SNPs from chromosome 1). We simulated causal cis-eQTL effect sizes for *G* = 1000 genes, gene effect sizes on a complex trait, and non-mediated effect sizes of each SNP on the complex trait from normal or point-normal distributions corresponding to various levels of 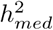 and 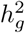. We then simulated complex trait phenotypes from these effect sizes, with normally distributed environmental effects. To emulate an external expression panel, we simulated expression phenotypes from the eQTL effect sizes using a separate set of genotypes, with normally distributed environmental effects (*N*_*eQTL*_ = 100-1000 samples). For all simulations other than Figure 2c, eQTL effect sizes used to generate expression panel phenotypes and complex trait phenotypes were identical (i.e. 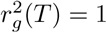).

**Figure 2:**
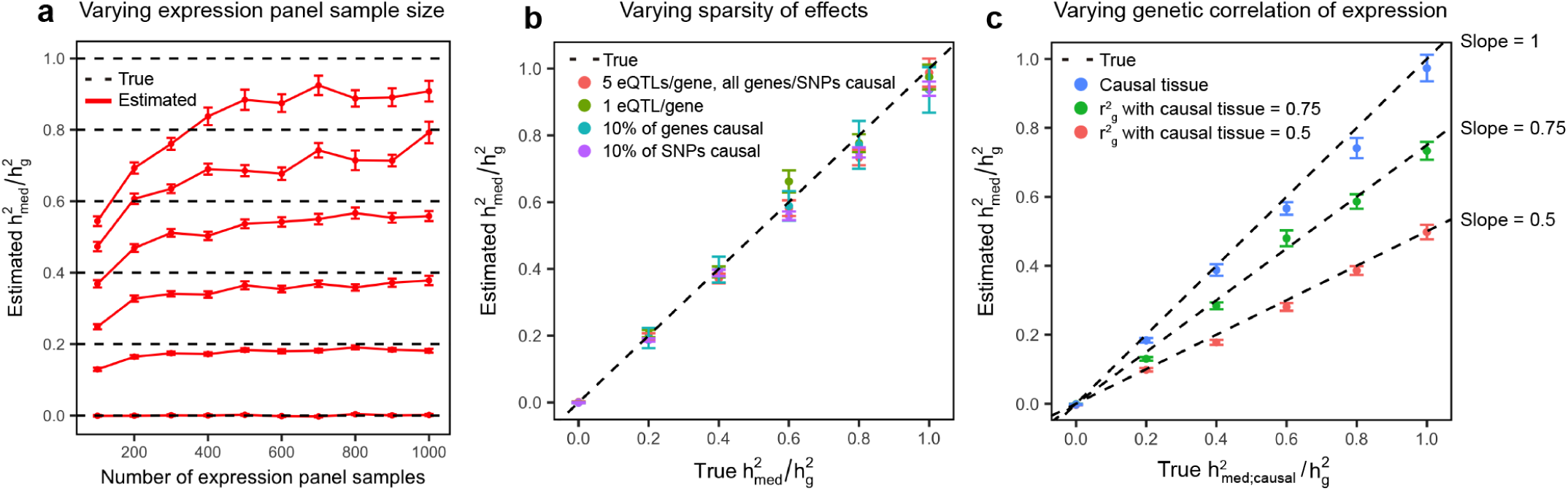
Simulation results assessing bias of MESC. We simulated expression and complex trait architectures corresponding to various levels of 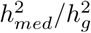. GWAS sample size was fixed at 10,000 and 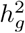 was fixed at 0.5. Error bars represent mean standard errors across 300 simulations. (**a**) Impact of expression panel sample size on estimates of 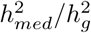. Expression scores were estimated from simulated expression panel samples using LASSO with REML correction. See Supplementary Figure 1 for estimates of 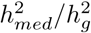 using other methods of estimating expression scores. (**b**) Impact of sparse genetic/eQTL architectures on estimates of 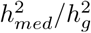. Expression panel sample size was set to 1,000. (**c**) Estimates of 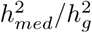 with 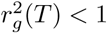. Expression panel sample size was set to 1,000.

First, we evaluated the impact of expression panel sample size and different methods of estimating expression scores on the bias of MESC. We used five different methods to estimate expression scores from simulated expression panels with sample sizes ranging from 100 to 1000. These methods include: (1) eQTL summary statistics, (2), LASSO^41^, (3) LASSO with REML correction, (4) BLUP^42^, and (5) BLUP with REML correction (see “Comparing different methods of estimating expression scores” in Supplementary Note for description of these methods). We then used these expression scores to estimate 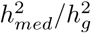 from the simulated complex trait phenotypes. Of the five methods, we observed that LASSO with REML correction gave the best performance, providing approximately unbiased estimates of 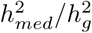 with around 500 or more expression panel samples (Figure 2a). LASSO outperformed other methods due to the sparsity prior it places on effect sizes, which matches the sparse nature of our simulated cis-eQTL effect sizes. Meanwhile, because LASSO produces biased effect size estimates, scaling the effect sizes to match the REML-predicted expression cis-heritability is necessary to obtain effect size estimates that are unbiased (Supplementary Note). All other methods of estimating expression scores produced biased results at all sample sizes tested (Supplementary Figure 1) due to large sampling noise and/or systematic bias in their estimates of expression scores (Supplementary Figure 2). Notably, all methods (including LASSO with REML correction) produced biased estimates of 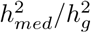 at *N*_*eQTL*_ comparable to individual tissue expression panels (*N*_*eQTL*_ < 200). To evaluate the performance of our expression score meta-analysis procedure (Methods), we performed additional simulations in which we estimated 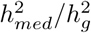 using expression scores meta-analyzed across multiple simulated tissues with identical eQTL effect sizes and independent environmental noise. We obtained approximately unbiased estimates of 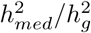 with 5 tissues × 200 samples per tissue (Supplementary Figure 3), demonstrating that we were able to ameliorate low-*N*_*eQTL*_ dependent downward bias in 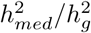 via meta-analysis of expression scores across tissues.

To test the bias of MESC in sparse genetic/gene architectures, we performed simulations in which we varied the number of eQTLs per gene, the proportion of SNPs with non-mediated effects, and the proportion of genes with gene-trait effects. We observed that MESC was robust to the number of eQTLs per gene and sparsity of gene effect sizes or non-mediated effect sizes (Figure 2b). To test the bias of MESC across different levels of total disease heritability, we performed simulations in which we varied 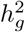 from 0.05 to 1, obtaining unbiased estimates of 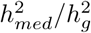 for all values of 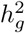 (Supplementary Figure 4)

To test the bias of MESC in frequency-dependent genetic architectures^43–45^, we performed simulations in which eQTL and non-mediated per-allele effect size magnitude were inversely proportional to minor allele frequency (Supplementary Note), consistent with purifying selection acting on gene expression^46, 47^ and complex trait^44, 45^. We obtained nearly unbiased estimates of 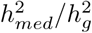 across diverse frequency-dependent genetic architectures for both gene expression and disease when stratifying SNPs by 10 minor allele frequency bins^43^ (Supplementary Figure 5).

Next, we tested the calibration of jackknife standard errors computed by MESC for the 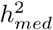 enrichment of a gene category *D*, defined as (proportion of 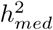 in D) / (proportion of genes in D). This is the main quantity that we aim to estimate for analyses involving gene sets. We simulated a gene category containing 200 genes. We then performed two sets of simulations in which this gene category was either enriched or not enriched for 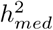. We observed well-calibrated 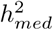 enrichment standard errors for the gene category under both the null and causal enrichment scenarios (Supplementary Figure 6).

Finally, we sought to verify that MESC estimates the quantity 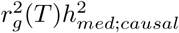 when estimating expression scores from a tissue with squared genetic correlation 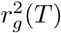 with the unobserved causal tissue. For purposes of this simulation, we assumed a single causal tissue. We simulated expression phenotypes for assayed tissues that were genetically correlated with expression in the causal tissue 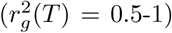, then predicted expression scores from the assayed tissues and estimated 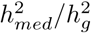 from these expression scores (Methods). We obtained unbiased estimates of 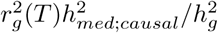 for each value of 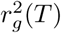 (Figure 2c).

In summary, we show that MESC produces approximately unbiased estimates of 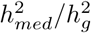 and well-calibrated standard errors under a wide variety of genetic architectures at *N*_*eQTL*_ > 500. Given that many individual tissue expression panels have *N*_*eQTL*_ of 200 or fewer^5^, this suggests that meta-analysis of expression scores across tissues is likely necessary to obtain unbiased estimates of 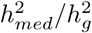 in practice. Note that all simulations in this section were conducted assuming that the two main effect size independence assumptions were satisfied across the genome; see below for simulations under violations of these assumptions.

### Simulations under violations of effect size independence assumptions

We performed simulations to assess the bias of MESC in the presence of violations to the two main effect size independence assumptions, and to assess how well partitioning genes and SNPs ameliorated this bias (see “Model assumptions” in Methods for discussion of these assumptions).

In order to simulate violations to gene-eQTL effect size independence, we simulated eQTL effect sizes and gene effect sizes so that the per-gene 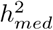 was constant across genes, but the magnitude of eQTL effect sizes and gene effect sizes were strongly inversely proportional across genes (Methods). This emulates the scenario in which negative selection causes large-effect eQTLs for large effect genes to be selected out of the population, which in turn causes low-heritability genes to have larger gene-trait effects. Non-mediated effect sizes were simulated in the same manner as previous simulations. As expected, when estimating 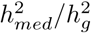 in this scenario using expression scores corresponding to a single gene category, we obtained downwardly biased estimates of 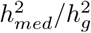. When we estimated 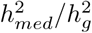 with genes stratified by their estimated expression cis-heritability into 5 gene categories *D*, we obtained approximately unbiased estimates of 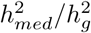 (Figure 3a), as well as approximately unbiased estimates of 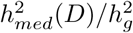 for each *D* (Supplementary Figure 7). This result demonstrates that partitioning genes into expression cis-heritability bins can capture a continuous underlying relationship between eQTL effect sizes and gene-trait effect sizes across the genome, enabling us to obtain unbiased estimates of 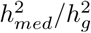 in the presence of violations to gene-eQTL effect size independence.

**Figure 3:**
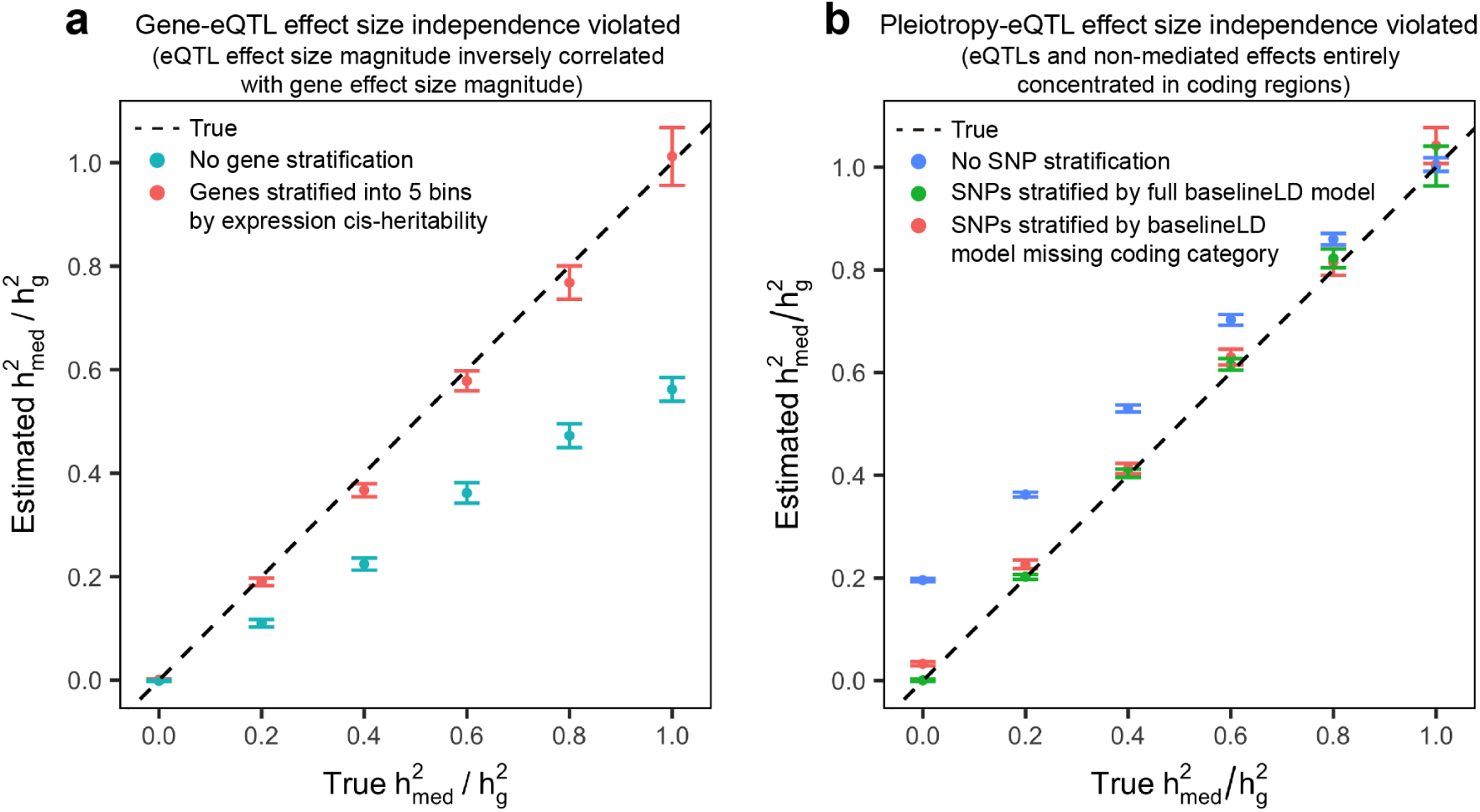
Simulation results for violations of effect size independence assumptions. Error bars represent mean standard errors across 300 simulations. (**a**) We simulated eQTL effect sizes and gene effect sizes so that the per-gene 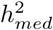 was constant across genes, but the magnitude of eQTL effect sizes and gene effect sizes were strongly inversely proportional across genes. (**b**) We simulated 100% of eQTL effects and non-mediated effects to lie within coding regions. Results in which all eQTL effects and non-mediated effects lie in conserved regions or transcription start sites can be found in Supplementary Figure 9.

In order to simulate violations to pleiotropy-eQTL effect size independence, we performed simulations in which specific SNP categories were enriched for both eQTL effects and non-mediated effects. This emulates the presence of regulatory hotspots in the genome with a high concentration of both eQTL effects and non-mediated effects, and it induces a genome-wide positive correlation between eQTL effect size magnitude and non-mediated effect size magnitude. We selected three functional SNP categories from the baselineLD model^2, 43^—transcription start sites, coding regions, and conserved regions—that have been shown to be highly enriched for both expression cis-heritability^48^ and complex trait heritability^2^. We then simulated 10x expression cis-heritability enrichment and 10x total heritability enrichment for SNPs in any of these three categories, which is close to empirical estimates^2, 48^. Gene effect sizes were simulated in the same manner as previous simulations, and we estimated 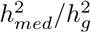 using a single SNP category containing all SNPs. However, we observed no discernible upward bias in these estimates of 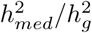 (Supplementary Figure 8), demonstrating that stronger violations to pleiotropy-eQTL effect size independence are necessary to induce bias in estimates of 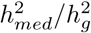.

In order to simulate more extreme patterns of colocalization between eQTL effects and non-mediated effects, we performed three sets of simulations in which 100% of eQTLs and disease heritability were entirely concentrated in each of the three SNP categories. When we estimated 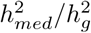 for each simulation using a single SNP category containing all SNPs, we obtained upwardly biased estimates of 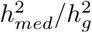 (Figure 3b). We then estimated 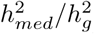 for each simulation in two additional ways: (1) with SNPs stratified by the full baselineLD model, and (2) with SNPs stratified by a misspecified baselineLD model missing the causal category and window around the causal category, emulating a realistic scenario in which the SNP model does not fully capture all sources of non-mediated effects. When we estimated 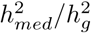 with SNPs stratified by the full baselineLD model, we obtained unbiased estimates of 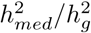. When we estimated 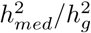 with SNPs stratified by the misspecified baselineLD model, we obtained estimates of 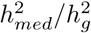 that were nearly unbiased (Figure 3b, Supplementary Figure 9), demonstrating that the remaining SNP categories in the baselineLD model were able act as a reasonable proxy to the missing causal category. In summary, we show that partitioning SNPs by the baselineLD model can account for biases in 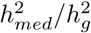 caused by non-independence between eQTL effect sizes and non-mediated effect sizes, even in highly exaggerated scenarios e.g. when 100% of heritability is concentrated in a single SNP category.

### Comparison to other methods in simulations

To our knowledge, no published methods specifically aim to estimate heritability mediated by expression levels. The closest analogues are approaches that measure the genome-wide heritability enrichment of eQTLs^17–20^ using GCTA^33^ or stratified LD score regression (S-LDSC)^2, 43^. If pleiotropy-eQTL effect size independence holds *across the genome*, then the heritability enrichment of a SNP category corresponding to the set of all eQTLs will accurately reflect 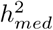 (i.e. the heritability enrichment will be 1x when 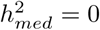 and will increase when 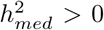). Given pleiotropy-eQTL effect size independence, we observed that S-LDSC has a well-calibrated false positive rate for detecting heritability enrichment of the eQTL category when 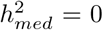, with increasing power to detect heritability enrichment of the eQTL category as 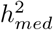 increased (Supplementary Figure 10).

However, in the presence of violations to pleiotropy-eQTL effect size independence, S-LDSC will detect significant heritability enrichment of the eQTL category even in the absence of mediation. Like the simulation performed in Supplementary Figure 8, we simulated 10x enrichment of eQTL effect sizes in three SNP categories (coding regions, transcription start sites, and conserved regions). With 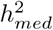 fixed at 0, we then varied the heritability enrichment of the three SNP categories from 2.5x to 10x. We observed that S-LDSC detected significant heritability enrichment of the eQTL category in the absence of mediation when total heritability enrichment was 5x or greater, whereas MESC had a well-calibrated false positive rate at all levels of enrichment (Figure 4). Note that this result does not imply that S-LDSC or other heritability partitioning methods are flawed, but rather that they cannot specifically distinguish mediated effects from non-mediated effects when they are applied to annotations generated from eQTL data. We did not compare MESC to the colocalization methods of Chun et al.^9^ or Ongen et al.^10^, since these methods only operate on genome-wide significant GWAS loci and also do not attempt to distinguish pleiotropy from mediation.

**Figure 4:**
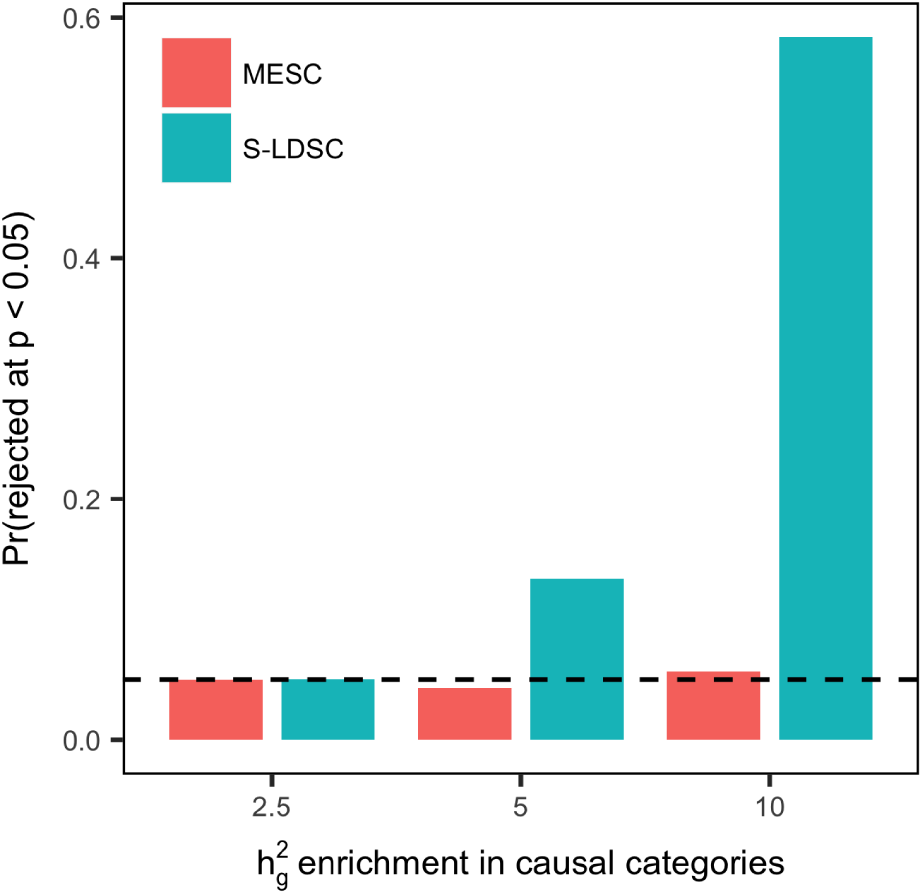
Comparison of MESC to heritability partitioning. We fixed 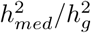 at 0 for all simulations. We simulated 10x enrichment of eQTL effect sizes for SNPs within three causal SNP categories (coding regions, transcription start sites, and conserved regions). We then simulated heritability enrichment in the same three SNP categories ranging from 2.5x to 10x. In the figure, we show the proportion of simulations in which the null hypothesis that 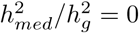 is rejected by MESC, and the proportion of simulations in which the null hypothesis of no 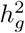 enrichment for the set of all eQTLs is rejected by stratified LD-score regression (S-LDSC). 300 simulations were performed.

### Estimation of expression-mediated heritability for 42 diseases and complex traits

We applied MESC to estimate the proportion of heritability mediated by the cis-genetic component of assayed expression levels 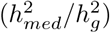 for 42 independent diseases and complex traits from the UK Biobank^40^ and other publicly available datasets (average *N* = 323K; see Supplementary Table 1 for list of traits). We obtained estimates of expression cis-heritability for each gene in each of 48 tissues from the GTEx consortium^5^ using GCTA^33^. We estimated individual-tissue expression scores from raw expression data (with appropriate quality control steps) and individual-level genotypes in 48 tissues from GTEx. (Methods; see Supplementary Table 2 for list of tissues). In contrast to approaches that restrict to genes with significantly heritable expression^11, 12^, we included in our analysis all gene-tissue pairs for which LASSO converged when estimating eQTL effect sizes, resulting in an average of 13,674 genes analyzed per tissue (see Supplementary Table 2 for number of genes per tissue). To capture dependencies between eQTL effect sizes and non-mediated effect sizes and avoid upward bias in 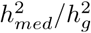 estimates, we partitioned SNPs by 72 functional categories specified by the baselineLD v2.0 model^2, 43^ (Methods). To capture dependencies between eQTL effect sizes and gene-trait effect sizes and avoid downward bias in 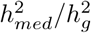 estimates, we partitioned genes into 5 bins based on the magnitude of their estimated expression cis-heritability. Notably, we restricted all our analyses to Hapmap3 SNPs^49^, in which case we estimated the proportion of *common* disease heritability mediated by gene expression levels (see Supplementary Note for discussion of rare vs. common variant 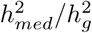). Finally, we meta-analyzed expression scores in two ways: within groups of GTEx tissues with common biological origin, and across all 48 GTEx tissues (Methods). For each trait, we used MESC to estimate three versions of 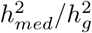 using three different types of expression scores: (1) individual-tissue expression scores, (2) tissue-group meta-analyzed expression scores, and (3) all-tissue meta-analyzed expression scores.

Across all traits, we observed an average 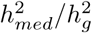 of 0.11 (S.E. 0.02) from all-tissue meta-analyzed expression scores. We did not observe a relationship between 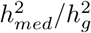 and 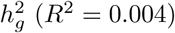 (Supplementary Figure 11). Of the 42 traits, 26 had 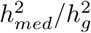 estimates greater than 0 at nominal significance (p-value < 0.05), with 10 reaching Bonferroni significance (p-value < 0.05 / 42). In Figure 5a, we report 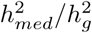 estimates from all-tissue and tissue-group meta-analyzed expression scores for a representative set of 10 genetically uncorrelated traits. 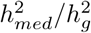 estimates for all 42 traits from all three types of expression scores can be found in Supplementary Figure 12 and Supplementary Table 4. Between the three types of expression scores, we observed the lowest estimates of 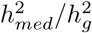 from individual-tissue expression scores (Figure 5b, Supplementary Figure 12). Moreover, we observed a positive correlation between the sample size of the tissue and magnitude of the estimated 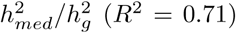 (Supplementary Figure 13). These results demonstrate that tissue sample size affected our ability to obtain unbiased estimates of 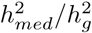 from individual-tissue expression scores, consistent with our simulations that suggest downward bias in 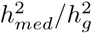 estimates at *N*_*eQTL*_ < 200 (Figure 2a).

**Figure 5:**
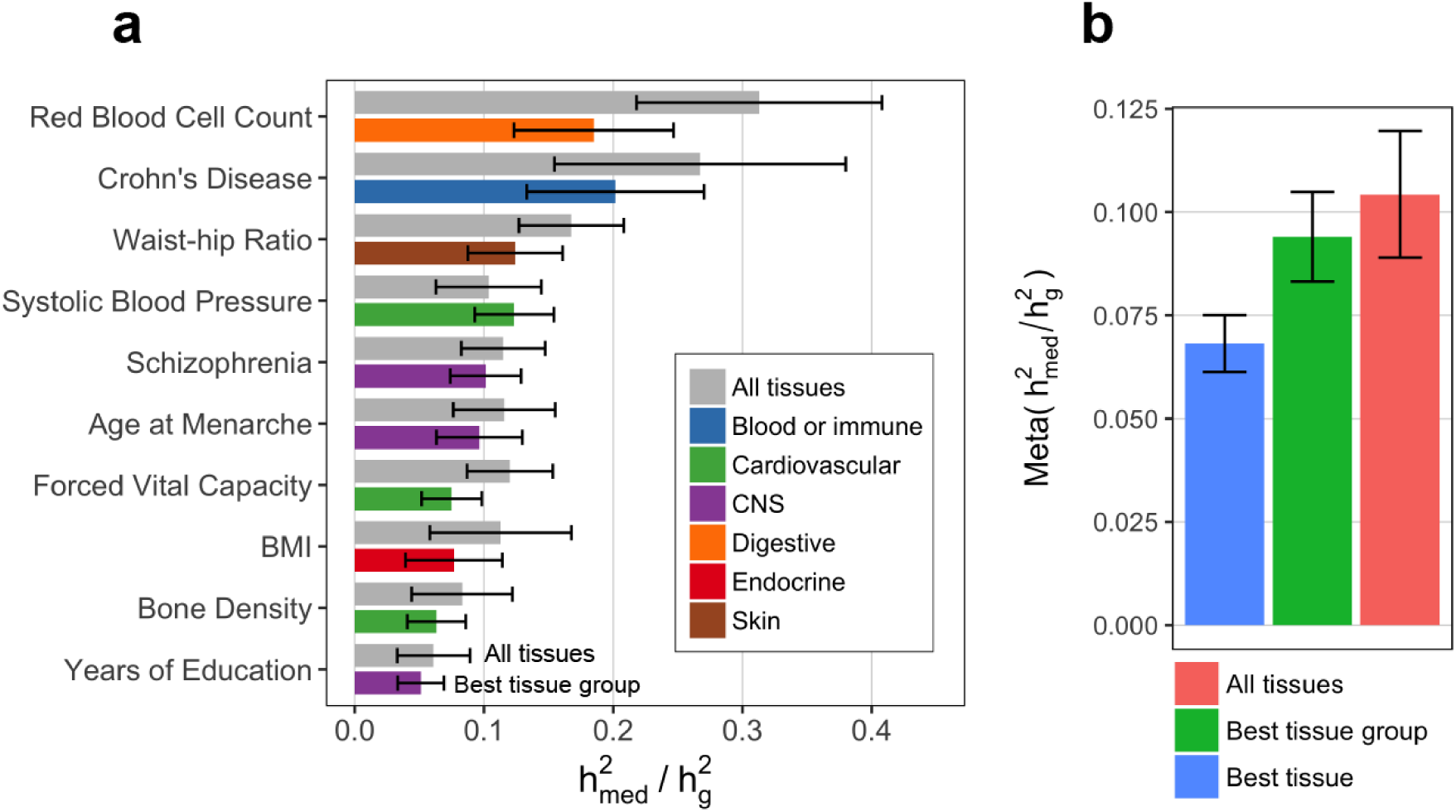
Estimates of proportion of heritability mediated by expression from GTEx. (**a**) Estimated proportion of heritability mediated by the cis-genetic component of assayed gene expression levels 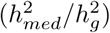 for 10 genetically uncorrelated traits. See Supplementary Note for procedure behind selecting these 10 traits and Supplementary Figure 12 for estimates of 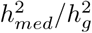 for all 42 traits. Error bars represent jackknife standard errors. For each trait, we report the 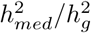 estimate for “All tissues” (expression scores meta-analyzed across all 48 GTEx tissues) and “Best tissue group” (expression scores meta-analyzed within 7 tissue groups). Here, “best” refers to the tissue group resulting in the highest estimates of 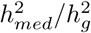 compared to all other tissue groups. (**b**) 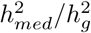 estimates meta-analyzed across all 42 traits. Here, “Best tissue” refers to the individual tissue resulting in the highest estimates of 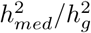 compared to all other tissues.

When we estimated 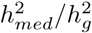 using tissue-group expression scores and all-tissue expression scores, we obtained consistently higher estimates of 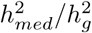 than individual-tissue expression scores (Figure 5b). This result suggests that we were able to account for low-*N*_*eQTL*_ dependent downward bias in 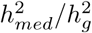 by meta-analysis of expression scores across tissues, consistent with our simulation result in Supplementary Figure We observed that the tissue-group resulting in the highest estimate of 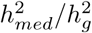 for each trait was mostly consistent with the known causal tissue of the trait (Figure 5a).

As independent validation, we used cis-eQTL summary statistics from eQTLGen^50^ (*N*_*eQTL*_ = 31,684 in blood only) to estimate 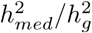 for the same 42 traits we analyzed above. We obtained very similar 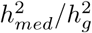 estimates as GTEx all-tissue expression for blood/immune traits and lower 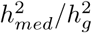 for non-blood/immune traits (Supplementary Figure 14, Supplementary Table 5), consistent with the fact that that eQTLGen only captures expression levels in blood while GTEx all-tissue meta-analysis captures expression levels across diverse tissues.

To explore the nature of dependencies between eQTL effect sizes, gene-trait effects, and non-mediated effects across the genome, we estimated 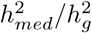 for all traits as before, but without stratifying genes and/or SNPs (Supplementary Figure 15). When we did not stratify genes by expression cis-heritability bins, we obtained much lower estimates of 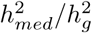 (mean estimate 0.011 with S.E. 0.003). This result is consistent with our simulation result in Figure 3a, which shows that not stratifying genes in the presence of a genome-wide negative correlation between the magnitude of eQTL effect sizes and gene-trait effect sizes leads to downwardly biased estimates of 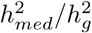. Moreover, when we did not stratify SNPs by the baselineLD model, we obtained much higher estimates of 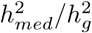 (mean estimate 0.45 with S.E 0.03). This result is consistent with our simulation result in Figure 3b, which shows that not partitioning SNPs in the presence of genome-wide positive correlation between the magnitude of eQTL effect sizes and non-mediated effect sizes leads to upwardly biased estimates of 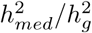. Together, these results demonstrate that both gene-eQTL independence and pleiotropy-eQTL independence are strongly violated in practice, justifying the necessity of stratifying genes by expression cis-heritability bins and stratifying SNPs by the baselineLD model to obtain unbiased estimates of 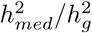.

Finally, we sought to investigate whether modifying the expression cis-heritability bins and baselineLD model influenced our 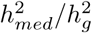 estimates. When we stratified genes in 10 bins rather than 5, we obtained very similar 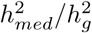 estimates (Supplementary Figure 16). Moreover, we observed that our 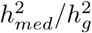 estimates were robust when making small changes to the baselineLD model (mean 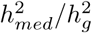 when individually removing each SNP category = 0.11) and quite robust to even large changes in the baselineLD model (mean 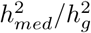 when removing 50% of all SNP categories = 0.13) (Supplementary Figure 17). These results demonstrate that our choice of stratifying genes by 5 expression cis-heritability bins and stratifying SNPs by the baselineLD model can robustly correct for biases in 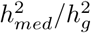 estimates.

### Genes with low expression heritability explain more expression-mediated disease heritability

To investigate the relationship between magnitude of expression cis-heritability 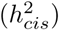 and amount of complex trait heritability mediated by those genes, we looked at the proportion of 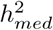 (defined as 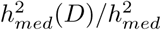 for gene category D) mediated by genes stratified into 10 bins by their 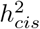 GTEx all-tissue meta-analyzed expression scores. Across 26 traits with 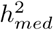. We performed all analyses using significantly greater than 0, we observed an inverse relationship between meta-tissue 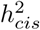 and proportion of 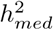 across gene bins (Figure 6a, Supplementary Table 6), with 32% of 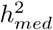 explained by the lowest 2 bins (mean meta-tissue 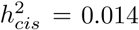 and only 3% of 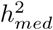 explained by the highest 2 bins (mean meta-tissue 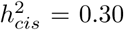. Because 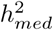 explained by a given gene is defined as the 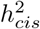 of the gene multiplied by its squared causal effect on the complex trait (in normalized units), the inverse relationship between 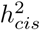 and 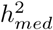 across gene bins implies that genes with less heritable expression (i.e. weaker/fewer eQTLs) have larger *causal effect sizes* on the complex trait (Figure 6b).

**Figure 6:**
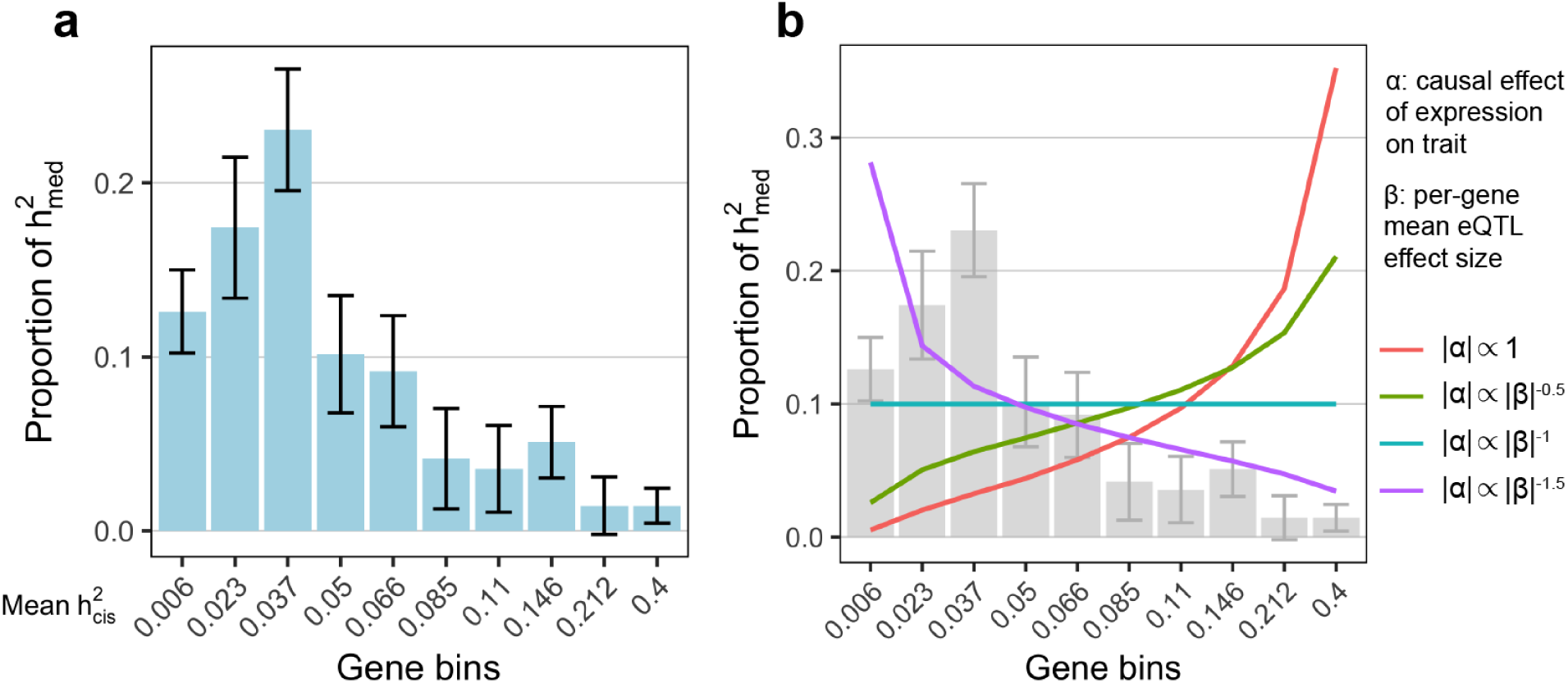
Low heritability genes explain more expression-mediated disease heritability. (**a**) Estimated proportion of expression-mediated heritability 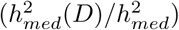 for 10 gene bins stratified by magnitude of expression cis-heritability. Results are meta-analyzed across 26 traits with nominally significant 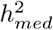. Results for individual traits can be found in Supplementary Table 6. (**b**) Expected proportion of 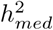 for various hypothetical relationships between |*α*| (causal effect magnitude of gene on disease) and |*β*| (mean per-gene eQTL effect size magnitude) across genes. |*α*| ∝ 1 implies that the magnitude of *α* is independent of the magnitude of *β*. |*α* | ∝ |*β*|^*p*^ with *p* < 0 implies an inverse relationship between the magnitude of *α* and *β* across genes, where *p* determines the strength of the inverse relationship. Empirical estimates of proportion of 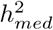 from **a** are replicated in the background.

There are several reasons why genes with less heritable expression might have larger causal effects on the complex trait. One explanation is that negative selection purifies out strong eQTLs for genes with large effect on complex traits^5, 35^. Alternatively, genes with low meta-tissue 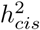 may consist of genes with tissue-specific eQTLs, which have been shown to be enriched for disease heritability^5, 19, 20^.

In support of the first explanation, we observed that the probability of being loss-of-function intolerant^51^ (i.e. pLI) and the level of selection against protein-truncating variants^52^ (i.e. *s*_*het*_) were both inversely correlated with meta-tissue 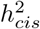 (Spearman’s *ρ* = −0.23 and −0.21 respectively) (Supplementary Figure 18). Moreover, consistent with the first explanation, we observed that the proportion of meta-tissue 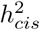 explained by the top eQTL was correlated with overall meta-tissue 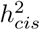 (*ρ* = 0.36; i.e. genes with low meta-tissue 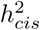 tended to be affected by multiple weaker eQTLs rather than a single strong eQTL) (Supplementary Figure 19). To investigate the second explanation, we considered the possibility that most low meta-tissue 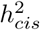 genes were primarily genes with nonzero individual-tissue 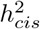 in only one or a small number of tissues, which is consistent with the genes having tissue-specific eQTLs. However, we did not observe strong evidence for this hypothesis (see “Role of tissue specificity in explaining low heritability genes” in Supplementary Note and Supplementary Figure 20). Moreover, we did not observe a strong relationship between meta-tissue 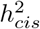 and the mean expression level of the gene across tissues (*ρ* = −0.02), nor the number of tissues in which the gene was expressed (*ρ* = 0.02) (Supplementary Figure 21).

In summary, our results support the hypothesis that negative selection contributes to the inverse relationship between expression cis-heritability and proportion of 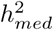.

### Expression-mediated heritability enrichment in functional gene sets

To gain insight into the distribution of expression-mediated effect sizes across various functional gene sets, we estimated 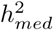 enrichment, defined as (proportion of 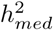) / (proportion of genes), for these gene sets. We gathered gene sets relevant for disease and specific biological pathways from publicly available sources (see Supplementary Table 7 for list of gene sets). We analyzed 827 gene sets from three main sources: (1) 10 gene sets reflecting various broad metrics of gene essentiality (Methods; URLs); (2) 780 gene sets reflecting specific biological pathways, including gene sets from the KEGG^53^, Reactome^54^, and Gene Ontology (GO)^55^ pathway databases (Methods; URLs); and (3) 37 gene sets composed of genes specifically expressed in 37 different GTEx tissues^56^ (Methods). We restricted our analyses to large gene sets with at least 200 genes (Methods), since we observed large standard errors in 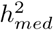 enrichment estimates for gene sets with 200 or fewer genes (Supplementary Figure 22). We applied MESC to estimate the 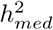 enrichment for each of the gene sets in 26 significant traits. As before, we stratified SNPs by the baselineLD model and genes by 5 cis-heritability bins. In addition to the baselineLD annotations, we considered including a SNP annotation corresponding to 100 Kb windows around each gene within each gene set, which enforces a more stringent criterion for identifying 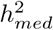 enrichment (Supplementary Note); however, we observed that including this SNP annotation had very little impact on our 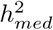 enrichment estimates (Supplementary Figure 23), so for simplicity we excluded it from our main analyses. We performed all analyses using GTEx all-tissue meta-analyzed expression scores (with the exception the tissue-specific expression analyses in Figure 7c, in which case we used GTEx tissue-group meta-analyzed expression scores).

**Figure 7:**
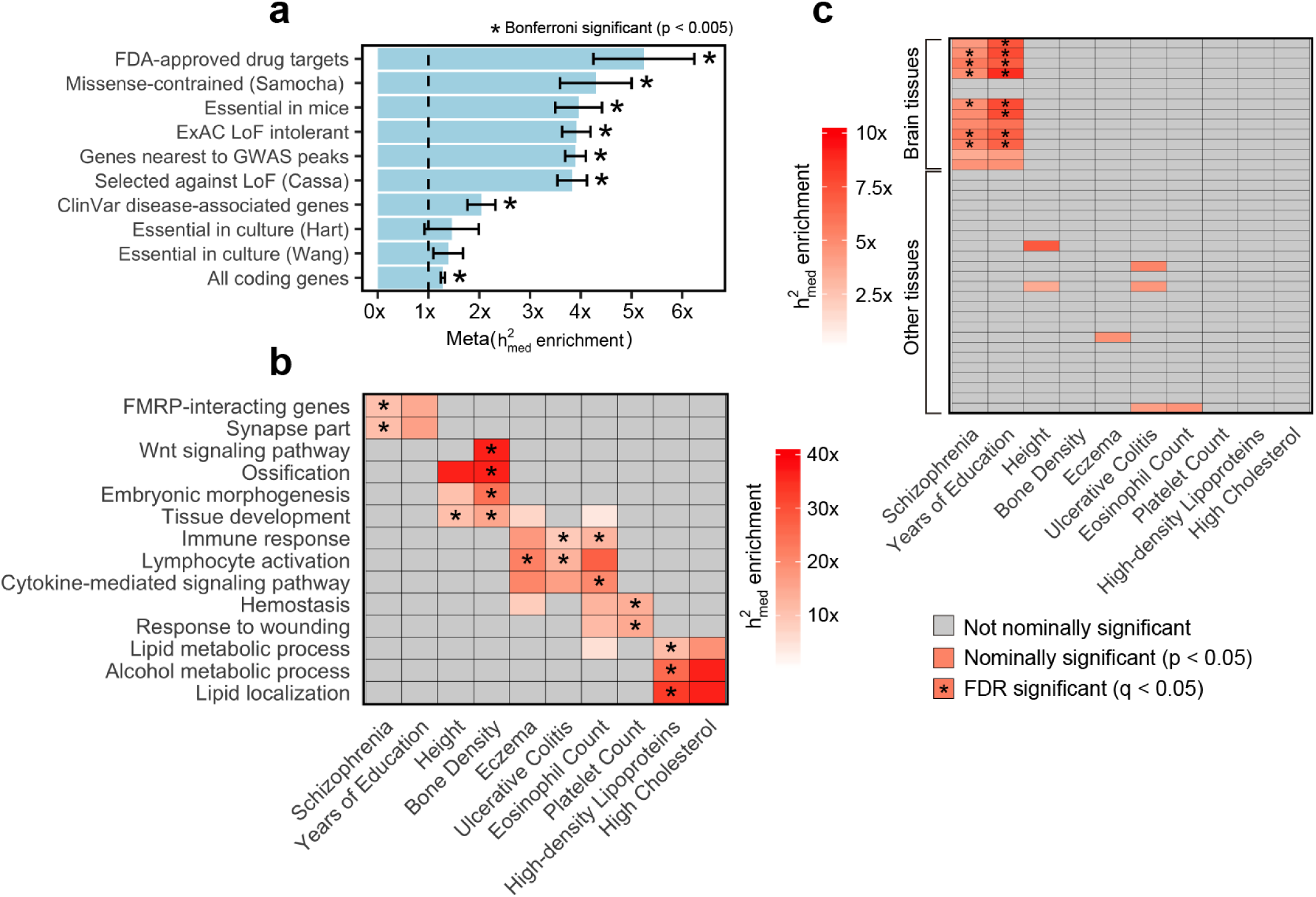
Expression-mediated heritability enrichment estimates for functional gene sets. For all plots, x axis represents complex traits and y axis represents gene sets. (**a**) 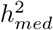 enrichment estimates for 10 broadly essential gene sets meta-analyzed across 26 complex traits. 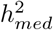 enrichment estimates for individual traits can be found in Supplementary Figure 24. (**b**) For ease of display, we report 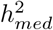 enrichment estimates for a representative set of 14 pathway-specific gene sets across 10 complex traits. 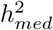 enrichment estimates for additional complex traits and gene sets can be found in Supplementary Figure 25 and Supplementary Table 8. (**c**) 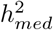 enrichment estimates for 37 gene sets corresponding to specifically expressed genes in 37 GTEx tissues. Brain tissues (13 total) are indicated as so in the figure. 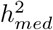 enrichment estimates for additional complex traits, with individual GTEx tissues labelled, can be found in Supplementary Figure 26.

Out of 21,502 gene set-complex trait pairs (827 gene sets × 26 complex traits), we observed 226 gene set-complex trait pairs (composed of 117 unique gene sets) with FDR-significant 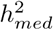 enrichment (FDR < 0.05 accounting for 21,502 hypotheses tested). Significant 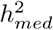 enrichment estimates ranged from 1.5x to 51x across gene-set complex trait pairs. The full list of 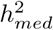 enrichment estimates for all 21,502 gene set-complex trait pairs is reported in Supplementary Table 8.

In Figure 7a, we show 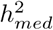 enrichment estimates for all 10 broadly essential gene sets meta-analyzed across 26 complex traits. 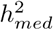 enrichment estimates for individual traits can be found in Supplementary Figure 24. We observed Bonferroni-significant meta-trait 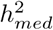 enrichment (p < 0.05 / 10) for 8 gene sets, including ExAC loss-of-function intolerant genes^51^ (3.9x enrichment; p = 2.3 × 10^−25^), FDA-approved drug targets^57^ (5.2x enrichment; p = 2.0 × 10^−5^), genes essential in mice^58–60^ (4.0x enrichment; p = 1.1 × 10^−10^), and genes nearest to GWAS peaks^61^ (3.9x enrichment; p = 5.0 × 10^−46^). When identifying significant meta-trait 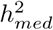 enrichment for broadly essential gene sets, we correct for only 10 hypotheses tested, since we compare broadly essential gene sets to only one another rather than the full set of 827 gene sets.

Of the 780 pathway gene sets, we observed that 97 had a significant 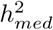 enrichment (FDR < 0.05) in at least one of the 26 complex traits. In Figure 7b, we show the 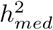 enrichment estimates of a representative set of 140 gene set-complex traits pairs, consisting of 14 pathway-specific gene sets × 10 traits. More comprehensive results, consisting of 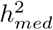 enrichment estimates for all 97 significant pathway gene sets × 26 complex traits, can be found in Supplementary Figure 25. Most gene sets exhibited highly trait-specific patterns of enrichment that were consistent with the known biology of the trait, including fragile X mental retardation protein (FMRP)-interacting genes for schizophrenia^62, 63^, Wnt signaling for bone density^64^, and hemostasis for platelet count^65^.

Finally, we investigated whether genes specifically expressed in 37 different GTEx tissues^56^ were enriched for 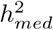. It has previously been shown that SNPs near the top 10% of genes most differentially expressed in putative causal tissues/cell types for specific traits are highly enriched for complex trait heritability^56^. However, it is unknown whether the effects of these SNPs are mediated by the expressed genes, or whether they impact the trait through alternate mechanisms in the same loci. We examined the same set of specifically expressed genes in GTEx tissues investigated in ref.^56^. We found significant 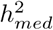 enrichment (FDR < 0.05) in genes specifically expressed in brain tissues for brain-related traits (schizophrenia and years of education) (Figure 7c), suggesting that the complex trait heritability of SNPs near genes specifically expressed in causal tissues (at least for brain traits) is mediated by the expression of those genes.

Given that MESC can be used to prioritize disease-relevant gene sets based on the magnitude of their 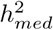 enrichment, it falls alongside a large class of methods that aim to perform gene set enrichment analysis from GWAS data^66–72^. We compared results from MESC to two other popular gene set enrichment methods applied to the same GWAS summary statistics we analyzed, MAGMA^69^ and DEPICT^68^ (Supplementary Note). We observed that around 1/3 of the gene set-complex trait pairs identified as having significant 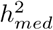 enrichment by MESC were not detected as significantly enriched by MAGMA or DEPICT, including biologically plausible gene set-complex trait pairs such as “phospholipid metabolic process” for high-density lipoprotein level and “synapse part” for schizophrenia (Supplementary Figure 27 and Supplementary Table 9). This result demonstrates that MESC captures unique signal from eQTL data that is missed by other methods.

## Discussion

We have developed a new method, mediated expression score regression (MESC), to estimate complex trait heritability mediated by the cis-genetic component of assayed expression levels 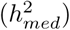 from GWAS summary statistics and eQTL effect sizes estimated from an external expression panel. Our method is distinct from existing methods that identify and quantify overlap between eQTLs and GWAS hits (including colocalization tests^6–10^, transcriptome-wide association studies^11–15^, and heritability partitioning by eQTL status^17–20^) in that it specifically aims to distinguish directional mediated effects from non-directional pleiotropic and linkage effects. Moreover, our polygenic approach does not require individual eQTLs or GWAS loci to be significant and is not impacted by the sparsity of eQTL effect sizes, so unlike other approaches^9–13, 19^ we do not exclude genes or SNPs from our analyses based on any significance thresholds. We applied our method to summary statistics for 42 traits and eQTL effect sizes estimated from 48 GTEx tissues. We show that across traits, a significant but modest proportion of complex trait heritability (on average 0.11 with S.E. 0.02) is mediated by the cis-genetic component of assayed expression levels. Though many previous approaches have hypothesized that SNPs impact complex traits by directly modulating gene expression levels, our results provide the first concrete genome-wide evidence for this hypothesis. On the other hand, the fact that our 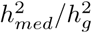 estimates are low for most traits suggests that eQTLs estimated from steady-state expression in bulk post-mortem tissues (e.g. from GTEx) do not capture most of the mediated effect of complex trait heritability, motivating additional assays to better identify molecular mechanisms impacted by regulatory GWAS variants.

There are two potential explanations for our low 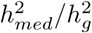 estimates. Recall that our estimates of 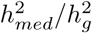 are a function of two quantities: (1) the proportion of heritability mediated by the cis-genetic component of expression levels *in the true causal cell types/contexts for the trait* 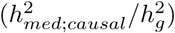 multiplied by (2) the average squared cis-genetic correlation between expression in *assayed tissues T* and expression in causal cell types/contexts for the trait 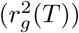). Thus, low 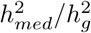 estimates may be due to one of the following explanations:

1. The proportion of complex trait heritability mediated by the cis-genetic component of gene expression levels is in fact high in causal cell types/contexts for the trait, but eQTL data from bulk assayed tissues (e.g. from GTEx) is a poor proxy for eQTL data in causal cell types/contexts, causing 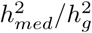 to be low. In other words, 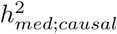 is high, while *r*^2^(*T*) is low.
2. The proportion of complex trait heritability mediated by the cis-genetic component of gene expression levels is low even in causal cell types/contexts for the trait. In particular, complex trait heritability may be mediated in ways other than through gene expression in cis, including mediation through protein-coding changes or splicing, or mediation through expression levels purely in trans. Note that our model *does* capture cis-by-trans effects, in which a SNP acts as a trans-eQTL for a gene via its effects in cis on the expression of another gene (Supplementary Note).

Both above explanations (with the first explanation perhaps more parsimonious than the second) imply that eQTLs measured in bulk post-mortem tissues (e.g. from GTEx) are inadequate to explain the majority of regulatory effects from GWAS variants. This motivates additional assays to more fully capture these regulatory effects. If explanation 1 is predominant, then context-specific expression^73^ and single-cell expression^74^ assays are desirable to capture expression levels in highly cell type/context specific scenarios. If explanation 2 is predominant, then additional assays such as splicing^75^, histone mark^4^, chromosome conformation^76^, and trans-eQTL^50^ assays can be informative for probing other molecular mechanisms impacted by GWAS variants. Specifically regarding trans-eQTLs, we provide a derivation which shows that much larger gene expression assays than currently available are necessary to estimate disease heritability mediated by gene expression in trans (Supplementary Note). We anticipate that the additional assays listed above will allow us to explain a greater proportion of mediated complex trait heritability, and that MESC will be used to benchmark the progress made by future QTL studies.

As an aside, the fact that our 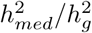 estimates are low may appear to be related to the fact that expression cis-heritability 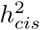 is also low, but we show that 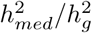 as estimated by MESC has no *a priori* relationship with the magnitude of 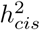 (Supplementary Note). In other words, the large amount of environmental/stochastic noise in gene expression measurements has no impact on our 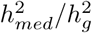 estimates, since 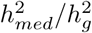 depends on only the *genetic component* of gene expression.

We observed that expression scores meta-analyzed across tissues gave us higher estimates of 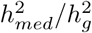 than individual-tissue expression scores, suggesting that disease-associated cis-eQTLs often affect expression in multiple tissues. This result is consistent with previous studies that reported higher heritability enrichment of cis-eQTLs meta-analyzed across all GTEx tissues compared to individual tissues^19^, higher prediction accuracy for imputed expression using joint prediction from multiple tissues compared to individual tissues^77^, and high cis-genetic correlations of expression between tissues overall^48, 78^. Nevertheless, we did obtain generally higher estimates of 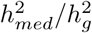 using expression scores from the presumed causal tissue group for a given disease compared to other tissue groups, suggesting some tissue-specific effects.

Despite the fact that assayed expression levels only explain a small proportion of overall disease heritability, we were still able to extract useful biological insight from partitioning expression-mediated heritability across gene categories. In particular, we observed a strong inverse relationship between proportion of 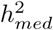 and expression cis-heritability across genes, suggesting that genes with low expression cis-heritability have large effects on complex traits. This result may be explained by negative selection purifying out strong eQTLs for genes with large effect on disease^35^, or by genetic compensatory mechanisms that buffer against the phenotypic effects of genes with strong genetic effects on their expression^79, 80^. Alternatively, genes with low expression cis-heritability may have highly cell type-specific eQTLs whose effect sizes are diluted by the inclusion of other cell types in a bulk assayed tissue sample. Regardless of the underlying nature of low-heritability genes, this result suggests that integrative association tests that prioritize genes based on probability of colocalization between eQTLs and GWAS hits^6, 8, 9^ and/or significance of genetic correlation between expression and trait^11–13^ may not detect the most mechanistically important genes, since these methods have lower power for genes with weaker eQTLs. Instead, our result suggests that genes with weaker eQTLs should be prioritized, and it motivates the implementation of larger eQTL studies and/or cell-type specific assays to more accurately detect these weak eQTLs.

We did not observe significantly nonzero 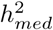 for low polygenicity traits (pigmentation traits and balding) and many neurological traits (autism, number children ever born, depressive symptoms, morning person, neuroticism, smoking status, anorexia). There is evidence that pigmentation traits are driven in large part by coding changes in MC1R^81^ and a small number of other genes^82–84^. Moreover, there is evidence that eQTLs important for neurological traits can be unique to fetal brain expression and thus highly context-specific^85, 86^. Neither the protein-coding changes in pigmentation genes nor the context-specific eQTLs in brain genes will be captured by steady-state expression level measurements in assayed postmortem tissues, and thus any SNP effects mediated by them will also not be observed in our analysis.

There are several limitations to our method. First, our method makes the assumption that the magnitude of eQTL effect sizes is uncorrelated with the magnitude of both gene-trait effect sizes (gene-eQTL independence) and non-mediated effect sizes (pleiotropy-eQTL independence) within each SNP/gene category included in the model. Violations to genome-wide gene-eQTL independence can be resolved by stratifying genes by the magnitude of their expression cis-heritability; given that expression cis-heritability is directly estimable for each gene in practice, violations to genome-wide gene-eQTL independence are readily resolvable (see Figure 3a and Supplementary Figure 16). Meanwhile, violations to genome-wide pleiotropy-eQTL independence can be resolved by stratifying SNPs by the magnitude of their eQTL effect sizes; however, due to LD, we cannot obtain precise estimates of eQTL effect sizes for individual SNPs. Instead, our model assumes that the SNP categories in the baselineLD model act as a reasonable proxy for eQTL effect size magnitude. If the magnitude of eQTL effect sizes and non-mediated effect sizes are positively correlated within the baselineLD categories included in our analyses, then our estimates of 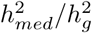 would be up-wardly biased. However, given that the baselineLD model was able to account for extreme violations to pleiotropy-eQTL independence in simulations (Figure 3b), and that the average 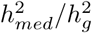 across traits was only 0.11, this upward bias (if present) is likely to be small. An alternative approach to estimating 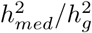 involves regressing pairwise products of GWAS summary statistics on pairwise products of eQTL summary statistics^87^. Unlike MESC, this approach yields unbiased estimates 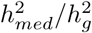 even when pleiotropy-eQTL independence is violated within SNP categories. However, the approach requires aggressive LD pruning in order to retain SNP pairs in linkage equilibrium, causing the approach to be highly underpowered in practice (L.J.O, unpublished data). The second limitation of our method is that it relies on the accurate estimation of expression scores from external expression panel samples. In order for our method to be well-powered, it requires large expression panel sample sizes that can only be obtained through meta-analysis across individual tissues at current sample sizes. Third, the quantity that our method estimates in practice (i.e. heritability mediated by *assayed* gene expression levels) can potentially be much smaller than the theoretical quantity of heritability mediated by expression levels in causal cell types/contexts if assayed gene expression levels do not adequately capture expression levels in causal cell types/contexts. Fourth, our method can only provide reliable 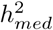 enrichment estimates for large gene sets on the order of 200 or more genes, so smaller gene pathways or individual genes cannot be prioritized using our method. Fifth, our method does not capture non-additive effects of SNPs on gene expression or gene expression on trait.

Despite these limitations, our method provides a novel framework to distinguish mediated effects from pleiotropic and linkage effects and will be useful for quantifying the improvement of new molecular QTL studies over existing assays in capturing regulatory disease mechanisms. Moreover, partitioning mediated heritability can provide insight into regulatory effects mediated by specific gene sets or pathways.

## Acknowledgments

We thank B. Pasaniuc, R. Ophoff, H. Shi, S. Groha, K. Siewert, S. Gazal, and A. Liu for helpful discussions. This research was funded by NIH grants T32HG002295, R01 MH115676, R01 MH107649, R01 MH101244, and U01 HG009379.

## Author contributions

D.W.Y, L.J.O, A.L.P, and A.G conceived of the project. D.W.Y and A.G. designed experiments. D.W.Y performed the experiments and analyzed the data. D.W.Y. and A.G. wrote the manuscript with input from L.J.O. and A.L.P.

## URLs

MESC software, https://github.com/douglasyao/mesc; 1000G Phase 3 data, ftp://ftp.1000genomes.ebi.ac.uk/vol1/ftp/release/20130502, baselineLD annotations, https://data.broadinstitute.org/alkesgroup/LDSCORE/; S-LDSC software, https://github.com/bulik/ldsc, GTEx v7 data, https://www.gtexportal.org/home/datasets; Macarthur lab gene sets, https://github.com/macarthur-lab/gene_lists; Molecular Signatures Database gene sets, http://software.broadinstitute.org/gsea/msigdb/collections.jsp

## Methods

### Definition of 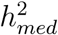

We model trait ***y*** for *N* individuals as follows:

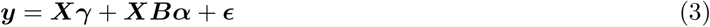

where ***y*** is an *N* -vector of phenotypes (standardized to mean 0 and variance 1), ***X*** is an *N* × *M* genotype matrix for *M* SNPs (standardized to mean 0 and variance 1), ***γ*** is an *M* -vector of non-mediated SNP effect sizes on the trait (including pleiotropic, linkage, and trans-eQTL-mediated effects), ***B*** is an *M* × *G* matrix of cis-eQTL effect sizes *in the causal cell types/contexts* for *G* genes, ***α*** is a *G*-vector of causal gene expression effect sizes on the trait, and ***ϵ*** is an *N* -vector of environmental effects. We treat all variables as random. We define 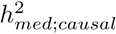 as follows:

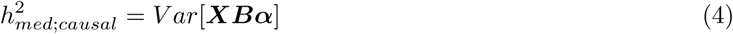

Under the assumption that *α* and *β* are independent of each other, we can rewrite this as follows:

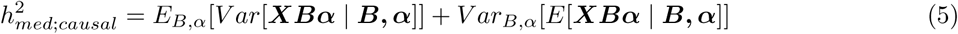

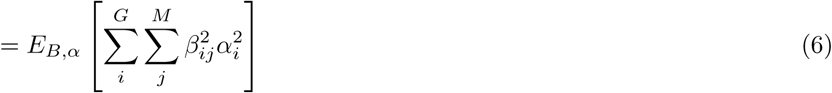

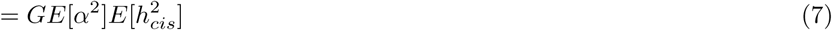

where *E*[*α*^2^] is the average squared per-gene effect of expression on trait and 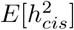 is the average cis-heritability of expression across all genes. (6) follows because *E*[***XBα*** | ***B, α***] = 0. We define 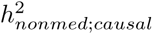 in a similar fashion:

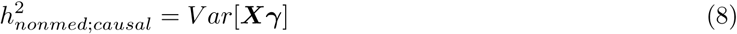

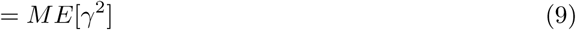

where *E*[*γ*^2^] is the average squared per-SNP effect on trait that is not mediated by gene expression.

In practice, expression levels in causal cell types/contexts for the complex trait are likely not assayed. Given a set of assayed tissues *T* (which may or may not be causal for the complex trait), we define 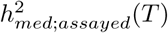 as follows:

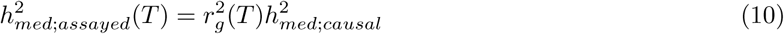

while we define 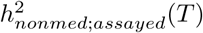 as 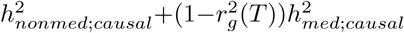. Here, 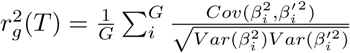 and denotes the average squared genetic correlation between expression in assayed tissues *T* vs. in causal cell types/contexts, where 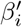 represents cis-eQTL effect sizes on gene *i* in *T*. Note that *β*′ can refer to either single tissue or meta-tissue cis-eQTL effect sizes, depending on whether *T* contains one or multiple tissues.

### MESC with SNP effect sizes known

For illustrative purposes, we walk through a derivation for MESC in the idealized scenario that we know (1) the true eQTL effect sizes, *β*, of each SNP on each gene (in the causal cell types/contexts for the trait) and (2) the true phenotypic effect sizes, *ω*, of each SNP on *y*. In practice, due to LD between SNPs, these quantities are not directly estimable.

Under the generative model (3), the total effect of SNP *k* on the complex trait is

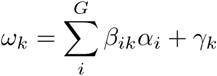

Given conditional independence of *α* and *γ* given *β*, upon squaring *ω*_*k*_ we have

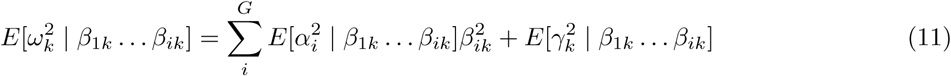

Assuming *unconditional* independence of *α* and *γ* (which requires that we make additional effect size independence assumptions involving *β*; see below), this simplifies to

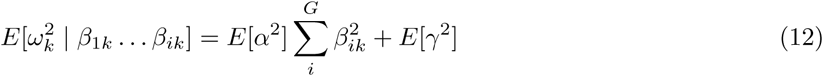

We use equation (12) to estimate *E*[*α*^2^] by regressing *ω*^2^ for all SNPs on 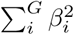 and taking the slope, while we estimate *E*[*γ*^2^] by taking the intercept. See Figure 1c for a plot illustrating this approach.

*E*[*α*^2^] can be multiplied by 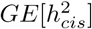 to obtain 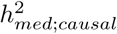, while *E*[*γ*^2^] can be multiplied by *M* to obtain 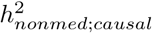. Note that violations to effect-size independence assumptions will lead to biased estimates of 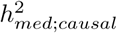 and 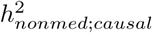, but can be remedied by stratifying SNPs and genes (see below).

### Model assumptions

In this section, we describe the two main effect size independence assumptions that are needed to derive equation (12), we discuss how they might be violated, and we show that conditioning on SNP- and gene-level annotations can ameliorate any resulting bias. These assumptions are:

- Across all genes, the magnitude of gene effect sizes is uncorrelated with the magnitude of eQTL effect sizes (i.e. *Cov*(*α*^2^, *β*^2^) = 0). We refer to this assumption as gene-eQTL effect size independence.
- Across all SNPs, the magnitude of non-mediated SNP effect sizes is uncorrelated with the magnitude of eQTL effect sizes (i.e. *Cov*(*γ*^2^, *β*^2^) = 0). We refer to this assumption as pleiotropy-eQTL effect size independence.

#### Violation of gene-eQTL effect size independence

Gene-eQTL effect size independence is violated in the scenario that less heritable genes have larger causal effect sizes on the trait, which is supported by evidence that evolutionarily constrained genes tend to have fewer eQTLs^5^. A negative correlation between the magnitude of gene effect sizes and eQTL effect sizes across the genome will result in downwardly biased estimates of *E*[*α*^2^] and upwardly biased estimates of *E*[*γ*^2^], as illustrated in Supplementary Figure 29a. The downward bias in 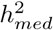 arises due to the fact that during regression, SNPs with larger eQTL effect sizes are implicitly weighted more than SNPs with smaller eQTL effect sizes, so the average slope will be biased toward the value of *α*^2^ for high heritability genes.

In order to account for violations to gene-eQTL effect size independence, we can stratify genes by the magnitude of their cis-heritability so that within each gene category, gene-eQTL effect size independence approximately holds. We can then obtain unbiased estimates of 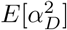 for each gene category *D* as illustrated in Supplementary Figure 29b. In practice, we stratify genes by 5 bins according to their cis-heritability, which we show adequately captures genome-wide dependence of gene effect sizes on eQTL effect sizes in simulations (Figure 3a).

#### Violation of pleiotropy-eQTL effect size independence

Pleiotropy-eQTL effect size independence is violated in the presence of regulatory hotspots with high biological activity in the genome^2, 36–39^, resulting in an increased number and/or magnitude of both eQTLs and pleiotropic/linkage effects in these hotspots. A positive correlation between the magnitude of non-mediated effect sizes and eQTL effect sizes across the genome will result in upwardly biased estimates of *E*[*α*^2^] and downwardly biased estimates of *E*[*γ*^2^], as illustrated in Supplementary Figure 30a.

In order to account for violations to pleiotropy-eQTL effect size independence, we can stratify SNPs by the magnitude of their eQTL effect sizes so that within each SNP category, pleiotropy-eQTL effect size independence approximately holds. We can then obtain unbiased estimates of overall *E*[*α*^2^] and 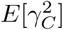 for each SNP category *C* as illustrated in Supplementary Figure 30b. Note that in practice, LD between SNPs makes it is difficult or impossible to identify the exact SNPs that act as eQTLs. This in turn makes it impractical to stratify SNPs according to eQTL effect size. Therefore, we instead stratify SNPs by a set of comprehensive functional SNP annotations, the baselineLD model^2, 43^, which should capture most known regulatory hotspots in the genome and act as a reasonable proxy to eQTL effect sizes (see Figure 3b for simulation results).

### MESC with eQTL effect sizes in non-causal tissues

In this section, we show that when we carry out the regression procedure described in “MESC with SNP effect sizes known” using eQTL effect sizes assayed in non-causal tissues *T*, we obtain an estimate of the quantity 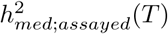, as defined above.

Let *β* represent cis-eQTL effect sizes in causal cell types/contexts for the trait, and *β*′ represent cis-eQTL effect sizes in assayed tissues *T*. We start with regression equation (12):

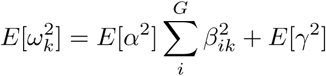

The ordinary least squares estimate of the coefficient from regressing *ω*^2^ on 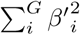 is

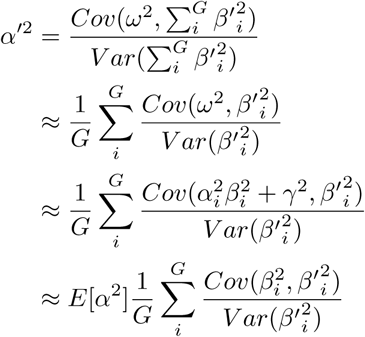

The third line follows given the *Cov*(*γ*^2^, *β*′^2^) = 0 and *Cov*(*α*^2^, *β*′^2^) = 0. Let 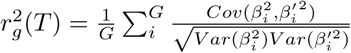 represent the average squared genetic correlation between expression in *T* vs. in causal cell types. Given this definition, we have

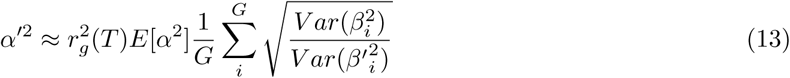

For simplicity, we make the assumption that 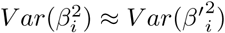 across genes. Note that violations to this assumption can realistically occur in practice but will not bias our estimate of 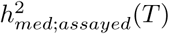 (Supplementary Note). Given this assumption, we have

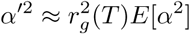

We can then multiply *α*′^2^ by 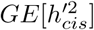 to obtain an unbiased estimate of 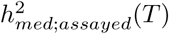, where 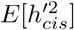 is the average expression cis-heritability of genes in *T*.

### MESC using summary statistics

We have previously derived an estimator for 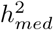 in the idealized scenario that SNP effect sizes are given. In practice, we use GWAS summary statistics, which are affected by sampling noise and by LD. It has previously been shown that LD and sampling noise can be accounted for by regressing GWAS *χ*^2^ statistics on LD scores, which measure the total of LD for each SNP^2, 34^. Under our generative model (3), the marginal OLS estimate of the total effect size of a SNP *k* on the trait is given by

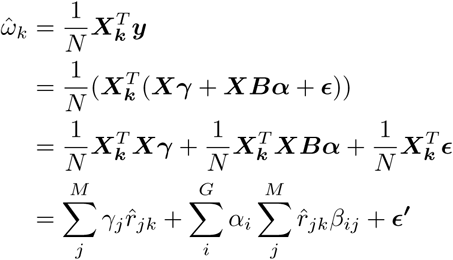

Let 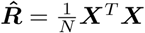 denote the in-sample LD matrix, and let 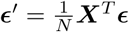 denote the noise term in the summary statistics. The *χ*^2^ statistic for SNP *k* (defined as 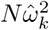) is:

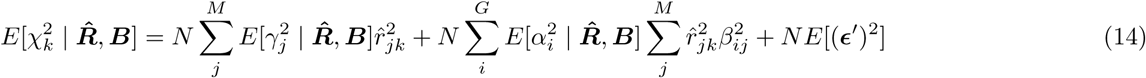

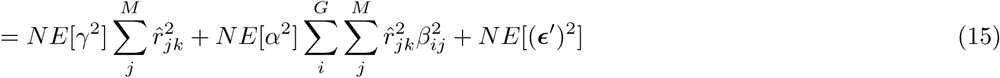

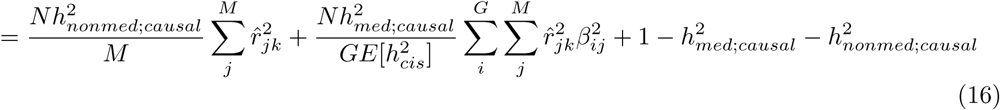

In order for the equation regarding the unconditional expectation of 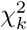 to hold true (15), we must make two independence assumptions involving LD-dependent genetic architecture, in addition to the independence assumptions described in “Model assumptions” (see above). LD-dependent architecture, if not accounted for, is known to produce bias in heritability estimates^88, 89^. These assumptions are:

- Across all genes (indexed by *i*), the magnitude of *α*_*i*_ is uncorrelated with the LD scores of eQTLs for gene *i*
- Across all SNPs (indexed by *k*), the magnitude of *γ*_*k*_ is uncorrelated with the LD score of SNP *k* (16) follows (15) from our definitions of 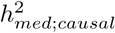 and 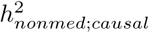. Since 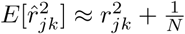, we have

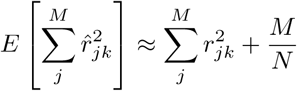

and

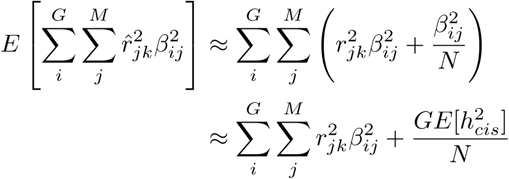

Thus,

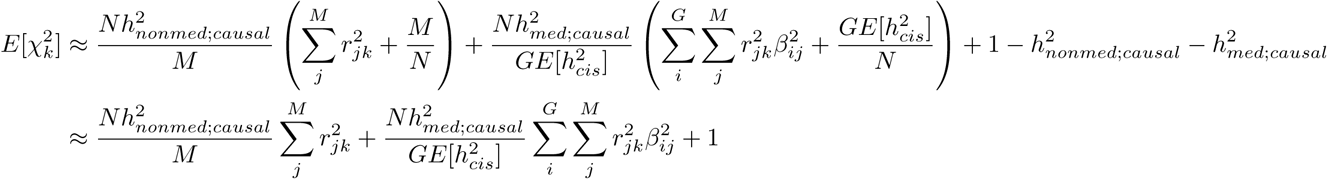

Defining LD scores 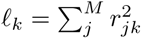 and expression scores 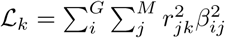, we arrive at our main equation for summary MESC regression:

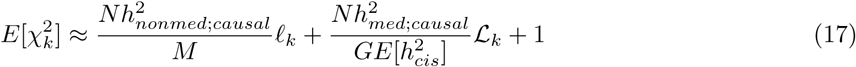

Analogous to the derivation in “MESC with eQTL effect sizes assayed in non-causal tissues” (see above), we show that if we perform this regression using expression scores in assayed tissues *T* rather than expression scores in causal cell types/contexts, we will estimate 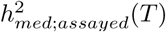 rather than 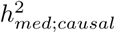 (see “MESC with assayed expression scores” in Supplementary Note).

### Stratified MESC

In this section, we extend summary-based MESC to estimate 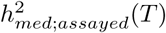 partitioned over groups of genes. Note that stratified MESC can be viewed as a special form of stratified LD score regression^2^ (Supplementary Note). Given *D* potentially overlapping gene categories 𝒟_1_, …, 𝒟_*D*_, we define 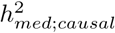 partitioned over gene categories as follows:

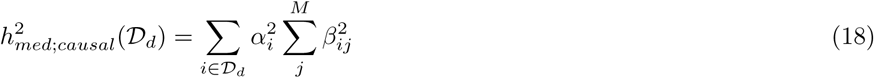

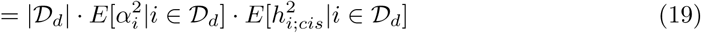

where 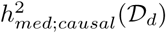 is the heritability mediated in cis through the expression of genes in category 𝒟_*d*_, |𝒟_*d*_| is the number of genes in 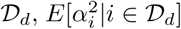 is the average squared causal effect of expression on trait for genes in 𝒟_*d*_, and 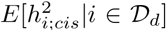 is the average cis-heritability of expression of genes in 𝒟_*d*_. Similar to equation (12), equation (19) relies on an independence assumption between *α* and *β*, namely that *α*_*i*_ ⊥ *β*_*i*_ | *i* ∈ 𝒟_*d*_.

For gene *i*, we model the variance of gene effect size *α*_*i*_ as

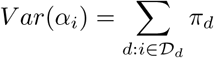

Assuming that gene categories 𝒟_*d*_ form a disjoint partition of the set of all genes, we have

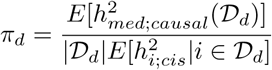

If gene categories are overlapping, then *π*_*d*_ can be conceptualized as the contribution of annotation 𝒟_*d*_ to 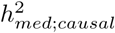 conditional on contributions from all other gene categories included in the model.

Given *C* potentially overlapping SNP categories 𝒞_1_, …, 𝒞_*C*_, we define 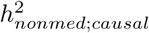 partitioned over SNP categories as follows:

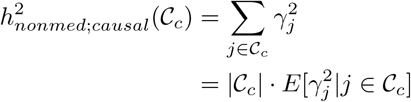

where 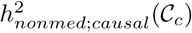 is the non-mediated heritability of SNPs in category 𝒞_*c*_, | 𝒞_*c*_| is the number of SNPs in 𝒞_*c*_, and 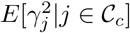 is the average squared non-mediated effect size of SNPs in 𝒞_*c*_.

For SNP *j*, we model the variance of non-mediated effect size *γ*_*j*_ as follows:

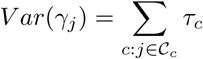

Assuming that SNP categories 𝒞_*c*_ form a disjoint partition of the set of all SNPs, we have

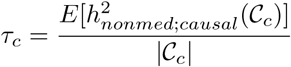

If SNP categories are overlapping, then *τ*_*c*_ can be conceptualized of as the contribution of annotation 𝒞_*c*_ to 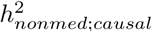 conditional on contributions from all other SNP categories included in the model.

The equation for stratified MESC is

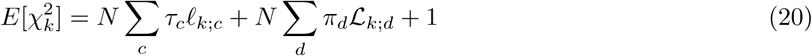

where 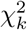 is the GWAS *χ*^2^-statistic of SNP *k, N* is the number of samples, *ℓ*_*k*;*c*_ is the LD score of SNP *k* with respect to SNP category 𝒞_*c*_ (defined as 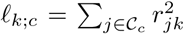), and ℒ _*k*;*d*_ is the expression score of SNP *k* with respect to gene category 𝒟_*d*_ (defined as 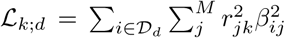). See Supplementary Note for a derivation of this equation. Analogous to unstratified MESC, when we perform this regression using expression scores in assayed tissues *T* rather than expression scores in causal cell types/contexts, we will estimate 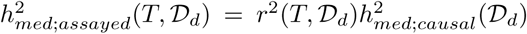, where *r*^2^(*T*, 𝒟_*d*_) is the average squared genetic correlation of expression between *T* and causal cell types/contexts for genes in 𝒟_*d*_.

### Estimation of expression scores

In order to carry out the regression described in equation (20), we must first estimate expression scores ℒ_*k*;*d*_ (where 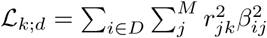) from an external expression panel. ℒ_*k*;*d*_ can be conceptualized as the total expression cis-heritability of genes in *D* that is tagged by SNP *k*, and is equivalent in expectation to squared eQTL summary statistics for SNP *k* summed across genes in *D* 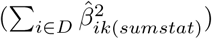 modulo a constant error term that arises from squaring the sampling noise in the eQTL summary statistics. Thus, we can in theory use eQTL summary statistics to compute ℒ_*k*;*d*_ and estimate 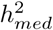 (see Supplementary Note).

However, because gene expression tends to have low cis-heritability across genes^5^ and most individual tissue expression panels have small sample sizes (less than 500 samples per GTEx tissue), expression scores estimated using eQTL summary statistics can be very noisy and can lead to downwardly biased estimates of 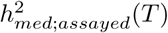 due to regression attenuation^90^ (Supplementary Figure 1). Therefore, we estimate individual-tissue expression scores as follows in order to reduce sampling noise. Assuming that individual level genotypes for expression panel samples are available, we leverage prior knowledge regarding the sparse nature of eQTL effect sizes and use LASSO^41^ to obtain regularized estimates of causal eQTL effect sizes 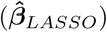, then multiply 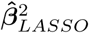 by the element-wise squared LD matrix ***R***^2^ to obtain 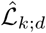:

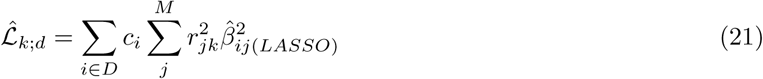

Here, *c*_*i*_ refers to a correction factor we apply to 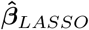 to both reduce bias and improve prediction accuracy compared to uncorrected estimates. In particular, we observed that LASSO without correction produces downwardly-biased estimates of eQTL effect sizes in simulations (Supplementary Figure 2), leading to upwardly-biased estimates of 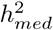 (Supplementary Figure 1). We calculate *c*_*i*_ as follows. First, we use REML to obtain an unbiased estimate of 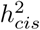for each gene *i* 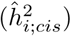, which in expectation is equivalent to 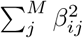. For each gene, we then scale 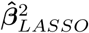 by a factor *c*_*i*_ so that 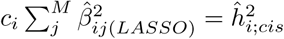. We can obtain approximately unbiased estimates of the squared LD between two SNPs using the formula 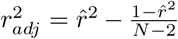, where 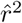 denotes the standard biased estimator of *r*^2^.

Other methods to estimate expression scores were also explored (Supplementary Note); we observed that the approach described in this section provided the best performance in simulations (Supplementary Figures 1,2).

### Meta-analysis of expression scores

Given our method of computing expression scores in individual tissues outlined above, we meta-analyze expression scores across tissues as follows. We first obtain meta-tissue expression cis-heritability 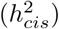 estimates for each gene by averaging individual-tissue 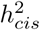 estimates across tissues. We scale individual-tissue LASSO-predicted causal eQTL effect sizes to the meta-tissue 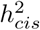 (see above), then average the scaled causal eQTL effect sizes across tissues. Finally, we multiply the averaged causal eQTL effect sizes by the element-wise squared LD matrix to obtain expression scores. In simulations, we show that this method of meta-analyzing expression scores produces nearly unbiased estimates of 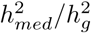 at 5 tissues × 200 samples per tissue (Supplementary Figure 3), which is comparable to the number expression panel samples in given tissue group (Supplementary Table 2).

There are several reasons why meta-analyzing expression scores across tissues is desirable. We would like to select a set of tissues *T* in which eQTL effect sizes are as close as possible to those in causal cell types/contexts for the complex trait, which would maximize 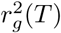. 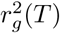 in practice refers to the squared correlation between *estimated* eQTL effect sizes in assayed tissues vs. true eQTL effect sizes in causal cell types/contexts, so sampling noise in eQTL effect size estimates will decrease 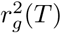 (see Supplementary Note). In simulations, we showed that 200 expression panel samples (comparable to the size of individual-tissue expression panels from GTEx) was inadequate to obtain unbiased estimates of 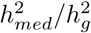 (Figure 2a, Supplementary Figures 1,2). Thus, meta-analysis of expression scores across tissues is necessary in order to eliminate downward bias in 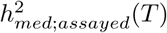 due to sampling noise in expression scores (Supplementary Figure 3). Moreover, agnostic of expression panel sample size, if meta-tissue eQTLs better tag eQTLs in causal cell types/contexts for the trait than individual tissue eQTLs, we will also increase 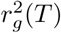 by meta-analyzing expression scores across tissues.

### Simulations

All simulations were conducted using genotypes from UK Biobank^40^ restricted to HapMap 3 SNPs^49^ on chromosome 1 (*M* = 98,499 SNPs). All simulations followed the same overall procedure, though with varying parameters; see below for specific parameters used in each individual simulation. The overall procedure is outlined here in chronological order:

1. *Simulation of expression data.* We simulated eQTL effect sizes for *G* = 1,000 genes, with eQTLs for each gene randomly selected in a 1 Mb window. We then simulated expression phenotypes for 1,000 expression panel samples (randomly selected from UK Biobank) using an additive generative model with appropriate environmental noise added, representing an expression panel. The exceptions were Figure 2a and Supplementary Figure 1, in which we varied the number of expression panel samples from 100 to 1,000, and Supplementary Figure 3, which had 200 samples per tissue.
2. *Simulation of GWAS data.* We simulated non-mediated SNP effect sizes and gene-trait effect sizes for all SNPs and genes corresponding to various levels of 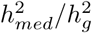. Total 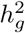 was fixed at 0.5 for all simulations (other than for Supplementary Figure 4, in which we varied 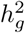). Together with the eQTL effect sizes simulated in the previous step, we used these effect sizes to simulate trait phenotypes using an additive generative model with appropriate environmental noise added for 10,000 GWAS samples (randomly selected from UK Biobank and distinct from the expression panel samples). We then produced GWAS summary statistics from this simulated data set using ordinary least squares.
3. *Estimation of expression scores.* We estimated expression scores from the expression panel samples using LASSO with REML correction (see “Estimation of expression scores” for description). For computational ease, we did not actually use REML to predict expression cis-heritability for each gene in each simulation; we instead took the true expression cis-heritability of the gene and added noise drawn from *N* (0, 0.01^2^) to simulate REML prediction error, which is consistent with empirical standard error estimates produced by GCTA (Supplementary Figure 32). This procedure is reasonable given that GCTA is unbiased, which we evaluated in sparse genetic architectures consisting of a single large effect and many small ones that mimics the cis-genetic architecture of gene expression (Supplementary Figure 33). In addition, although REML prediction error appears to be heteroskedastic, we obtained virtually identical estimates of 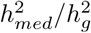 when modeling the standard error of 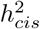 according to the best quadratic fit line between 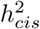 and 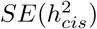 (Supplementary Figure 34), demonstrating that simulating REML error from *N* (0, 0.01^2^) was adequate for our simulation.
4. *Estimation of* 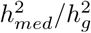. We estimated 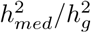 using MESC with the previously estimated expression scores, in-sample LD scores (computed from the 10,000 GWAS samples), and GWAS summary statistics.

The main parameter being changed across simulations was the distribution of eQTL effect sizes, non-mediated SNP effect sizes, and gene-trait effect sizes across the genome (including various levels of sparsity and correlations between effect sizes).

**Figure 2a: Impact of expression panel sample size on** 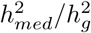 **estimates**

For this simulation, we varied 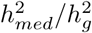 from 0 to 1. For each gene, we selected 5 SNPs to act as cis-eQTLs with locations randomly selected within a random 1 Mb window. The average cis-heritability across all genes was set at 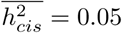. One eQTL for each gene was randomly selected to explain 80% of the total cis-heritability of the gene and had an effect size drawn from 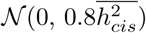. The remaining eQTLs had effect sizes drawn from 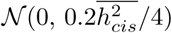. Expression phenotypes were simulated for each gene with environmental noise drawn from 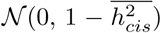. Non-mediated effect sizes were simulated for each SNP from 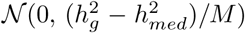. Gene-trait effect sizes simulated for each gene from 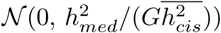). Complex trait phenotypes were simulated with environmental noise drawn from 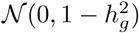.

**Figure 2b: Impact of sparse genetic/eQTL architectures on** 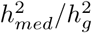 **estimates**

We set 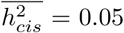. For the simulation involving 1 eQTL per gene, we drew its effect size from 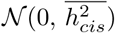. For the simulation involving 10% of genes being causal, we simulated gene-trait effect sizes from a point-normal distribution, with nonzero effects drawn from 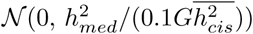. For the simulation involving 10% of SNPs being causal, we simulation non-mediated SNP effect sizes from a point-normal distribution, with nonzero effects drawn from 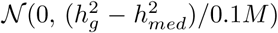. The remainder of simulation parameters were the same as in Figure 2a.

**Figure 2c:** 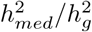 **estimates with** 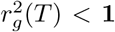

We set 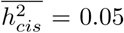. We simulated one eQTL per gene, with its effect size in the assayed tissue and causal tissue drawn from a multivariate normal distribution with mean **0** and 2 × 2 covariance matrix Σ with diagonal values 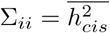 and off-diagonal values 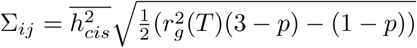, where *p* represents the proportion of SNPs with non-zero eQTL effect sizes on any gene (i.e. 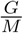). Simulating eQTL effect sizes in this fashion results in an average squared genetic correlation of 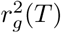 across all genes. We simulated non-mediated effect sizes, expression phenotypes for the assayed tissues, and complex trait phenotypes in the same manner as in Figure 2a. eQTL effect sizes used to generate complex trait phenotypes were taken in the causal tissue. Expression scores were estimated from only the assayed expression phenotypes.

**Figure 3a: Impact of violations to gene-eQTL independence on** 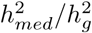 **estimates**

We simulated 1 eQTL for each gene with effect size drawn from a normal distribution with mean 0 and variance 100 · 2^*k*/200^/(Σ_*k*_ 2^*k*/200^, where *k* randomly indexes the genes from 1 to 1000. Simulating eQTL effect sizes in this fashion results in a realistic continuous distribution of eQTL effect sizes, where the quintiles for expression cis-heritability across genes are 0.016, 0.032, 0.064, 0.13, and 0.26. Next, we simulated gene-trait effect sizes from a normal distribution with mean 0 and variance 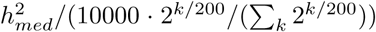.This causes the magnitude of gene-trait effects to be strongly inversely correlated with the magnitude of eQTL effect sizes across genes, but the per-gene 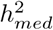 remains constant. We simulated non-mediated effect sizes, expression phenotypes, and complex trait phenotypes in the same manner as in Figure 2a.

**Figure 3b: Impact of violations to pleiotropy-eQTL independence on** 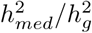 **estimates**

We set 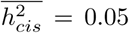. We simulated 1 eQTL per gene with effect size drawn from 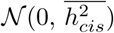. All eQTLs were selected to fall in coding regions. Next, for all SNPs in coding regions, we simulated non-mediated SNP effect sizes from 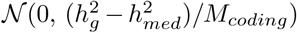, where *M*_*coding*_ is the number of SNPs that fall into coding regions. We simulated gene-trait effect sizes, expression phenotypes, and complex trait phenotypes in the same manner as in Fig 2a.

**Figure 4: Comparison of MESC to heritability partitioning**

We set 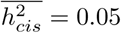. We simulated 1 eQTL per gene with effect size drawn from 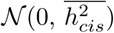. To simulate 10x enrichment of eQTLs in coding, TSS, and conserved regions, we selected eQTL locations so that 10x more eQTLs per SNP were located in the three SNP categories than the remainder of the genome. We simulated non-mediated effect sizes so that the heritability enrichment of the three SNP categories was 2.5x, 5x, or 10x. We simulated expression phenotypes and complex trait phenotypes in the same manner as Figure 2a. For stratified LD-score regression, we defined the eQTL category as the set of all true eQTLs. For MESC, we stratified SNPs by the baselineLD model with the three SNP categories and windows around the SNP categories removed from the model.

### Data and quality control

#### Genotypes

For MESC, we used European samples in 1000G^91^ as reference SNPs to compute LD scores. Regression SNPs were obtained from HapMap 3^49^. SNPs with GWAS *χ*^2^ statistics > max{80, 0.001*N*} (where *N* is the number of GWAS samples) and in the major histocompatibility complex (MHC) region were excluded. See Supplementary Note of ref.^2^ for justification of these procedures.

For computing expression scores, we downloaded genotypes derived from sequencing data for GTEx v7 from the GTEx Portal (URLs) as described in ref.^5^. We retained SNPs that were from HapMap 3^49^.

#### Expression data

We obtained processed and quantile normalized gene expression data for GTEx v7 from the GTEx Portal (URLs) as described in ref.^5^. For each tissue, the following covariates were included in all analyses: 3 genetic principal components, sex, platform, and 14-35 expression factors^92^ as selected by the main GTEx analysis.

### Estimation of expression scores from GTEx data

We used REML as implemented in GCTA^33^ to estimate the expression cis-heritability for each gene in each individual GTEx tissue. We then used LASSO as implemented in PLINK^93^ (with the LASSO tuning parameter set as the estimated expression cis-heritability of the gene) to estimate eQTL effect sizes for each gene in each individual GTEx tissue. In all procedures, we excluded gene-tissue pairs for which LASSO did not converge when predicting effect sizes. For Figure 5a and Supplementary Figure 12, we obtained causal eQTL effect size estimates in three different ways:

- *Meta-analysis across all tissues.* For each gene, we averaged the expression cis-heritability estimates across all 48 tissues. Within each tissue, we scaled the LASSO-predicted eQTL effect sizes to the averaged cis-heritability value. We then averaged the scaled eQTL effect sizes for each gene across all tissues. Genes were retained if they had a LASSO-converged eQTL effect size in at least one tissue.
- *Meta-analysis in tissue groups.* Of the 48 tissues, we grouped together 37 of them into 7 broad tissue groups: adipose, blood/immune, cardiovascular, CNS, digestive, endocrine, and skin (Supplementary Table 2). Within each tissue group, we averaged the expression cis-heritability estimates for each gene and scaled the LASSO-predicted eQTL effect sizes to the averaged cis-heritability value. We then averaged the scaled eQTL effect sizes for each gene across the tissues for each tissue group. Genes were retained in each tissue group if they had a LASSO-converged eQTL effect size in at least one tissue within that tissue group.
- *Individual tissues.* For each individual tissue, we scaled the LASSO-predicted eQTL effect sizes to the within-tissue-group averaged cis-heritability estimates.

The final eQTL effect sizes were then multiplied by the element-wise squared LD matrix in order to obtain expression scores (see “Estimation of expression scores”).

All remaining results in the manuscript, including Figures 6, 7a,b and Supplementary Figures 13, 15, 16, 17, 22, 23, 24, 25, 27, were obtained from expression scores meta-analyzed across all 48 GTEx tissues. The only exceptions were the tissue-specific expression analyses in Figure 7c and Supplementary Figure 26, in which we meta-analyzed expression scores within the tissue group of the focal tissue.

### Set of 42 independent traits

Analogous to previous studies^19, 94^, we initially considered a set of 34 traits from publicly available sources and 55 traits from UK Biobank for which GWAS summary statistics had been computed using BOLT-LMM v2.3^95, 96^. We restricted our analysis to 47 traits with z-scores of total SNP heritability above 6 (computed using stratified LD-score regression). The 47 traits included 5 traits that were duplicated across two datasets (genetic correlation of at least 0.9). For duplicated traits, we retained the data set with the larger sample size, leaving us with a total of 42 independent traits. When meta-analyzing estimates across traits, we performed random effects meta-analysis using the R package rmeta.

### BaselineLD categories

In all our analyses, we stratified SNPs by 72 functional categories specified by the baselineLD model v2.0^2, 43^ (URLs). These annotations include coding, conserved, regulatory (e.g., promoter, enhancer, histone marks, transcription factor binding sites), and LD-related annotations. The original baselineLD model v2.0 contains 76 total categories; we removed 4 categories corresponding to QTL MaxCPP annotations^19^ because the information contained in these annotations is redundant with the eQTL effect size information contained in expression scores.

#### Gene set analyses

In order for us to obtain to unbiased estimates of 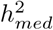 enrichment for the gene sets in our analysis, we must ensure that the gene-eQTL effect size independence assumption holds within each gene set (see “Model assumptions” above). Thus, in order to capture potential correlations between the magnitude of eQTL effect sizes and gene-trait effect sizes within gene sets, we partitioned each gene set into three separate bins based on the magnitude of their expression cis-heritability relative to other genes in the gene set. We then estimated 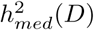 for each individual bin and aggregated these values together to estimate the overall 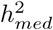 enrichment of the gene set.

#### Broad gene sets

We obtained gene sets corresponding to all coding genes, genes near significant GWAS hits in the NHGRI GWAS catalog^61^, genes essential in mice^58–60^, genes essential in cultured cell lines^97^, genes with any disease association in ClinVar^98^, and genes that are FDA-approved drug targets^57^ from the Macarthur lab GitHub page (URLs). We obtained an additional gene set for genes essential in cell lines^99^, genes depleted for protein-truncating mutations^51, 52^, and genes depleted for missense mutations^100^ from the supplementary data of the respective papers.

#### Pathway gene sets

We initially considered a set of 7,246 gene sets from the “canonical pathways” and “GO gene sets” collections from the Molecular Signatures Database^101^ (URLs), consisting of gene sets from BioCarta, Reactome, KEGG, GO, PID, and other sources. We restricted our analysis to 780 gene sets for which the number of genes with LASSO estimates of eQTL effect sizes that converged in individual GTEx tissues was at least 100 when averaged across all individual tissues. (Note that this roughly corresponds to gene sets with greater than 200 total genes; see Supplementary Table 7).

#### Tissue specific expression gene sets

We initially considered the full set of 48 GTEx tissues. We restricted our analysis to 37 gene sets for which the focal tissue belonged to one of the 7 main tissue groups we defined in our previous analyses (Supplementary Table 2). From ref.^56^, we obtained the set of 10% most specifically expressed genes in each of the 37 tissues.

