## Supplementary Note and Figures for "Quantifying genetic effects on disease mediated by assayed gene expression levels"

##### Modes of expression causality

In Supplementary Figure 28, we depict 9 different causality scenarios between SNPs, gene expression levels, and complex trait. Scenarios A-D constitute the main causality scenarios described in the main manuscript text; scenarios E-I describe additional causality scenarios. We provide a description of each scenario and its contribution to estimates of $h^2_{med}$.

###### A. Mediation

Here, the SNP affects the expression levels of the gene in cis, which then affect the complex trait. This is the desired scenario, since it is consistent with the hypothesis that SNPs exert their effects on complex traits via modulating gene expression levels. The presence of mediation will result in nonzero estimates of $h^2_{med}$.

###### B. Pleiotropy

Here, the SNP independently affects the expression of the gene in cis and the complex trait. Under the assumption that the magnitude of pleiotropic effects is uncorrelated with the magnitude of eQTL effects (see "Model assumptions"), the presence of pleiotropy will not contribute to estimates of $h^2_{med}$.

###### C. Linkage

Here, the SNP that affects gene expression in cis is in LD with another SNP that independently affects the trait. Under the assumption that the magnitude of linkage effects is uncorrelated with the magnitude of eQTL effects (see "Model assumptions"), the presence of linkage will not contribute to estimates of $h^2_{med}$.

###### D. Reverse mediation

Here, the SNP directly affects the complex trait, which then affects the expression levels of the gene in cis. Although this scenario can in theory contribute to nonzero estimates of $h^2_{med}$, the contribution will be negligible given that genetic effects on a complex (i.e. polygenic) trait are much smaller than genetic effects on gene expression (see "Reverse mediation").

###### E. Mediation in unobserved cell type/context

Here, the SNP affects the expression levels of the gene in the *causal cell type/context* for the complex trait, which then affects the complex trait. In practice, we only have access to *assayed* expression levels. Estimates of $h^2_{med}$ using assayed expression levels will be nonzero if the assayed expression levels are correlated with expression levels in causal cell types/contexts (see "Assayed vs. total underlying $h^2_{med}$").

###### F. Trans mediation

Here, the SNP affects the expression levels of the gene in *trans*, which then affects the complex trait. If trans-eQTL effect sizes are uncorrelated with cis-eQTL effect sizes, this scenario will not contribute to estimates of $h^2_{med}$. Note that this scenario refers to *purely trans* effects that are not mediated in cis at any point. An alternative scenario is cis-by-trans mediation (see below), which is subsumed by scenario A.

##### G. Cis-by-trans mediation

Here, the SNP affects the expression levels of gene 1 in cis, gene 1 affects gene 2 in trans, and gene 2 affects the complex trait. Although the SNP acts as a trans-eQTL for gene 2, its effects are mediated in cis at some point, so this scenario is subsumed by scenario A (i.e. mediation).

##### H. Mediation by unobserved cis intermediary

Here, the SNP affects an unobserved intermediary in cis, which then has pleiotropic effects on a gene's expression levels and the complex trait. The intermediary can be the expression levels/splicing/activity of another gene, and can also refer to any other molecular process. Note that this scenario refers specifically to *cis* intermediaries; the distinction between *cis* intermediaries and *trans* intermediaries is that SNP effect sizes on known molecular phenotypes are much larger in cis than in trans[1–3]. In scenario H, it is not appropriate to assume that eQTL effect sizes and pleiotropic effect sizes are independent, as they are both affected by a common intermediary. Thus, this scenario can contribute to nonzero estimates of $h^2_{med}$. Because this scenario does not strictly speaking involve mediation through the gene's expression levels, its contribution to $h^2_{med}$ might be viewed as spurious.

However, there are several reasons why scenario H's potential contribution to $h^2_{med}$ is likely not of major concern. First, if the intermediary represents the expression levels of another gene, then scenario H's contribution to $h^2_{med}$ is justifiable given that mediation is actually occurring through the expression levels of the intermediary. Second, because we perform regression using all SNPs in the genome, scenario H must be pervasive across most loci in the genome in order for it to have a substantial impact on estimates of $h^2_{med}$. Third, if the intermediary does not refer to the expression levels of a gene (e.g. it represents splicing or coding changes in a gene, or it represents some unknown molecular process), we argue that the contribution of the non-causal gene to $h^2_{med}$ is still of biological interest due to the fact that the gene's expression levels are correlated with a truly causal intermediary.

Should the third scenario be pervasive across the genome and have a substantial contribution toward $h^2_{med}$, we can amend our definition of $h^2_{med}$ in order to accommodate it. In "Definition of $h^2_{med}$," we refer to our estimate of $h^2_{med}$ as the heritability mediated and/or "tagged" by assayed expression levels, and we state that "tagging" occurs due to assayed expression levels acting as a proxy to expression levels in causal cell types/contexts. We can extend this definition of $h^2_{med}$ so that tagging also refers to assayed expression levels acting as a proxy to unobserved intermediaries *other than* expression levels. We believe that this extended definition of $h^2_{med}$ is of equal biological interest to a definition of $h^2_{med}$ that excludes tagging of unobserved intermediaries other than expression levels.

##### I. Mediation by unobserved trans intermediary

Here, the SNP affects an unobserved intermediary in trans, with then has pleiotropic effects on gene expression levels in cis and the complex trait. The only difference between scenario I and scenario H is that here the intermediary is affected in trans rather than in cis. Because trans-effects on molecular phenotypes are much smaller than cis-effects, the contribution of scenario I to nonzero $h^2_{med}$ will be much smaller than the contribution of scenario H (see "Reverse mediation" for related intuition).

#### Reverse mediation

In our generative model, we do not model the effects of reverse mediation, which we define as the scenario in which a SNP influences the complex trait independently of the SNP's effects on the expression of a gene, and the complex trait itself then influences the expression of the gene. Such a scenario will induce a genetic correlation between the gene's expression and the complex trait and could potentially bias our estimates of $h^2_{med}$. However, we posit that the bias (if present at all) is negligible for the following reasons. (1) Because we use an external expression panel to estimate eQTL effect sizes, the complex trait of interest must be represented in the expression panel samples in order for its effects on expression to be present in our analyses[4,5]. Thus, we can rule out the possibility of reverse mediation influencing our results for any disease phenotypes. (2) Assuming that the complex trait of interest is represented in the expression panel samples, the total bias in estimates of $h^2_{med}$ caused by reverse mediation is guaranteed to be very small under the assumption that SNP effect sizes on a trait are much smaller than eQTL effect sizes[5], which is true for polygenic traits. To illustrate this point, it is useful to think of $h^2_{med}$ in terms of the covariance between eQTL effect sizes $\beta$ and total SNP effect sizes $\omega$ (for simplicity, we assume that each gene has only one eQTL):

$$h^2_{med} = \sum_i^G Cov(\beta_i, \omega_i)^2 / Var(\beta_i)$$

Assuming that $Cov(\beta_i, \omega_i)$ for SNP $i$ is nonzero due to proper mediation, we have the following expression for $Cov(\beta_i, \omega_i)^2_{med}$:

$$\omega_i = \beta_i \alpha_i + \gamma_i$$
$$Cov(\beta_i, \omega_i)^2_{med} = \alpha_i^2 Var(\beta_i)^2$$

A typical trait has $h^2_{med} = 0.1$ and $Var(\beta_i) = 0.05$, so a realistic value for $\alpha_i^2$ is $\frac{h^2_{med}}{GVar(\beta_i)} = \frac{0.1}{20000 \cdot 0.05} = 0.0001$ (we assume here that all genes are causal; the proportion of causal genes will not affect the points conveyed by these calculations so long as the number of causal SNPs is at least as large as the number of causal genes; see below). Thus, we have $Cov(\beta_i, \omega_i)^2_{med} \approx 0.0001 \cdot 0.05^2 = 2.5 \times 10^{-7}$. On the other hand, if we assume that $Cov(\beta_i, \omega_i)$ for SNP $i$ is nonzero due to reverse mediation, we have the following expression for $Cov(\beta_i, \omega_i)^2_{revmed}$:

$$\beta_i = \theta_i \sum_j^M \omega_j + \beta_{i(SNP)}$$
$$Cov(\beta_i, \omega_i)^2_{revmed} = \theta_i^2 Var(\omega_i)^2$$

Here, $\theta_i$ represents the effect size of the complex trait on the expression of gene $i$ and $\beta_{i(SNP)}$ represents the direct effect size of SNP $i$ on gene $i$ without the effects of reverse mediation. The upper limit for $\theta_i^2$ is $\beta_i^2/h^2$, which occurs if $\beta_{i(SNP)} = 0$. Thus, we can rewrite the above as

$$Cov(\beta_i, \omega_i)^2_{revmed} \leq \beta_i^2/h^2 Var(\omega_i)^2$$

A typical complex trait has $h^2 = 0.5$. If we assume that the number of causal SNPs is at least as large as the number of causal genes, we have $Var(\omega_i) \leq h^2/G = 2.5 \times 10^{-5}$. Thus, we have $Cov(\beta_i, \omega_i)^2_{revmed} \leq 0.05/0.5 \cdot (2.5 \times 10^{-5})^2 = 6.25 \times 10^{-11}$. The reason why $Cov(\beta_i, \omega_i)^2_{revmed}$ is orders of magnitude smaller than $Cov(\beta_i, \omega_i)^2_{med}$ is that $Cov(\beta_i, \omega_i)^2_{revmed}$ involves the product of the *square* of the squared per-SNP effect on complex trait and the squared SNP effect on gene expression, while $Cov(\beta_i, \omega_i)^2_{med}$ involves the product of the squared per-gene effect on complex trait and the *square* of the squared SNP effect on gene expression. Because individual SNP effects on gene expression are much larger than individual SNP/gene effects on a polygenic trait, squaring the latter causes the overall magnitude of $Cov(\beta_i, \omega_i)^2$ to be much smaller than squaring the former.

#### Relationship between MESC and stratified LD score regression

MESC is similar in form to stratified LD score regression (S-LDSC), which aims to estimate total heritability partitioned across SNP categories from summary statistics[6,7]. In particular, the $\tau$ coefficient estimated by S-LDSC is directly related to the $\pi$ coefficient we obtain from MESC in equation (20) (in Methods). In S-LDSC, the variance of total effect size of SNP $k$ on the trait ($\omega_k$) is modeled as follows:

$$Var(\omega_k) = \sum_c a_c(k)\tau_c$$

where $a_c$ refers to a continuous-valued SNP annotation. Meanwhile, in MESC, $Var(\omega_k)$ is modeled as follows:

$$Var(\omega_k) = \sum_d \pi_d \sum_{i \in D} \beta_{ik}^2 + \sum_{c: k \in C} \tau_c^{nonmed}$$

We label $\tau_c^{nonmed}$ as so in order to distinguish it from $\tau_c$ used in S-LDSC. Here, note that we can treat the value $\sum_{i \in D} \beta_{ik}^2$ as a continuous SNP annotation $a_c$, which means that the expression scores $\mathcal{L}_{k;d}$ used in equation (20) are equivalent to LD scores with continuous annotation $a_c(k) = \sum_{i \in D} \beta_{ik}^2$. Thus, $\pi_d$ as defined above and in equation (20) is equivalent to $\tau_c$ as defined in S-LDSC for the SNP annotation that corresponds to $\sum_{i \in D} \beta_{ik}^2$.

The main implication for this equivalence between MESC and S-LDSC is that significantly nonzero $\pi_d$ as estimated by MESC can be interpreted as significantly nonzero $\tau_c$ from S-LDSC. There is considerable interest in identifying SNP annotations with significantly nonzero $\tau_c$ conditional on the baselineLD model and other SNP annotations[6–12], since this means that the SNP annotation is informative for explaining trait heritability *beyond* the set of comprehensive but non-trait-specific SNP annotations contained in the baselineLD model (as well any additional SNP annotations included in overall model). When using MESC, we also include all SNP annotations in the baselineLD model in our analyses, albeit for a different purpose than in studies using S-LDSC; our reason for including the baselineLD model is to account for correlations between the magnitude of non-mediated effect sizes and eQTL effect sizes (see "Model assumptions"). Nevertheless, we can still interpret significantly nonzero $\pi_d$ for a given gene category $D$ as implying that the SNP annotation corresponding to the eQTL effect sizes of all SNPs on genes in $D$ is informative for explaining trait heritability beyond the baselineLD model.

#### Relationship between MESC and Mendelian randomization

In this section, we describe the motivation behind the regression procedure carried out in MESC and compare it to Mendelian randomization (MR). Our goal is to estimate $h^2_{med}$, where $h^2_{med} = \sum_i^G \sum_j^M \beta_{ij}^2 \alpha_i^2$. One way we could estimate $h^2_{med}$ would be to first estimate $\alpha_i^2$ for each individual gene, then multiply $\alpha_i^2$ by the cis-heritability of the gene and sum up this quantity across all genes to obtain $h^2_{med}$. In principle, we could estimate $\alpha_i$ for each individual gene $i$ using some type of MR approach, where the exposure of interest is the expression level of gene $i$, and the outcome is the trait. However, typical MR approaches are problematic for this aim. In the presence of non-mediated effects of genetic variants on the trait, MR is highly underpowered to estimate $\alpha_i$ with a small number of genetic instruments[13,14]. This is a common scenario if we use gene expression as the exposure, since many genes have only a few detectable cis-eQTLs for their expression[15]. Alternatively, we could consider a MR approach with multiple genetic variants, which in principle can distinguish mediated from non-mediated effects so long as the InSIDE (instrument strength independent of direct effect) assumption holds[14] (Note that the InSIDE assumption is essentially the same as the pleiotropy-eQTL independence assumption we describe in "Model assumptions"). However, this approach is highly underpowered in the common scenario that genes have only a few detectable cis-eQTLs[13], and this approach cannot be applied to genes with only one cis-eQTL. In summary, we cannot use typical MR approaches to estimate $h^2_{med}$ due to the sparse cis-genetic architecture of gene expression.

Unlike MR approaches, MESC is able to estimate $h^2_{med}$ in the presence of sparsity of eQTLs for individual genes by estimating gene-trait effects *across* many genes. To illustrate this, we contrast MESC and MR with multiple genetic variants (see Supplementary Figure 31 for an illustration). MR with multiple genetic variants essentially involves regressing SNP-trait effects on eQTL effects for a single gene. The slope from this regression will be the effect of the gene on the trait given that the InSIDE assumption is satisfied. Meanwhile, MESC essentially involves regressing *squared* SNP-trait effects on *squared* eQTL effects *summed across* a set of genes. The slope from this regression will be the *average squared* effect of all genes in the gene set on the trait given that both the pleiotropy-eQTL independence assumption (which is effectively the InSIDE assumption extended across eQTLs for all genes in the gene set) and an additional assumption are satisfied. This additional assumption is that eQTL effect sizes are independent of gene-trait effect sizes across genes (see "Model assumptions"). We can then calculate $h^2_{med}$ with our average gene-trait effect estimate using an equivalent definition of $h^2_{med}$ that models $\beta$ and $\alpha$ as random variables (See "Definition of $h^2_{med}$"). Thus, MESC can still reliably estimate $h^2_{med}$ if individual genes have one or a small number of eQTLs, since it essentially aggregates information about gene-trait effects across many genes.

In summary, MESC can be conceptualized as an analogue to MR that models exposure effects as random and jointly estimates the average squared effect of multiple exposures on an outcome.

#### Use of eQTL summary statistics in MESC

We can use summary statistics from eQTL studies to estimate $h^2_{med}(\mathcal{D}_d)$ since expression score $\mathcal{L}_{k;d}$ is equivalent to the sum of marginal OLS estimates of eQTL effect sizes for SNP $k$ on genes in $\mathcal{D}_d$ ($\sum_{i\in\mathcal{D}_d}\hat{\beta}^2_{ik(sumstat)}$) modulo an error term that depends on $|\mathcal{D}_d|$ and the sample size of the eQTL study. This error term will be captured by the intercept during regression. To illustrate this, we model the expression of gene $i$ for $N_{exp}$ expression panel samples as follows:

$$\boldsymbol{y}_{i(exp)} = \boldsymbol{X}\boldsymbol{\beta}_i + \boldsymbol{\epsilon}_{i(exp)}$$

where $\boldsymbol{y}_{i(exp)}$ is an $N_{exp}$-vector of gene expression measurements (standardized to mean 0 and variance 1), $\boldsymbol{X}$ is an $N_{exp} \times M$ genotype for $M$ SNPs (standardized to mean 0 and variance 1), $\boldsymbol{\beta}_i$ is an $M$-vector of eQTL effect sizes, and $\boldsymbol{\epsilon}_{i(exp)}$ is an $N_{exp}$-vector of environmental effects. Under this model, we have

$$\begin{aligned}
E\left[\sum_{i\in\mathcal{D}_d}\hat{\beta}^2_{ik(sumstat)}\right] &= \sum_{i\in\mathcal{D}_d}\left(\sum_j^M \hat{r}^2_{jk}\beta^2_{ij} + \frac{E[\epsilon^2_{i(exp)}]}{N_{exp}}\right)\\
&= \sum_{i\in\mathcal{D}_d}\sum_j^M \hat{r}^2_{jk}\beta^2_{ij} + \sum_{i\in\mathcal{D}_d}\frac{1-E[h^2_{cis}]}{N_{exp}}\\
&= \sum_{i\in\mathcal{D}_d}\sum_j^M r^2_{jk}\beta^2_{ij} + \frac{|\mathcal{D}_d|E[h^2_{cis}]}{N_{exp}} + \frac{|\mathcal{D}_d|(1-E[h^2_{cis}])}{N_{exp}}\\
&= \mathcal{L}_{k;d} + \frac{|\mathcal{D}_d|}{N_{exp}}
\end{aligned}$$

Thus, we can use the following alternate form of equation (20) (in Methods) to perform regression:

$$E[\chi^2_k] = N\sum_c \tau_c \ell_{k;c} + N\sum_d \pi_d \sum_{i\in\mathcal{D}_d}\hat{\beta}^2_{ik(sumstat)} + 1 + \frac{Nh^2_{med;causal}}{N_{exp}E[h^2_{cis}]}$$

#### MESC with assayed expression scores

In this section, we show that when we carry out the regression procedure described in "MESC with summary statistics" using expression scores in assayed tissues $T$ rather than in causal cell types/contexts, we obtain an estimate of $h^2_{med;assayed}(T)$.

Let $\beta$ represent cis-eQTL effect sizes in causal cell types/contexts for the trait, and $\beta'$ represent cis-eQTL effect sizes in assayed tissues $T$. For simplicity, assume no sampling noise and non-mediated effects in GWAS $\chi^2$ statistics. Upon regressing GWAS $\chi^2$ statistics on expression scores in assayed tissues (see equation (17)), we have

$$\begin{aligned}
\alpha'^2 &\approx \frac{1}{G}\sum_i^G \frac{Cov(\chi^2, \sum_j^M r^2_j {\beta'}^2_{ij})}{Var(\sum_j^M r^2_j {\beta'}^2_{ij})}\\
&\approx \frac{1}{G}\sum_i^G \frac{Cov(\sum_j^M r^2_j \alpha^2_i \beta^2_{ij}, \sum_j^M r^2_j {\beta'}^2_{ij})}{Var(\sum_j^M r^2_j {\beta'}^2_{ij})}\\
&\approx E[\alpha^2]\frac{1}{G}\sum_i^G \frac{\ell^2 Cov(\beta^2_i, {\beta'}^2_i)}{\ell^2 Var({\beta'}^2_i)}\\
&\approx E[\alpha^2]\frac{1}{G}\sum_i^G \frac{Cov(\beta^2_i, {\beta'}^2_i)}{Var({\beta'}^2_i)}
\end{aligned}$$

Here, $\ell^2 = Var(\sum_j^M r^2_j)$. The third and fourth line follow given that $r^2$ is independent of $\alpha$, $\beta$, and $\beta'$. See "MESC with assayed eQTL effect sizes" for the remainder of the derivation.

#### Stratified MESC derivation

Starting from equation (14):

$$E[\chi_k^2 \mid \boldsymbol{R}, \boldsymbol{B}] = N \sum_j^M E[\gamma_j^2 \mid \boldsymbol{R}, \boldsymbol{B}] \hat{r}_{jk}^2 + N \sum_i^G E[\alpha_i^2 \mid \boldsymbol{R}, \boldsymbol{B}] \sum_j^M \hat{r}_{jk}^2 \beta_{ij}^2 + NE[(\boldsymbol{\epsilon}')^2] \tag{7.1}$$

$$= N \sum_j^M \left( \sum_{c:j \in \mathcal{C}_c} \tau_c \mid \boldsymbol{R}, \boldsymbol{B} \right) \hat{r}_{jk}^2 + N \sum_i^G \left( \sum_{d:i \in \mathcal{D}_d} \pi_d \mid \boldsymbol{R}, \boldsymbol{B} \right) \sum_j^M \hat{r}_{jk}^2 \beta_{ij}^2 + NE[(\boldsymbol{\epsilon}')^2] \tag{7.2}$$

$$E[\chi_k^2] = N \sum_c \tau_c \sum_{j \in \mathcal{C}_c} \hat{r}_{jk}^2 + N \sum_d \pi_d \sum_{i \in \mathcal{D}_d} \sum_j^M \hat{r}_{jk}^2 \beta_{ij}^2 + NE[(\boldsymbol{\epsilon}')^2] \tag{7.3}$$

In order for (7.3) to be true, we must make the following assumptions:

- Within each gene category $\mathcal{D}_d$, $\pi_d$ is uncorrelated with the magnitude of eQTL effect sizes
- Within each SNP category $\mathcal{C}_c$, $\tau_c$ is uncorrelated with the magnitude of eQTL effect sizes
- $\pi_d$ is uncorrelated with the LD scores of eQTLs that affect genes in $\mathcal{D}_d$
- $\tau_c$ is uncorrelated with the LD scores of SNPs in $\mathcal{C}_c$

Since $E[\hat{r}_{jk}^2] \approx r_{jk}^2 + \frac{1}{N}$, we have

$$\begin{aligned}
E[\chi_k^2] &= N \sum_c \tau_c \sum_{j \in C_c} \left( r_{jk}^2 + \frac{1}{N} \right) + N \sum_d \pi_d \sum_{i \in \mathcal{D}_d} \sum_j^M \left( r_{jk}^2 \beta_{ij}^2 + \frac{\beta_{ij}^2}{N} \right) + NE[(\boldsymbol{\epsilon}')^2] \\
&= N \sum_c \tau_c \sum_{j \in C_c} r_{jk}^2 + \sum_c \sum_{j \in C_c} \tau_c + N \sum_d \pi_d \sum_{i \in \mathcal{D}_d} \sum_j^M r_{jk}^2 \beta_{ij}^2 + \\
&\quad \sum_d \sum_{i \in \mathcal{D}_d} \sum_j^M \left( \pi_d |\mathcal{D}_d| E[h_{cis}^2(\mathcal{D}_d)] \right) + NE[(\boldsymbol{\epsilon}')^2] \\
&= N \sum_c \tau_c \sum_{j \in C_c} r_{jk}^2 + h_{nonmed;causal}^2 + N \sum_d \pi_d \sum_{i \in \mathcal{D}_d} \sum_j^M r_{jk}^2 \beta_{ij}^2 + h_{med;causal}^2 + 1 - h_{nonmed;causal}^2 - h_{med;causal}^2 \\
&= N \sum_c \tau_c \sum_{j \in C_c} r_{jk}^2 + N \sum_d \pi_d \sum_{i \in \mathcal{D}_d} \sum_j^M r_{jk}^2 \beta_{ij}^2 + 1
\end{aligned}$$

Letting $\ell_{k;c} = \sum_{j \in C_c} r_{jk}^2$ and $\mathcal{L}_{k;d} = \sum_{i \in \mathcal{D}_d} \sum_j^M r_{jk}^2 \beta_{ij}^2$, we arrive at our main equation for stratified MESC:

$$E[\chi_k^2] = N \sum_c \tau_c \ell_{k;c} + N \sum_d \pi_d \mathcal{L}_{k;d} + 1$$

#### Impact of systemic differences in eQTL effect size magnitude between assayed vs. causal tissues

According to equation (13), we have

$$\alpha'^2 = r_g^2(T) E[\alpha^2] \frac{1}{G} \sum_i^G \sqrt{\frac{Var(\beta_i^2)}{Var({\beta'}_i^2)}}$$

There are reasonable scenarios in which $Var(\beta_i^2)$ can be systemically larger or smaller than $Var({\beta'}_i^2)$ across all genes (e.g. if causal genes for the trait are primarily influenced by cell type-specific eQTLs that are weaker/absent in assayed tissues). This will cause $\alpha'^2 \neq r_g^2(T)E[\alpha^2]$. However, because we multiply $\alpha'^2$ by $GE[h'^2_{cis}]$ to obtain $h^2_{med;assayed}$(T), we will still have $h^2_{med;assayed}(T) = r_g^2(T)h^2_{med;causal}$.

To illustrate this, consider a scenario where $\beta'^2$ is both correlated with $\beta^2$ and scaled by a factor $c$ relative to $\beta^2$. (13) thus becomes

$$\begin{aligned}\alpha'^2 &= r_g^2(T)E[\alpha^2]\frac{1}{G}\sum_i^G \sqrt{\frac{Var(\beta_i^2)}{Var(c\beta_i^2)}} \\ &= \frac{1}{c}r_g^2(T)E[\alpha^2]\end{aligned}$$

Note that scaling $\beta'^2$ by $c$ will not change the average squared correlation $r_g^2(T)$ between $\beta'^2$ and $\beta$. We then have

$$\begin{aligned}h^2_{med;assayed}(T) &= GE[\beta'^2]\frac{1}{c}r_g^2(T)E[\alpha^2] \\ &= GE[c\beta^2]\frac{1}{c}r_g^2(T)E[\alpha^2] \\ &= r_g^2(T)h^2_{med;causal}\end{aligned}$$

#### Comparing different methods of estimating expression scores

In this set of simulations, we evaluated the prediction accuracy and bias of different methods of estimating expression scores from simulated expression data with varying numbers of samples (Methods). Note that this set of simulations does not involve complex trait phenotypes; see Figure 2a and Supplementary Figure 1 for simulation results involving complex trait phenotypes.

In total, we compared five different methods of estimating expression scores $\mathcal{L}_k$ for SNP $k$. Here, $G$ represents genes within 1 Mb of SNP $k$, while $M$ represents SNPs within 1 Mb of SNP $k$:

1. *eQTL summary statistics.* $\hat{\mathcal{L}}_k = \sum_i^G \hat{\beta}^2_{ik(sumstat)}$, where $\hat{\beta}^2_{ik(sumstat)}$ represents the squared eQTL summary statistic of SNP $k$ for gene $i$.

2. *LASSO.* $\hat{\mathcal{L}}_k = \sum_i^G \sum_j^M r^2_{jk}\hat{\beta}^2_{ij(LASSO)}$, where $r^2_{jk}$ represents the squared correlation between SNP $j$ and SNP $k$, and $\hat{\beta}^2_{ij(LASSO)}$ represents squared *causal* eQTL effect sizes of SNP $j$ on gene $i$ estimated by LASSO[16].

3. *LASSO with REML correction.* $\hat{\mathcal{L}}_k = \sum_i^G \sum_j^M r^2_{jk}c_i\hat{\beta}^2_{ij(LASSO)}$. This method is identical to *LASSO* except that we scale $\hat{\beta}^2_{ij(LASSO)}$ by a factor $c_i$. We define $c_i = \hat{h}^2_{cis;i}/\sum_j^M \hat{\beta}^2_{ij(LASSO)}$, where $\hat{h}^2_{cis;i}$ is the expression cis-heritability of gene $i$ predicted by REML. This approach is the same as the one described in "Estimation of expression scores." For computational ease, we did not actually use REML to predict expression cis-heritability for each gene in each simulation, but rather we took the true expression cis-heritability of the gene and added a realistic amount of noise in order to simulate REML prediction error.

4. *BLUP.* $\hat{\mathcal{L}}_k = \sum_i^G \sum_j^M r^2_{jk}\hat{\beta}^2_{ij(BLUP)}$. Here, $\hat{\beta}^2_{ij(BLUP)}$ represents squared *causal* eQTL effect sizes of SNP $j$ on gene $i$ estimated by best linear unbiased predictor (BLUP)[17].

5. *BLUP with REML correction.* $\hat{\mathcal{L}}_k = \sum_i^G \sum_j^M r^2_{jk}c_i\hat{\beta}^2_{ij(BLUP)}$. Same as *LASSO with REML correction*, but using BLUP rather than LASSO.

We report mean prediction accuracy (in terms of $R^2$ between predicted and true expression scores) and bias (in terms of the slope from regressing predicted expression scores on true expression scores) with mean standard errors over 100 independent simulations. Across all expression panel sample sizes, we obtained the best prediction $R^2$ when using LASSO with REML correction (Supplementary Figure 2). The superior prediction $R^2$ of LASSO compared to other methods can be attributed to the fact that LASSO enforces a sparsity prior on effect sizes, which matches the sparse nature of cis-eQTL effect sizes. However, LASSO also produces biased estimates of effect sizes, which is why we need to scale the effects to match the expression cis-heritability estimated by REML (which is unbiased) to obtain unbiased estimates of expression scores. LASSO with REML correction gave us approximately unbiased estimates of expression scores at large enough sample sizes (>500). For OLS summary statistics, we observed unbiased estimates of expression scores at all sample sizes but inferior prediction $R^2$ to LASSO with REML correction. The remainder of the estimation methods were biased at all sample sizes and had comparable prediction $R^2$ to OLS summary statistics. We observed concordant results when varying the number of eQTLs per gene (Supplementary Figure 2a-c), eQTL window size (Supplementary Figure 2d), and REML prediction error (Supplementary Figure 2f). Prediction $R^2$ and bias across all methods varied when changing the mean cis-heritability of expression, though the relative performance of the five methods compared to each other remained consistent across different mean cis-heritability values (Supplementary Figure 2e).

Notably, we observed poor prediction $R^2$ of all methods for expression data sets of size 100-200, which is comparable to the size of most individual tissue expression panel data sets. This result suggests that we cannot reliably predict expression scores using available individual tissue expression panel data sets. In real data, this result is corroborated by the fact that we obtain low estimates of $h^2_{med}/h^2_g$ across all traits when using individual tissue expression panel data sets to estimate expression scores (Figure 5b).

#### Rare vs. common variant $h^2_{med}$

In all our analyses, we restrict the regression SNPs used by MESC to only Hapmap3 SNPs[18]. Because Hapmap3 SNPs essentially only tag common variants, by restricting to Hapmap3 SNPs we estimate the proportion of *common* disease heritability mediated by the cis-genetic component of gene expression ($h^2_{med(common)}/h^2_{common}$) (see following section for simulation results). We define $h^2_{med(common)}$ as $\sum_{j \in C} \sum_i \beta_{ij}^2 \alpha_i^2$ (given standardized genotypes and phenotypes), where $C$ represents the set of all SNPs with minor allele frequency (MAF) $> 0.05$, $\beta_{ij}$ represents the cis-eQTL effect size of SNP $j$ on gene $i$, and $\alpha_i$ represents the effect size of gene $i$ on disease. This quantity differs from the *total* disease heritability mediated by gene expression ($h^2_{med(common)} + h^2_{med(rare)}$), where $h^2_{med(rare)}$ is defined in the same manner as $h^2_{med(common)}$ but $C$ is replaced with the set of all SNPs with MAF $< 0.05$. We do not aim to estimate $h^2_{med(rare)}$ because doing so requires eQTL effect size estimates for rare variants, which cannot be reliably obtained from current expression panel data sets. Even if we had the data to estimate $h^2_{med(rare)}$ (i.e. many thousands of whole-genome sequencing expression samples), there are several reasons why we would expect the proportion of total disease heritability mediated by gene expression $(h^2_{med(common)} + h^2_{med(rare)})/(h^2_{common} + h^2_{rare})$ to be either similar or smaller than the quantity $h^2_{med(common)}/h^2_{common}$ that we estimate:

1. Most SNP heritability is explained by common variants[19,20]. Thus, we can expect the quantity $(h^2_{med(common)} + h^2_{med(rare)})/(h^2_{common} + h^2_{rare})$ to depend mostly on $h^2_{med(common)}$ and $h^2_{common}$.
2. Rare variant heritability has a much larger enrichment in coding regions than common variant heritability[21], suggesting that the effects of rare variants on disease tend to be mediated by protein-coding changes rather than changes in gene expression. Protein-coding changes are not reflected in $h^2_{med}$, so we would expect that $h^2_{med(rare)}/h^2_{rare} < h^2_{med(common)}/h^2_{common}$.

##### Role of singletons in $h^2_{med}$

A recent study[22] has shown that a substantial proportion of total expression cis-heritability (around 20%) in an expression panel of 360 individuals is explained by singletons with MAF $< 0.0001$ (i.e. singletons that are not observed in large genome reference panels). However, as mentioned above, rare variant effects do not contribute to $h^2_{med(common)}$ since they are not tagged by Hapmap3 SNPs. Furthermore, even if we had the data to estimate eQTL effect sizes for singletons, there is evidence that singletons contribute very little or nothing to disease heritability, as a recent study[23] has shown that virtually all narrow-sense heritability for height and BMI can be explained by SNPs with MAF $> 0.0001$ (which excludes the class of ultra-rare

SNPs defined as singletons in Hernandez et al.). In light of this result, we would not expect singleton effects on expression to substantially mediate any disease heritability.

#### Simulations under frequency-dependent genetic architectures

To evaluate the bias of MESC in estimating $h^2_{med(common)}$ in the presence of frequency-dependent genetic architectures (including rare and low-frequency variants), we conducted simulations in which both eQTL and GWAS per-allele effect size magnitude were inversely proportional to minor allele frequency, consistent with purifying selection acting on gene expression[22,24] and complex trait[19,20]. We conducted our simulations using real genotypes imputed to include rare and low-frequency variants from UK Biobank[25] ($N_{GWAS}$ = 100,000; $M$ = 1,539,668 SNPs from chromosome 1). We simulated cis-eQTLs for $G = 1000$ genes with variance of per-allele effect sizes proportional to $[p_i(1-p_i)]]^\alpha$, where $p$ is the minor allele frequency of SNP $i$ and $\alpha$ is a parameter ranging from $-0.33$ (corresponding to 5% of heritability explained by rare variants with MAF < 0.01 in our data set) to $-1.33$ (corresponding to 50% heritability explained by rare variants). We simulated frequency dependent non-mediated SNP effect sizes in a similar fashion as eQTL effect sizes. Finally, we simulated gene effect sizes on complex trait corresponding to $h^2_{med(common)}/h^2_{common} = 0.2$ and $h^2 = 0.5$. From these effect sizes, we simulated GWAS summary statistics, as well as eQTL summary statistics using a separate set of genotypes ($N_{eQTL}$ = 10,000).

We applied MESC to these summary statistics while performing the regression using only Hapmap3 SNPs (consistent with what we do in practice). In order to capture dependence between LD scores and GWAS effect sizes (which constitutes a model violation and leads to biased $h^2_g$ estimates if uncorrected[26,27]), we stratified regression SNPs by 10 MAF bins, which has been shown to adequately account for this dependence[7,27]. In all simulations, we obtained unbiased/slightly conservative estimates of $h^2_{med(common)}/h^2_{common}$ across diverse values of $\alpha$, including scenarios in which $\alpha$ for non-mediated effect sizes was different than $\alpha$ for eQTL effect sizes (Supplementary Figure 5). Thus, MESC is robust to frequency-dependent genetic architectures for both gene expression and disease.

#### Role of tissue specificity in explaining low heritability genes

Because we define the $h^2_{cis}$ of a gene by averaging individual-tissue $h^2_{cis}$ estimates across all tissues, a gene with low meta-tissue $h^2_{cis}$ can reflect two different scenarios: the gene has low individual-tissue $h^2_{cis}$ across many tissues, or the gene has high individual-tissue $h^2_{cis}$ in only one or a small number of tissues (i.e. the gene has tissue-specific eQTLs). To investigate the potential role of tissue-specific eQTLs in explaining low meta-tissue $h^2_{cis}$, we obtained three quantities for each gene: (1) the number of tissues in which the gene had a significantly nonzero $h^2_{cis}$ ($p < 0.05$), (2) the max $h^2_{cis}$ of the gene across all tissues, and (3) the average $h^2_{cis}$ of the gene across the tissues for which $h^2_{cis}$ was significantly nonzero. If the number of tissues in which the gene has truly nonzero $h^2_{cis}$ (an indicator of the tissue specificity of the gene) is the primary factor in determining the magnitude of the meta-tissue $h^2_{cis}$, we would expect that (1) be proportional to the magnitude of the meta-tissue $h^2_{cis}$, while (2) and (3) not be proportional to the magnitude of the meta-tissue $h^2_{cis}$. We observed that (1) was indeed proportional to the magnitude of the meta-tissue $h^2_{cis}$ (Supplementary Figure 20a); however, (2) and (3) were also proportional to the magnitude of the meta-tissue $h^2_{cis}$ (Supplementary Fig 20b,c), suggesting that statistical power due to the magnitude of $h^2_{cis}$, rather than tissue specificity, was primarily responsible for the fact that (1) was proportional to the meta-tissue $h^2_{cis}$. In summary, these results suggest that low $h^2_{cis}$ genes are not primarily genes with highly tissue-specific eQTLs, though we cannot rule out the possibility of tissue-specific eQTLs having some contribution to low $h^2_{cis}$ genes.

#### Impact of adding window around gene set to model

When estimating the $h^2_{med}$ enrichment of a given gene set, we stratify SNPs by the baselineLD model v2.0 in order to account for correlations between eQTL effects and non-mediated effects, and we stratify genes within the gene set into three bins to account for correlations between eQTL effects and gene effects. In addition to the 72 baselineLD model SNP annotations, we might consider adding an additional annotation that corresponds to all SNPs within a certain genomic distance (e.g. 100 Kb) of genes in the gene category. By including this annotation, we impose a stricter standard for identifying $h^2_{med}$ enrichment of the gene category.

To illustrate this, consider a scenario in which GWAS signal tends to physically localize around genes in a given gene set, but that none of the GWAS signal is actually mediated by the expression levels of those genes. Because cis-eQTLs also (by definition) physically localize around those genes, by chance we will observe that eQTLs for those genes will have larger GWAS effect sizes compared to the genomic background, in which case we will likely spuriously identify the gene set as having significant $h^2_{med}$ enrichment. By including a 100 Kb window around each gene, we require that eQTL effect size magnitude is correlated with GWAS effect size magnitude *within* the 100 Kb windows to detect significant $h^2_{med}$ enrichment, which will not occur if the GWAS signal is not mediated by the expression levels of the genes. In summary, including a SNP annotation corresponding to a window around each gene can eliminate false positive $h^2_{med}$ enrichment estimates that arise due to localization of GWAS signal around genes that is not mediated by gene expression.

In practice, we chose to include a 100 Kb window around genes, given precedence in the literature[9,12,28,29]. These studies report large heritability enrichment of a window of this size around many functional gene sets. When including this annotation for each gene set, we observed that the $h^2_{med}$ enrichment estimate for the gene set was very similar for most gene sets (Supplementary Figure 23), demonstrating that the $h^2_{med}$ enrichment of these gene sets was not due to coincidental overlap between non-mediated effects and eQTL effects near these genes. Given this result, we did not include the window around genes in any of our subsequent analyses, nor in any of the results we report in the manuscript.

#### Comparing MESC to other gene set enrichment methods

MESC can be used to prioritize disease-relevant gene sets using the $h^2_{med}$ enrichment of a gene set, defined as (proportion of $h^2_{med}$ in gene set) / (proportion of genes in gene set). Larger $h^2_{med}$ enrichment of the gene set suggests that the expression of genes in the gene set have larger causal effects on disease. Many other methods exist that also aim to prioritize causal gene sets using GWAS data[30–36]. MESC primarily differs from these other methods in that (1) it utilizes eQTL data, and (2) it specifically estimates causal effects of gene expression on disease, under a generative model for disease that connects SNP effects on gene expression to gene effects on disease. On the other hand, other popular methods for gene set enrichment analysis (e.g. MAGMA, DEPICT) are not based on eQTL data and do not model gene effects on disease. Instead, these methods prioritize gene sets under the assumption that causal genes should have more GWAS signal in close genomic proximity to them, which may not be true in some cases[37,38]. Thus, the two qualities above can make MESC desirable as a discovery tool, especially since eQTLs have been useful in elucidating the mechanistic basis of disease in many other settings[4,5,13,39–49].

However, there are also scenarios in which MESC will miss gene sets that play a causal role in disease. In particular, MESC focuses only on genes whose expression levels mediate the effects of GWAS hits, to the extent that can be detected in existing eQTL studies such as GTEx. SNP effects on disease might be mediated by mechanisms other than gene expression levels (e.g. protein-coding changes), or they may be mediated by gene expression levels in specific cell types or contexts that are not captured by existing eQTL studies. Moreover, a key drawback of MESC is that it produces large standard errors for small gene sets and thus can only be applied to large gene sets with more than 200 genes, whereas other methods can analyze gene sets of any size. Thus, we propose MESC as a complementary approach rather than replacement for other pathway enrichment methods.

To compare MESC to other pathway enrichment methods, we applied MAGMA[33] and DEPICT[32] to the same GWAS summary statistics for 26 traits with nominally significant $h^2_{med}$. We analyzed a total of 501 gene sets, which represent the intersection of gene sets we analyzed using MESC in our study and gene sets built into the DEPICT software. We ran MAGMA with default parameters. DEPICT requires that we specify a p-value threshold for defining significant GWAS loci; however, the recommended thresholds of $1 \times 10^{-5}$ and $5 \times 10^{-8}$ caused DEPICT to exceed its maximum number of loci for many traits. Thus, for each trait, we set the p-value threshold to the maximum of the following values that did not cause DEPICT to exceed its maximum number of loci: $1 \times 10^{-5}$, $5 \times 10^{-8}$, $5 \times 10^{-15}$, $5 \times 10^{-20}$, $5 \times 10^{-25}$, $5 \times 10^{-30}$, $5 \times 10^{-35}$, $5 \times 10^{-40}$, $5 \times 10^{-45}$, $5 \times 10^{-50}$. We then compared gene sets identified as significantly enriched by MAGMA and DEPICT to gene sets with significant $h^2_{med}$ enrichment (see Supplementary Table 9 for all estimates). Out of 13,206 total trait-gene set pairs, MESC identified 106 with significant $h^2_{med}$ enrichment (FDR $< 0.05$ while correcting for 13,206 $\times$ 3 = 39,078 hypotheses), compared to 85 for MAGMA and 957 for DEPICT (Supplementary Figure 27a). We observed correlated enrichment p-values across the three methods (MESC vs. MAGMA $R^2 = 0.14$, MESC vs. DEPICT $R^2 = 0.20$, MAGMA vs. DEPICT $R^2 = 0.10$) (Supplementary Figure 27b). Of the 106 significant trait-gene sets pairs identified by MESC, 32 were not detected as significant by either MAGMA or DEPICT (Supplementary Figure 27c), including biologically plausible trait-gene sets pairs such as "phospholipid metabolic process" for high-density lipoprotein level and "synapse part" for schizophrenia. These results demonstrate that MESC produces broadly concordant gene set enrichment estimates as the other methods, while also capturing unique signal that is present in only eQTL data.

#### Impact of environmental noise in gene expression measurements on expression-mediated heritability estimates

In this section, we show that the level of environmental noise in gene expression measurements (which differs across assays and affects both standardized eQTL effect sizes and the magnitude of expression cis-heritability, $h^2_{cis}$) does not impact our estimates of expression-mediated heritability $h^2_{med}$. In other words, $h^2_{med}$ depends on only the genetic component of gene expression levels. One consequence of this fact is that the magnitude of $h^2_{med}$ does not *a priori* depend on the magnitude of $h^2_{cis}$. For example, mean $h^2_{cis}$ can be very low in a given gene expression data set due to e.g. large stochastic fluctuations in gene expression levels or other sources of technical noise specific to the data set, while estimated $h^2_{med}$ from this gene expression data set can in principle be very high (i.e. close to total SNP heritability $h^2_g$).

To understand this intuitively, one can think of the units in which all SNP effect sizes operate. Recall our model for the effect size of SNP $j$ on complex trait:

$$\omega_j = \sum_i \beta_{ij}\alpha_i + \gamma_j$$

where $\omega_j$ represents the total effect size of SNP $j$ on the complex trait, $\beta_{ij}$ represents the effect size of SNP $j$ on the expression levels of gene $i$, $\alpha_i$ represents the effect size of gene $i$ on the complex trait, and $\gamma_i$ represents the non-mediated effect size of SNP $j$ on the complex trait. When complex trait and gene expression levels are standardized to zero mean and unit variance, $\beta_{ij}$ is expressed in terms of *additive increase in standardized expression levels per unit increase in standardized genotype* (which we abbreviate as $\frac{\text{std(expr)}}{\text{std(geno)}}$), while $\alpha_i$ is expressed in terms of *additive increase in standardized phenotype per unit increase of standardized expression levels* (which we abbreviate as $\frac{\text{std(pheno)}}{\text{std(expr)}}$). When we multiply $\alpha_i^2$ by $\sum_i \beta_{ij}^2$, we obtain a quantity corresponding to the heritability mediated by gene expression levels for SNP $j$ in units of $\left(\frac{\text{std(expr)}}{\text{std(geno)}}\right)^2 \left(\frac{\text{std(pheno)}}{\text{std(expr)}}\right)^2 = \left(\frac{\text{std(pheno)}}{\text{std(geno)}}\right)^2$.

Because std(expr) cancels out in this above product, the units in which gene expression levels are represented does not actually affect our final estimate of the heritability mediated by gene expression levels for SNP $j$ (provided that both $\alpha_i$ and $\beta_{ij}$ use the same units of expression). To elaborate, when we regress $\omega_j^2$ on $\sum_i \beta_{ij}^2$, we obtain an estimate of $E[\alpha^2]$ in units of $\left(\frac{\text{std(pheno)}}{\text{std(expr)}}\right)^2$. To obtain an estimate of per-SNP expression-mediated heritability, we then multiply $E[\alpha^2]$ by $E\left[\sum_i \beta_{ij}^2\right]$ (or equivalently $E[h^2_{cis}]$), which is in units of $\left(\frac{\text{std(expr)}}{\text{std(geno)}}\right)^2$. Suppose we were to scale $\sum_i \beta_{ij}^2$ by an arbitrary factor $c$, in which case our estimate of $E[\alpha^2]$ would be in units of $\left(\frac{\text{std(pheno)}}{c\cdot\text{std(expr)}}\right)^2$ and $E\left[c\sum_i \beta_{ij}^2\right]$ would be in units of $\left(\frac{c\cdot\text{std(expr)}}{\text{std(geno)}}\right)^2$. When multiplying $E[\alpha^2]$ by $E\left[c\sum_i \beta_{ij}^2\right]$, the product would be expressed in units of $\left(\frac{c\cdot\text{std(expr)}}{\text{std(geno)}}\right)^2 \left(\frac{\text{std(pheno)}}{c\cdot\text{std(expr)}}\right)^2 = \left(\frac{\text{std(pheno)}}{\text{std(geno)}}\right)^2$ and would thus be unchanged compared to before. This is essentially the same argument as made in the section "Impact of systemic differences in eQTL effect size magnitude between assayed vs. causal tissues" (Supplementary Note), in which we show that true differences in eQTL effect size magnitude agnostic of environmental noise in expression assays also do not affect our estimates of expression-mediated heritability.

Adding environmental noise to gene expression levels has the effect of scaling both squared standardized eQTL effect sizes and $h^2_{cis}$ by a constant factor, which we have shown above does not affect estimates of $h^2_{med}$. To illustrate this, consider the following generative model for the expression levels of gene $i$, in which genotypes are standardized to zero mean and unit variance but gene expression levels are *not* standardized:

$$\boldsymbol{y}_{i(exp)} = \boldsymbol{X}\boldsymbol{\beta}_i + \boldsymbol{\epsilon}_{i(exp)}$$

where $\boldsymbol{y}_{i(exp)}$ is a vector of *non-standardized* gene expression levels for gene $i$, $\boldsymbol{X}$ is a matrix of *standardized* genotypes, $\boldsymbol{\beta}_i$ is a vector of *non-standardized* cis-eQTL effect sizes for gene $i$, and $\boldsymbol{\epsilon}_{i(exp)}$ is a vector of environmental effects. In order to standardize cis-eQTL effect sizes (so that $\sum \beta^2_{i(std)} = h^2_{cis}$), we divide all non-standardized cis-eQTL effect sizes $\beta_i$ by $\sqrt{\sum \beta_i^2 + Var(\epsilon_{i(exp)})}$ to obtain $\beta_{i(std)}$. Now, note that adjusting the variance of the noise term $\epsilon_{i(exp)}$ is akin to scaling both $h^2_{cis}$ and $\beta^2_{i(std)}$ by the same constant factor. For example, let $Var(\epsilon_{i(exp)})$ be the original environmental variance, and let $Var(\epsilon'_{i(exp)})$ be the new environmental variance. Both original $h^2_{cis}$ and $\beta^2_{i(std)}$ are multiplied by the factor $\frac{\sum \beta_i^2 + Var(\epsilon_{i(exp)})}{\sum \beta_i^2 + Var(\epsilon'_{i(exp)})}$ to obtain the new $h^2_{cis}$ and $\beta^2_{i(std)}$.

In summary, we show that the level of environmental noise in gene expression panels (due to e.g. stochastic fluctuations in gene expression levels or other sources of assay noise) does not impact our estimates of $h^2_{med}$.

#### Prospects for estimating disease heritability mediated by trans-eQTLs

In all our analyses, we aim to estimate disease heritability mediated by gene expression *in cis*, rather than the full genetic component of gene expression that includes trans effects. In theory, we can also estimate disease heritability mediated by gene expression *in trans* using MESC, where we would simply replace cis-eQTL effect sizes with trans-eQTL effect sizes in all our analyses. However, trans-eQTLs are much more difficult to estimate than cis-eQTLs due to their much smaller effect sizes, impacting resulting estimates of $h^2_{med}$. In this section, we show that $h^2_{med}$ estimates produced by MESC are bounded by the average genetic prediction $r^2$ of gene expression multiplied by true $h^2_{med}$, so $r^2 < 1$ results in downward bias in estimated $h^2_{med}$ (note that here $r^2$ refers to the prediction accuracy of the only the genetic component of gene expression, which does not include environmental effects). For gene expression in cis, this downward bias is minimal at current sample sizes (see simulation result in Figure 2a), as we can obtain a prediction $r^2$ close to 1 for cis-eQTLs[4] (also see simulation in Supplementary Figure 2). However, for gene expression in trans, this downward bias becomes problematic at current sample sizes, since trans-eQTLs are highly polygenic and thus more difficult to estimate[3,15,50]. In order to obtain a reliable estimate of disease heritability mediated by trans-eQTLs, we would ideally want a genetic prediction $r^2$ of 0.8 or greater (comparable to the prediction $r^2$ of cis-eQTLs from currently available gene expression data sets), which we show requires expression panels on the order of 1,000,000 or more samples (see below). Note that these sample sizes are comparable to those needed for very accurate polygenic disease risk prediction. We also show that the expected prediction $r^2$ of trans-eQTLs from the largest gene expression data set (eQTLGen[50], $N$ = 31,684) is only 0.026, which is far too low to yield meaningful estimates of $h^2_{med}$. Thus, estimating disease heritability mediated by trans-eQTLs using MESC is not feasible with currently available gene expression data sets.

##### Relationship between genetic prediction $r^2$ of gene expression and magnitude of estimated $h^2_{med}$

Let $\beta$ represent eQTL effect sizes (either cis or trans) in causal cell types/contexts for the trait, let $\beta'$ represent eQTL effect sizes in assayed tissues $T$, and let $\hat{\beta}' = \beta' + \epsilon$ represent *estimated* eQTL effect sizes in assayed tissues $T$, where $\epsilon$ is a noise term. We assume that $\epsilon$ is independent of $\beta'$. Upon regressing squared GWAS effects sizes $\omega^2$ on squared estimated eQTL effect sizes $\sum_i^G \hat{\beta}'^2_i$, the estimate of the coefficient $\hat{\alpha}'^2$ is:

$$\hat{\alpha}'^2 \approx E[\alpha^2]\frac{1}{G}\sum_i^G \frac{Cov(\beta_i^2, \hat{\beta}_i'^2)}{Var(\hat{\beta}_i'^2)}$$
$$\approx E[\alpha^2]\frac{1}{G}\sum_i^G \frac{Cov(\beta_i^2, \beta'^2_i + \epsilon_i^2)}{Var(\hat{\beta}_i'^2)}$$
$$\approx E[\alpha^2]\frac{1}{G}\sum_i^G \frac{Cov(\beta_i^2, \beta'^2_i)}{Var(\hat{\beta}_i'^2)}$$

The first line follows from the same derivation as "MESC with eQTL effect sizes in non-causal tissues." As before, we define $r_g^2(T) = \frac{1}{G}\sum_i^G \frac{Cov(\beta_i^2, \beta_i'^2)}{\sqrt{Var(\beta_i^2)Var(\beta_i'^2)}}$ as the average squared genetic correlation between expression in assayed tissues $T$ vs. in causal cell types/contexts. Given this definition, we have:

$$\hat{\alpha}'^2 \approx r_g^2(T)E[\alpha^2]\frac{1}{G}\sum_i^G \frac{\sqrt{Var(\beta_i^2)Var(\beta_i'^2)}}{Var(\hat{\beta}_i'^2)}$$

We can establish an upper bound for $\hat{\alpha}'^2$ in terms of the genetic prediction accuracy of expression. Let $r^2_{pred}(T) = \frac{1}{G}\sum_i^G \frac{Cov(\hat{\beta}_i'^2, \beta_i'^2)}{\sqrt{Var(\hat{\beta}_i'^2)Var(\beta_i'^2)}}$ represent the average squared genetic prediction accuracy of expression across genes in tissues $T$. Note that $\frac{\sqrt{Var(\beta_i^2)Var(\beta_i'^2)}}{Var(\hat{\beta}_i'^2)} \leq \frac{Cov(\hat{\beta}_i'^2, \beta_i'^2)}{\sqrt{Var(\hat{\beta}_i'^2)Var(\beta_i'^2)}}$ under the assumption that $Var(\beta_i^2) \approx Var(\beta'^2_i)$. To illustrate this, see that the numerators of both sides of the inequality are equivalent: $Cov(\hat{\beta}_i'^2, \beta_i'^2) = Cov(\beta'^2_i + \epsilon_i^2, \beta_i'^2) = Cov(\beta'^2_i, \beta_i'^2) = Var(\beta'^2_i) \approx Var(\beta_i^2)$ on the left side, and $\sqrt{Var(\beta_i^2)Var(\beta_i'^2)} \approx \sqrt{Var(\beta_i^2)Var(\beta_i^2)} \approx Var(\beta_i^2)$ on the right side. However, in the denominators, $Var(\hat{\beta}_i'^2) \geq \sqrt{Var(\hat{\beta}_i'^2)Var(\beta_i'^2)}$, since $Var(\hat{\beta}_i'^2) \geq Var(\beta'^2_i)$. Thus, we have:

$$\hat{\alpha}'^2 \leq r^2_{pred}(T)r_g^2(T)E[\alpha^2]$$

We can multiply $\hat{\alpha}'^2$ by $GE[h'^2_{cis}]$ to obtain an estimate of $h^2_{med;assayed}(T)$ that has the following property:

$$\hat{h}^2_{med;assayed}(T) \leq r^2_{pred}(T)h^2_{med;assayed}(T)$$

##### Expected genetic prediction $r^2$ of gene expression in trans

Unlike cis-eQTLs, trans-eQTLs are known to be polygenic. Thus, we can invoke the following equation that relates sample size to polygenic prediction accuracy for gene expression in trans using the best linear unbiased predictor (BLUP)[51,52]:

$$r^2_{pred;trans}(T) = \frac{1}{G}\sum_i^G \frac{h^2_{i;trans}}{h^2_{i;trans} + \frac{M}{N}(1 - r^2_{pred;trans}(T))}$$
$$\approx \frac{N}{M}\frac{1}{G}\sum_i^G h^2_{i;trans}$$

where $h^2_{i;trans}$ is the expression trans-heritability of gene $i$, $M$ is the effective number of independent SNPs (approximately 60,000[53]), and $N$ is expression panel sample size. The largest expression panel available to date is from eQTLGen[50], with $N$ = 31,684 in blood. The average $h^2_{trans}$ of expression is around 0.05[3,4,54]. Thus, we can expect $r^2_{pred;trans}(T)$ trained on eQTLGen data to be around $\frac{31{,}684\cdot 0.05}{60{,}000} = 0.026$, which is far too low to yield meaningful estimates of $h^2_{med}$. In order to obtain $r^2_{pred;trans}(T)$ of 0.8 (comparable to the prediction $r^2$ of gene expression in cis from current expression panels), we would need $\frac{0.8\cdot 60{,}000}{0.05} = 960{,}000$ samples.

#### Choice of 10 traits for display in Figure 5

Starting with the full set of 26 traits with $h^2_{med}/h^2_g$ greater than 0 with nominal significance (p < 0.05), we pruned genetically correlated traits as follows. First, we selected the pair of traits with the greatest genetic correlation (estimated using cross-trait LD score regression[55]). Between the pair of selected traits, we then retained the trait with the larger $h^2_g$ z-score (estimated using stratified LD score regression[6]). We repeated this procedure until 10 traits were left.

#### Simulation: Supplementary Figure 2

We selected a 20 Mb region on chromosome 1 (base pair coordinates 60,000,000 to 80,000,000), which contained 8,604 SNPs. 100 genes were simulated within this region, with the average cis-heritability across all genes set at $\overline{h^2_{cis}} = 0.05$ or 0.01. For each gene, we simulated 1, 5, or 10 cis-eQTLs with locations randomly selected within a random 1 Mb or 10 Kb window within the overall 20 Mb region. For simulations with one eQTL per gene, the effect size for the eQTL was drawn from $\mathcal{N}(0,\, \overline{h^2_{cis}})$. For simulations with more than one eQTL per gene, one eQTL was randomly selected to explain 80% of the total heritability of the gene and had an effect size drawn from $\mathcal{N}(0,\, 0.8\overline{h^2_{cis}})$. The remaining eQTLs had effect sizes drawn from $\mathcal{N}(0,\, 0.2\overline{h^2_{cis}}/(N_{eQTL}-1))$, where $N_{eQTL}$ is the total number of eQTLs per gene. Expression values were simulated for each gene using an additive generative model with previously simulated effect sizes and environmental noise drawn from $\mathcal{N}(0,\, 1-\overline{h^2_{cis}})$. LASSO and BLUP prediction of eQTL effect sizes was performed using all SNPs within 1Mb of each simulated gene. To simulate REML prediction error in expression heritability estimates, we added noise drawn from $\mathcal{N}(0,\, 0.01^2)$ to the true heritability values, which is consistent with the average standard error of GCTA estimates of expression cis-heritability across all GTEx samples (Supplementary Figure 32).

#### Simulation: Supplementary Figure 3

All effect sizes and complex trait phenotypes were simulated in the same manner as Figure 2a. We simulated expression phenotypes for 1, 5, or 10 tissues with 200 samples per tissue using the same eQTL effect sizes used to generate complex trait phenotypes. We estimated expression scores in each individual tissue. We meta-analyzed expression scores across tissues by averaging the causal squared LASSO-predicted eQTL effect sizes across all tissues for each gene (after scaling the effect sizes to the estimated expression cis-heritability). We then used these averaged causal eQTL effect sizes to compute expression scores by multiplying them by the element-wise squared LD matrix. This meta-analysis procedure is the same as the one described in "Meta-analysis of expression scores" in Methods.

#### Simulation: Supplementary Figure 5

For all simulations, we set $\overline{h^2_{cis}} = 0.05$, $h^2_{med} = 0.1$, and $h^2 = 0.5$. For each of 1000 genes, we randomly selected 5 SNPs within a random 1 Mb window to act as cis-eQTLs with effect sizes drawn from $\mathcal{N}(0, [p_i(1-p_i)]^{\alpha_{eQTL}})$, where $p_i$ is the MAF of the given SNP $i$. To avoid extreme effect sizes for singletons or doubletons, effect sizes for SNPs with MAF < 0.01 were drawn from $\mathcal{N}(0, [0.01(1-0.01)]^{\alpha_{eQTL}})$. $\alpha_{eQTL}$ was set to either -0.33, -0.60, -0.99, or -1.33. Finally, we scaled the effect sizes of the cis-eQTLs so that the sum of their squared effects equalled $\overline{h^2_{cis}}$. Expression phenotypes were simulated for each gene with environmental noise drawn from $\mathcal{N}(0,\, 1-\overline{h^2_{cis}})$.

Similarly, we simulated non-mediated effect sizes for each SNP from $\mathcal{N}(0, [p_i(1-p_i)]^{\alpha_{GWAS}})$, or from $\mathcal{N}(0, [0.01(1-0.01)]^{\alpha_{GWAS}})$ for SNPs with MAF < 0.01. $\alpha_{GWAS}$ was set to any of the same four values $\alpha_{eQTL}$ could take on (without necessarily being equal to $\alpha_{eQTL}$). We then scaled these effect sizes so that the sum of the squared effects equalled $h^2 - h^2_{med}$. We simulated gene-trait effect sizes for each gene from $\mathcal{N}(0,\, (h^2_{med}/(G\overline{h^2_{cis}}))$. Complex trait phenotypes were simulated with environmental noise drawn from $\mathcal{N}(0, 1-h^2_g)$.

Expression scores were estimated by computing eQTL summary statistics from the simulated expression panel. In-sample LD scores were computed for all 1,539,668 SNPs from the 100,000 GWAS samples. Regression was performed using only Hapmap3 SNPs.

#### Simulation: Supplementary Figure 6

We set the overall $h^2_{med}/h^2_g = 0.4$. We simulated a gene category containing 200 random genes from the 1000 total genes. For the null scenario, we drew gene-trait effect sizes for all genes, including genes in the gene category, from $\mathcal{N}(0, h^2_{med}/(G\overline{h^2_{cis}}))$. For the enriched scenario, we simulated gene-trait effect sizes so that $h^2_{med}$ within the gene category was 2x the $h^2_{med}$ of genes outside of the category. eQTL effect sizes, non-mediated SNP effect sizes, expression phenotypes, and complex trait phenotypes were simulated in the same manner as in Figure 2a.

#### Simulation: Supplementary Figure 8

We set $\overline{h^2_{cis}} = 0.05$. We simulated 1 eQTL per gene with effect size drawn from $\mathcal{N}(0, \overline{h^2_{cis}})$. To simulate 10x enrichment of eQTLs in coding, TSS, and conserved regions, we selected eQTL locations so that 10x more eQTLs per SNP were located in the three SNP categories than the remainder of the genome. We simulated non-mediated effect sizes so that the heritability enrichment of the three SNP categories was 10x. Gene-trait effect sizes, expression phenotypes, and complex trait phenotypes were simulated in the same manner as in Figure 2a.

### Supplementary Figures

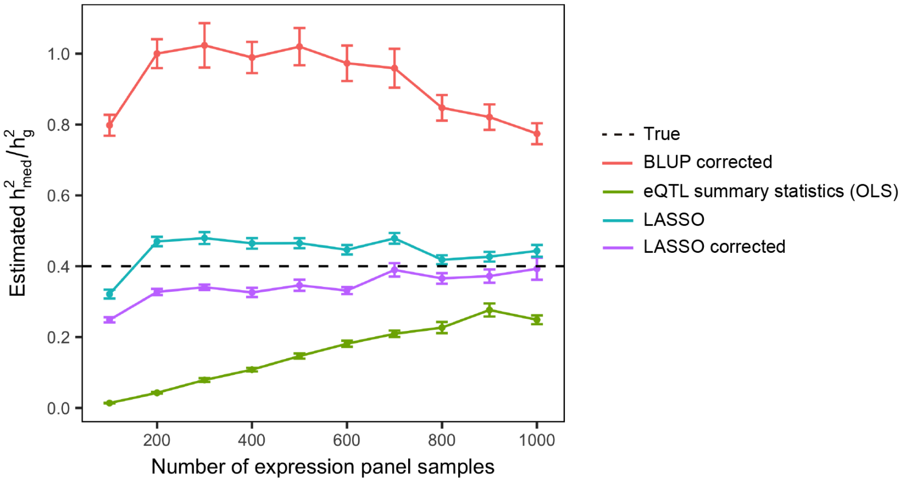


**Supplementary Figure 1. Impact of different methods of estimating expression scores on estimates of $h^2_{med}/h^2_g$.** For this simulation, $h^2_{med}/h^2_g = 0.4$. All other simulation parameters were the same as in Figure 2a. We exclude results for expression scores estimated using BLUP, since $h^2_{med}/h^2_g$ estimates obtained from these expression scores were severely upwardly biased (i.e. greater than 1 at all sample sizes).

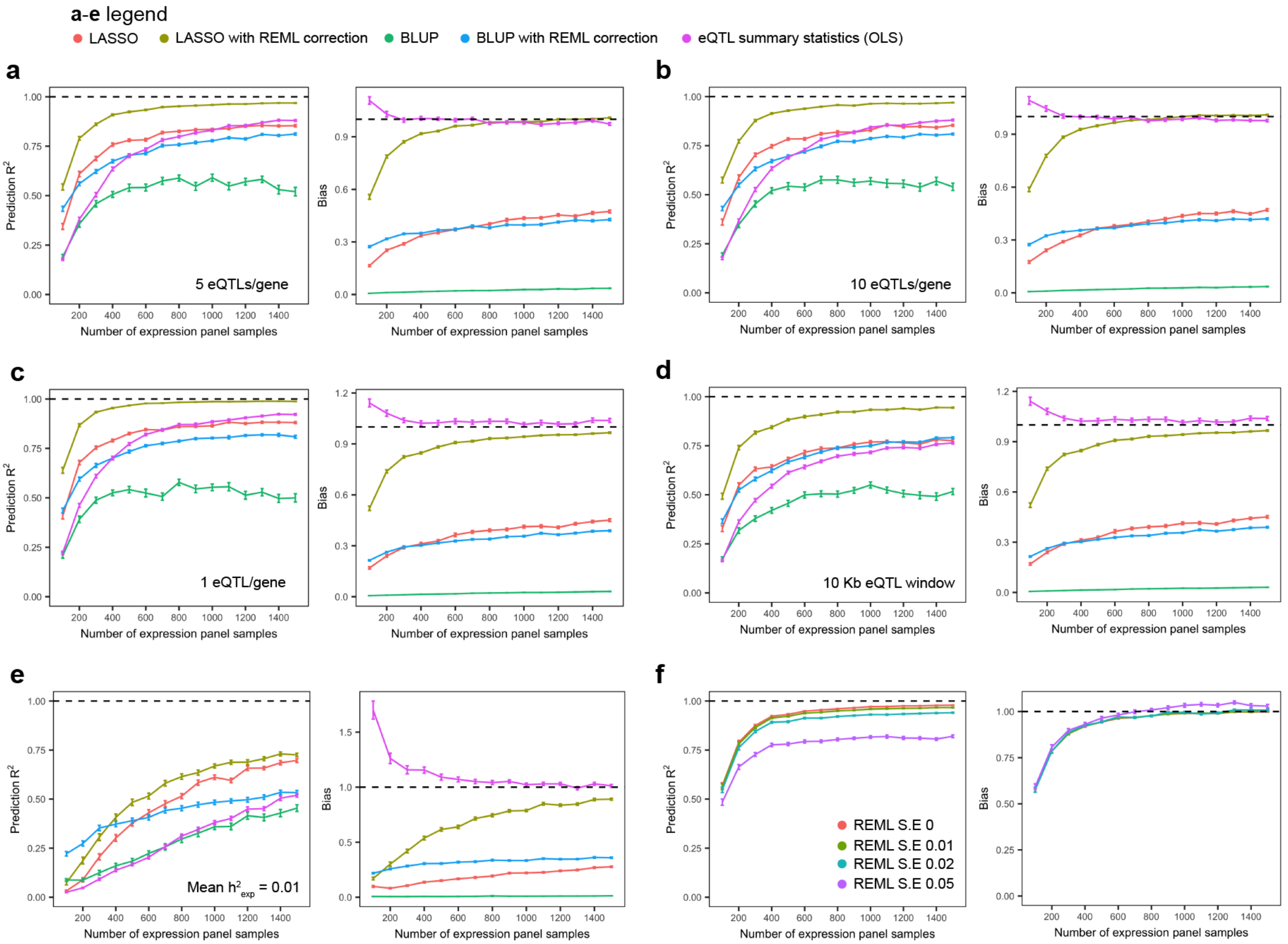


**Supplementary Figure 2. Accuracy and bias of different methods of estimating expression scores in various simulated cis-genetic architectures.** Left: $R^2$ between predicted and true expression scores at different expression panel sample sizes. Right: Bias of predicted expression scores (slope from regressing predicted expression scores on true expression scores). Default settings: 5 cis-eQTLs per gene, cis-eQTLs randomly selected within 1 Mb window, mean expression cis-heritability = 0.05. (**a-c**) 5, 10, and 1 simulated cis-eQTL per gene respectively. (**d**) eQTLs randomly selected within 10 Kb window. (**e**) Mean expression cis-heritability = 0.01. (**f**) LASSO with REML correction results for various levels of REML noise. Error bars for all plots represent mean standard errors across 100 simulations. See Supplementary Note for remaining details on this simulation.

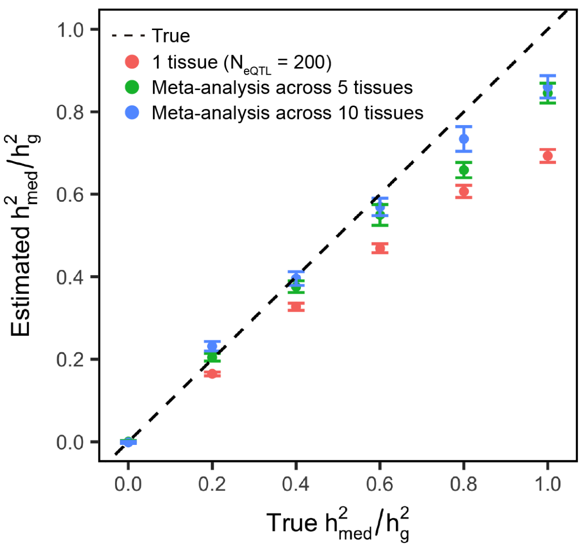


**Supplementary Figure 3.** $h^2_{med}/h^2_g$ **estimates from expression scores meta-analyzed across tissues.** We simulated expression phenotypes in multiple tissues, then estimated expression scores in individual tissues and meta-analyzed expression scores across tissues. Error bars represent mean standard errors across 100 simulations. See Supplementary Note for remaining details on this simulation.

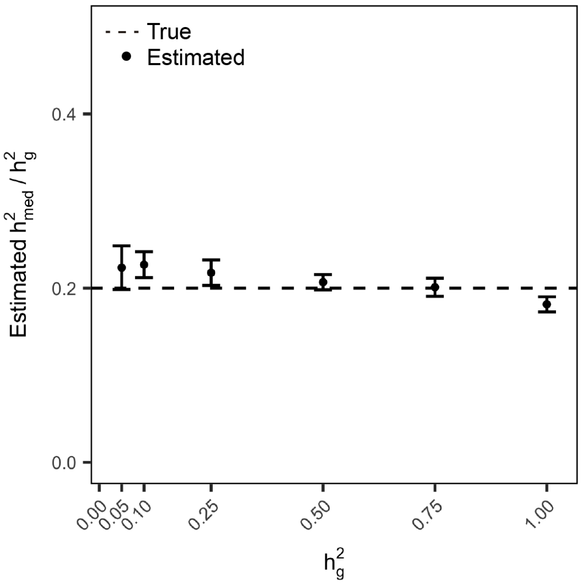


**Supplementary Figure 4.** $h^2_{med}/h^2_g$ **estimates when varying** $h^2_g$**.** Simulation was performed in the same manner as in Figure 2a (with expression panel size fixed at 1000). $h^2_{med}/h^2_g$ was fixed at 0.2 for all simulations. Error bars represent mean standard errors across 100 simulations.

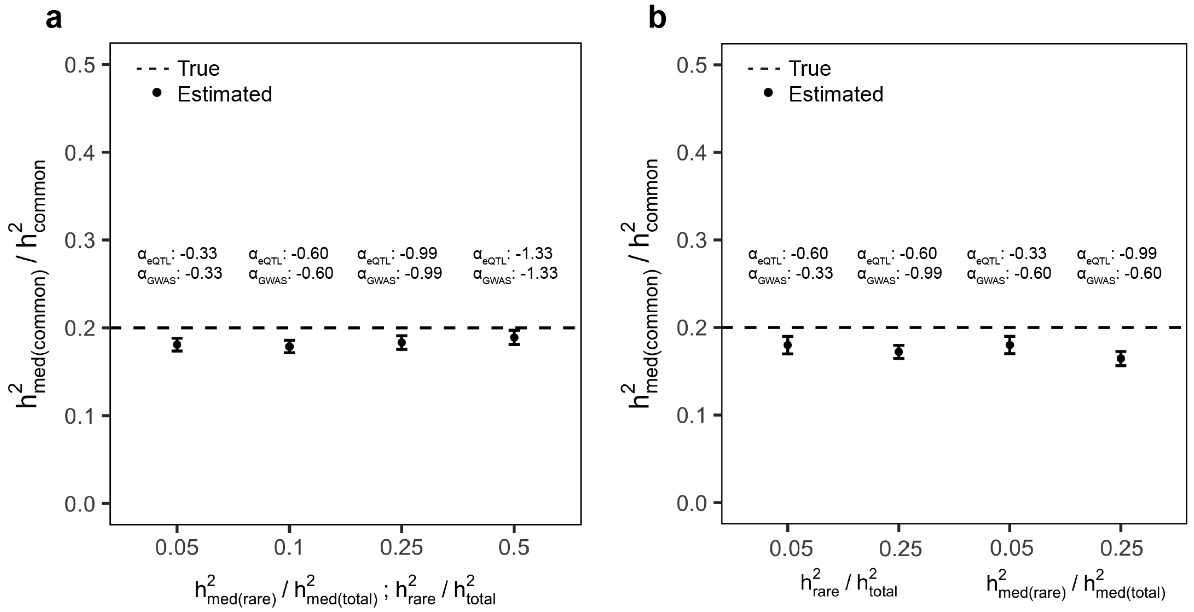


**Supplementary Figure 5. $h^2_{med}/h^2_g$ estimates under simulated frequency-dependent genetic architectures.** Error bars represent mean standard errors across 100 simulations. With $h^2_{med(common)}/h^2_{common}$ fixed at 0.2 and $h^2$ fixed at 0.5, we simulated eQTL effect sizes with variance of per-allele effect sizes proportional to $[p_i(1-p_i)]^{\alpha_{eQTL}}$ and non-mediated effect sizes with variance proportional to $[p_i(1-p_i)]^{\alpha_{GWAS}}$. We selected values of $\alpha_{eQTL}$ and $\alpha_{GWAS}$ to have the following property: $\alpha = -0.33$: 5% of total heritability explained by rare variants with MAF $< 0.01$; $\alpha = -0.60$: 10% heritability explained by rare variants; $\alpha = -0.99$: 25% heritability explained by rare variants; $\alpha = -1.33$: 50% heritability explained by rare variants. (**a**) $h^2_{med(common)}/h^2_{common}$ estimates with $\alpha_{eQTL} = \alpha_{GWAS}$, in which case the proportion of rare $h^2_{med}$ and rare $h^2$ is the same (x-axis). (**b**) $h^2_{med(common)}/h^2_{common}$ estimates with $\alpha_{eQTL} \neq \alpha_{GWAS}$. Left: proportion of rare $h^2_{med}$ is fixed at 0.1, while proportion of rare $h^2$ is varied from 0.05 to 0.25. Right: proportion of rare $h^2$ is fixed at 0.1, while proportion of rare $h^2_{med}$ is varied from 0.05 to 0.25. See Supplementary Note for remaining details on this simulation.

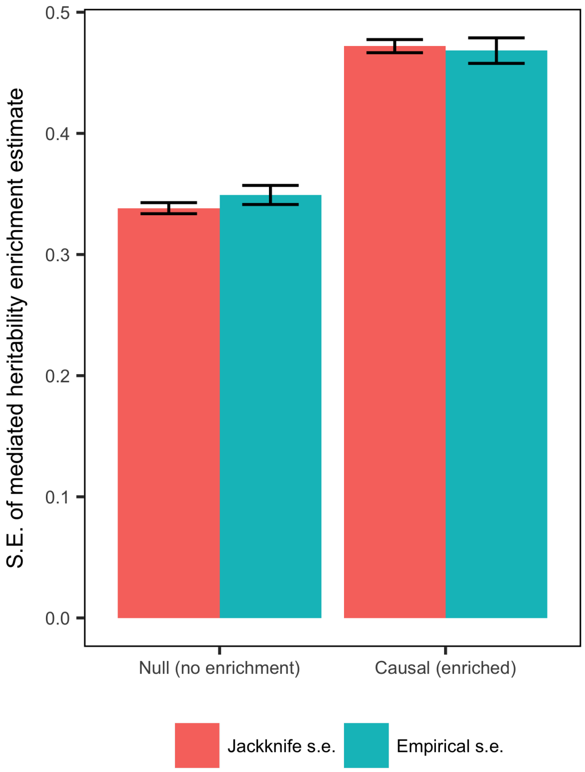


**Supplementary Figure 6. Calibration of jackknife standard errors for $h^2_{med}$ enrichment.** We simulated a gene category without $h^2_{med}$ enrichment (null) and a gene category with 2x $h^2_{med}$ enrichment (causal). Jackknife standard errors for $h^2_{med}$ enrichment are compared to empirical standard errors of $h^2_{med}$ enrichment estimates across 1000 simulations. See Supplementary Note for remaining details on this simulation.

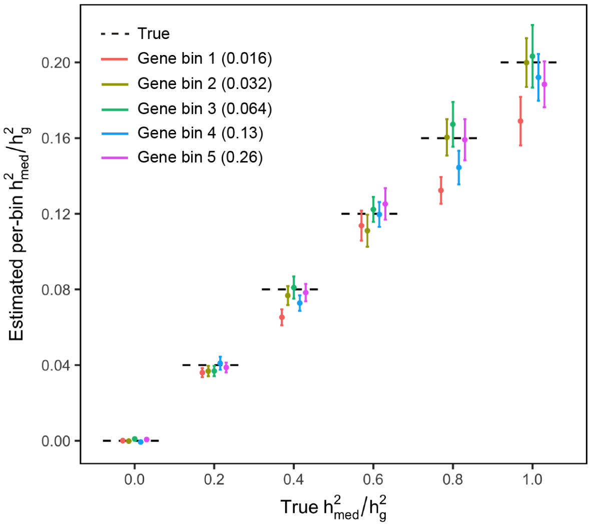


**Supplementary Figure 7. Per-bin $h^2_{med}/h^2_g$ estimates for simulation in Fig 3a.** In the legend, the number in parentheses indicates the average expression cis-heritability of genes in a given gene bin.

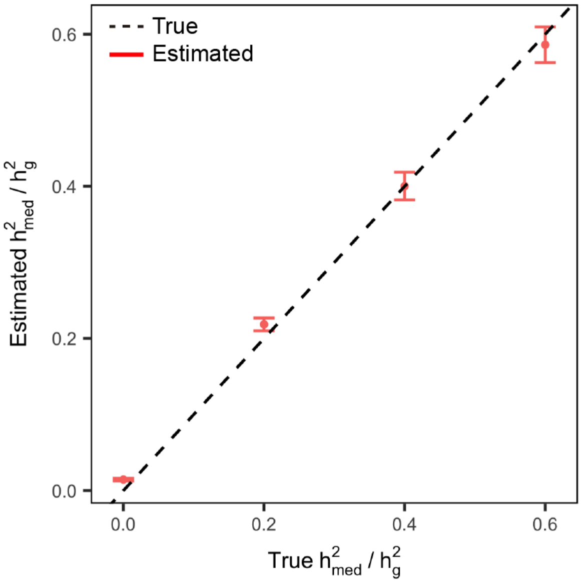


**Supplementary Figure 8.** $h^2_{med}/h^2_g$ **estimates for simulation involving realistic violation to pleiotropy-eQTL independence.** $h^2_{med}/h^2_g$ was varied from 0 to 0.6. Because the union of the three SNP categories with eQTL effect size enrichment (coding regions, conserved regions, and transcription start sites) comprises around 6% of the genome, the maximum value that $h^2_{med}/h^2_g$ can be is 0.6 if we have the condition that $h^2_g$ enrichment of the three SNP categories is 10x. 100 simulations were performed. See Supplementary Note for remaining details on this simulation.

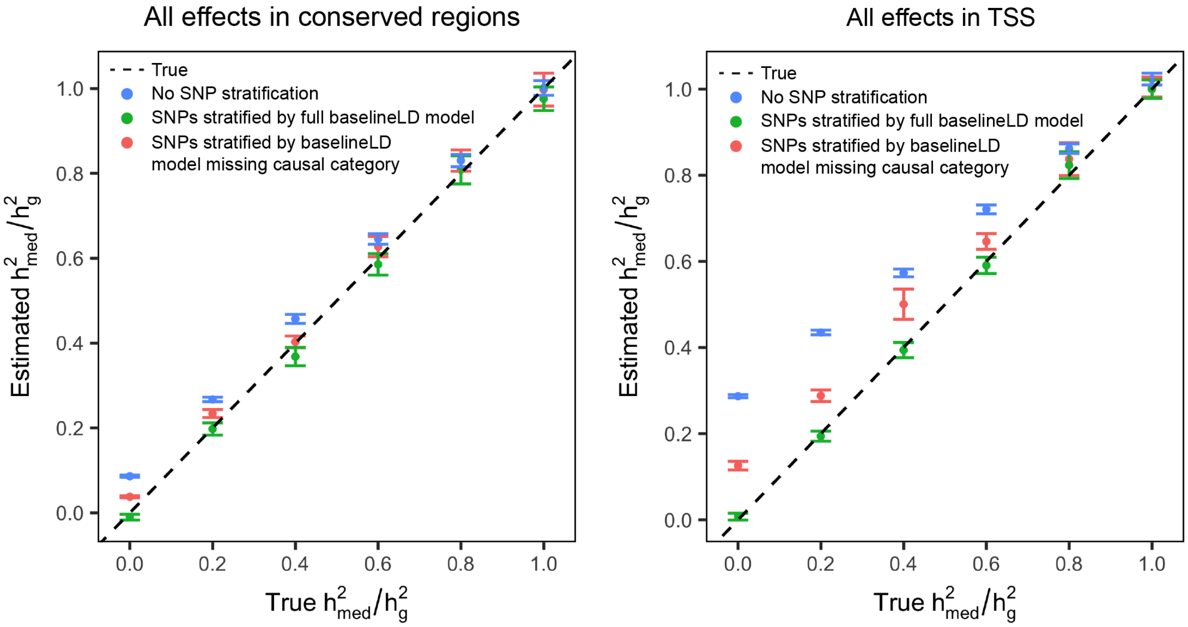


**Supplementary Figure 9.** Same simulation parameters as Figure 3b, but all non-mediated effects and eQTL effects localized in conserved regions and transcription start sites (TSS) respectively.

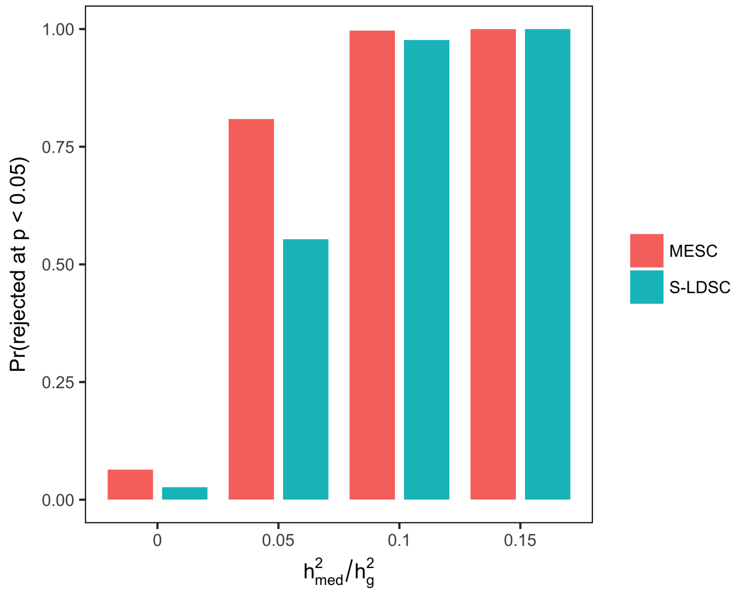


**Supplementary Figure 10. Null calibration and power of MESC and stratified LD-score regression given pleiotropy-eQTL effect size independence.** For various levels of $h^2_{med}/h^2_g$, this figure reports the proportion of simulations which the null hypothesis that $h^2_{med}/h^2_g = 0$ is rejected by MESC, and the proportion of simulations in which the null hypothesis of no $h^2_g$ enrichment for the set of all eQTLs is rejected by stratified LD-score regression (S-LDSC). All effect sizes, expression phenotypes, and complex trait phenotypes were simulated in the same manner as Figure 2a. For stratified LD-score regression, we defined the eQTL category as the set of all true eQTLs. 300 simulations were performed.

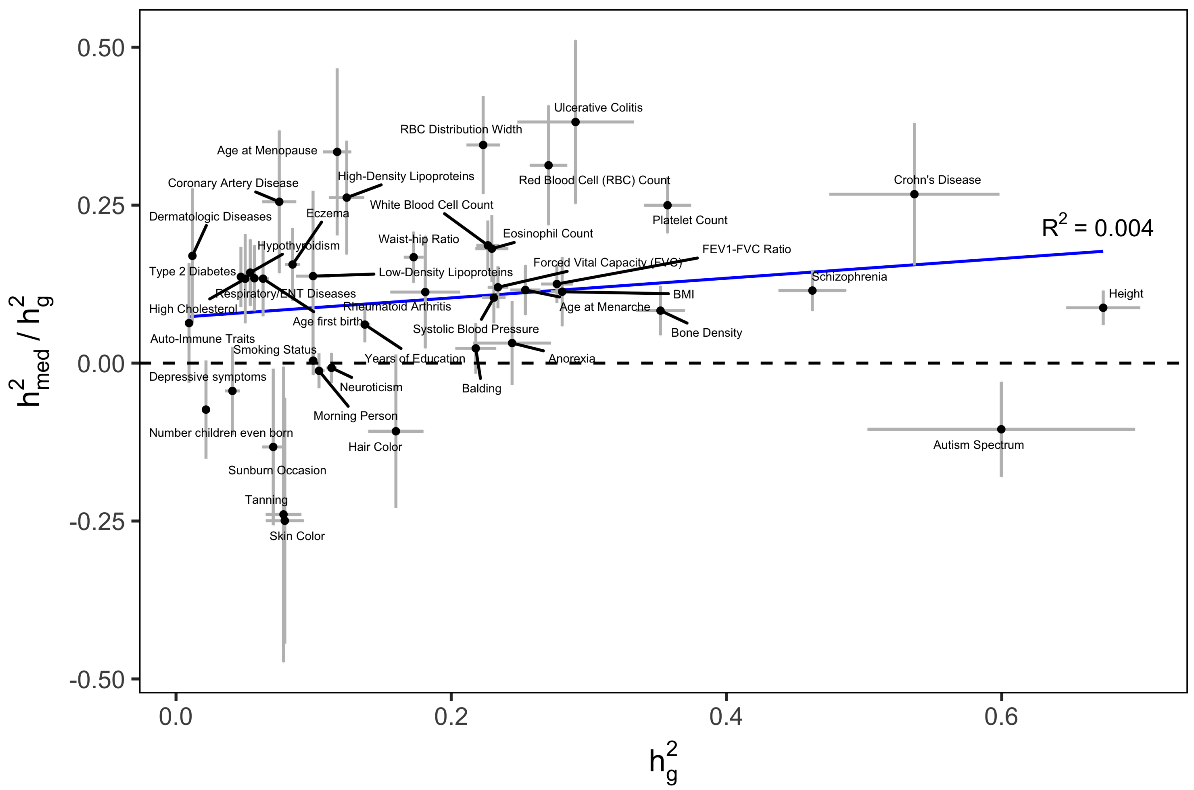


**Supplementary Figure 11. Relationship between $h^2_{med}/h^2_g$ and $h^2_g$.** $h^2_{med}/h^2_g$ estimates were obtained using all-tissue meta-analyzed expression scores. $h^2_g$ estimates were obtained using stratified LD-score regression. Error bars represent jackknife standard errors.

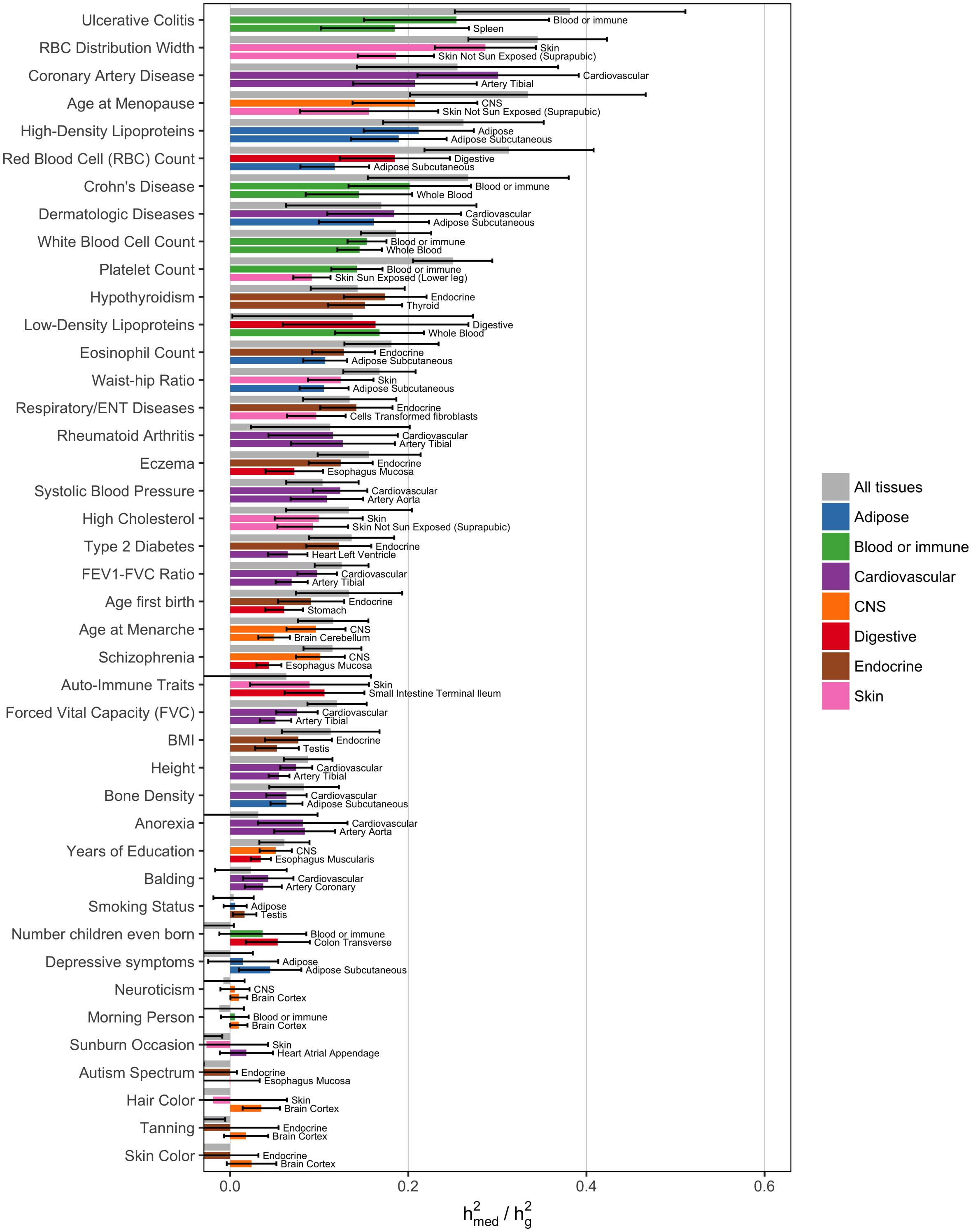


**Supplementary Figure 12.** Same as Figure 5a, but containing $h^2_{med}/h^2_g$ estimates for all 42 traits from all three types of expression scores: "All tissues" (expression scores meta-analyzed across all 48 GTEx tissues), "Best tissue group" (expression scores meta-analyzed within 7 tissue groups), and "Best tissue" (expression scores computed within individual tissues). Here, "best" refers to the tissue/tissue group resulting in the highest estimates of $h^2_{med}/h^2_g$ compared to all other tissues/tissue groups.

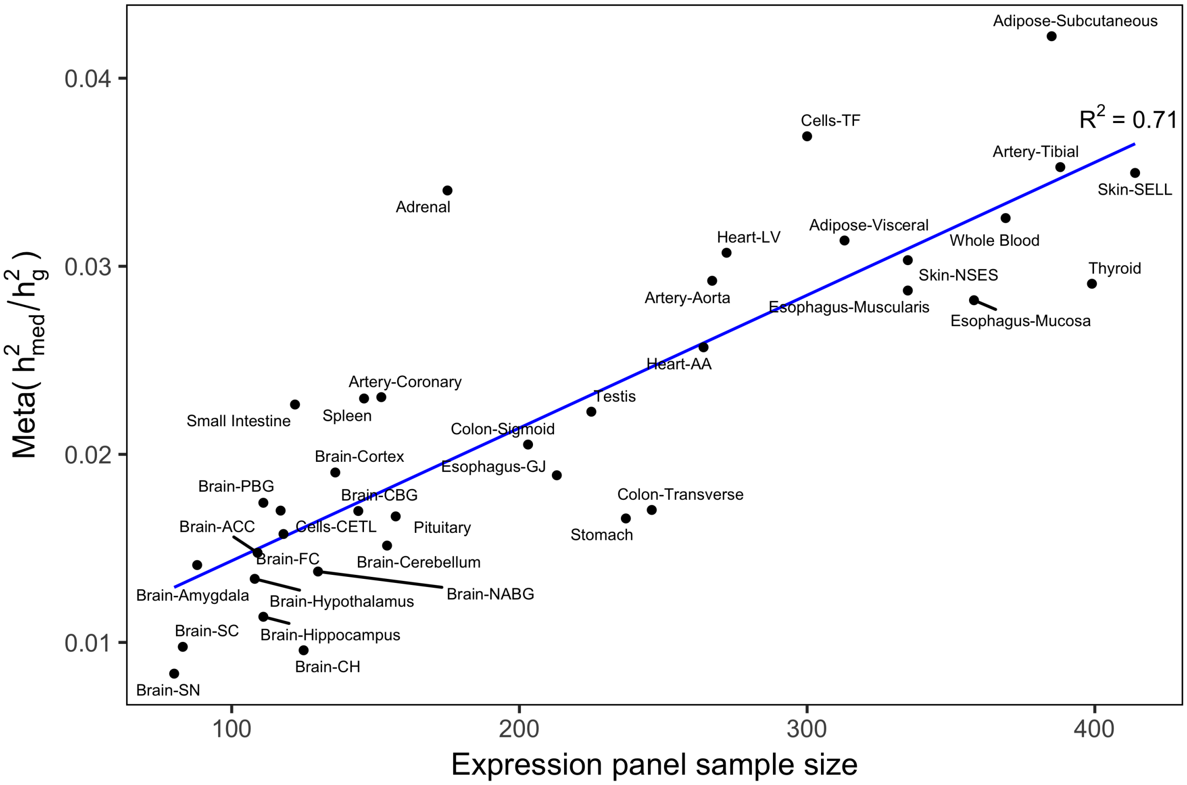


**Supplementary Figure 13. Relationship between individual tissue sample size and magnitude of $h^2_{med}/h^2_g$.** $h^2_{med}/h^2_g$ estimates from expression scores estimated in each of 48 individual GTEx tissues were meta-analyzed across 42 complex traits, then plotted against the number of samples in each tissue. We use the following abbreviations: adipose visceral, adipose visceral omentum; brain ACC, brain anterior cingulate cortex BA24; brain CBG, brain caudate basal ganglia; brain CH, brain cerebellar hemisphere; brain FC, brain frontal cortex BA9; brain NABG, brain nucleus accumbens basal ganglia; brain PBG brain putamen basal ganglia; cells CETL, cells EBV-transformed lymphocytes; cells TF, cells transformed fibroblasts; esophagus–GJ, esophagus gastroesophageal junction; heart AA, heart atrial appendage; heart LV, heart left ventricle; skin NSES, skin not sun exposed suprapubic; skin SELL, skin sun exposed lower leg; small intestine, small intestine terminal ileum.

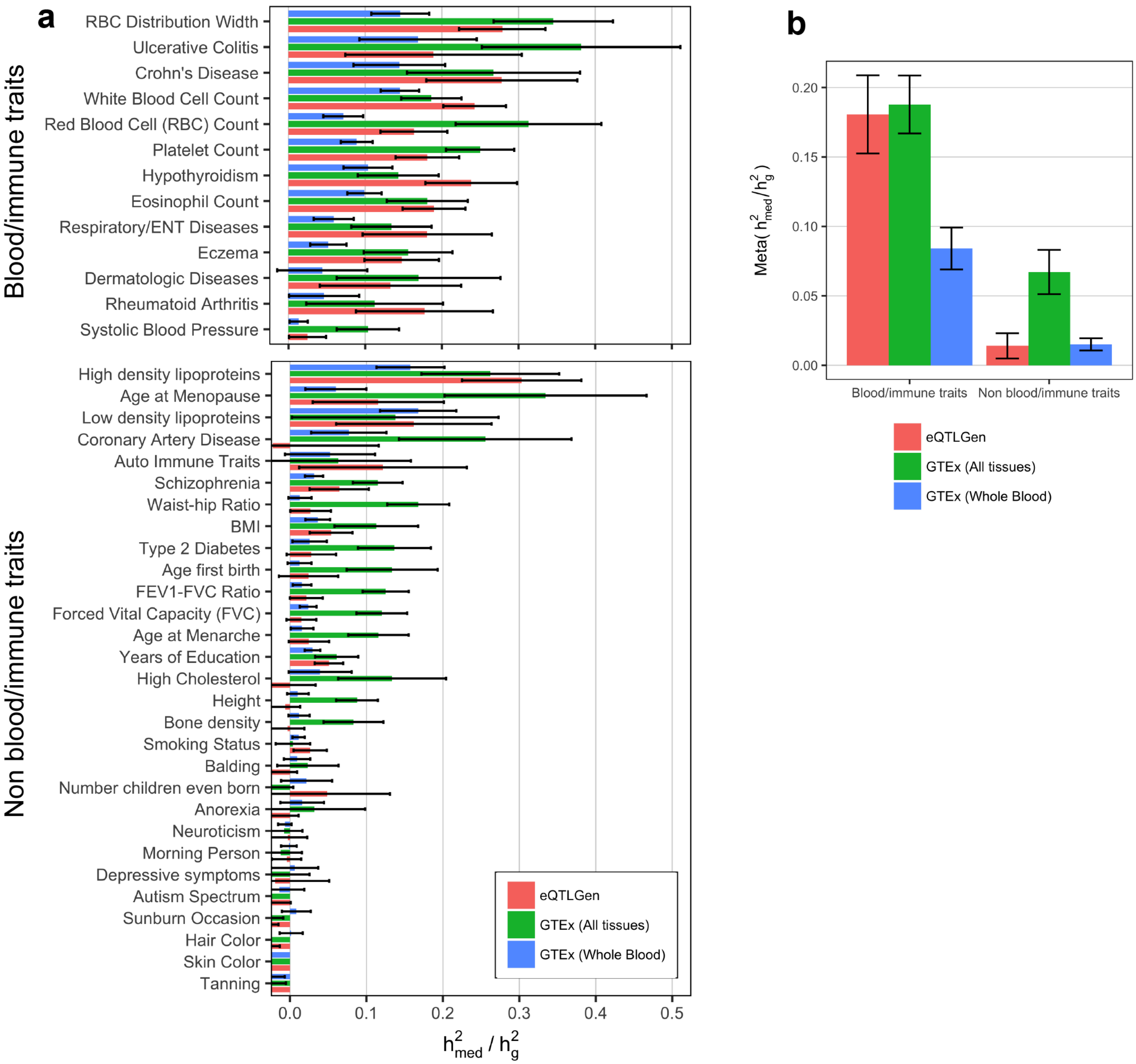


**Supplementary Figure 14. $h^2_{med}/h^2_g$ estimates for 42 diseases and complex traits using data from eQTLGen.** We estimated expression scores for all SNPs using cis-eQTL summary statistics from eQTLGen ($N$ = 31,684 blood samples), then estimated $h^2_{med}/h^2_g$ using GWAS summary statistics for the same 42 traits analyzed in the main text. In line with our previous analyses using GTEx data, we stratified expression scores by 5 expression cis-heritability bins. Expression cis-heritability estimates for eQTLGen data were obtained using LD-score regression[56]. For sake of comparison, we also display $h^2_{med}/h^2_g$ estimates obtained from expression scores from GTEx all-tissue meta-analysis and GTEx whole blood only. (**a**) $h^2_{med}/h^2_g$ estimates for 42 individual traits, organized into blood/immune and non-blood/immune traits. (**b**) Results from **a** meta-analyzed across traits. Note that low estimates of $h^2_{med}/h^2_g$ for GTEx whole blood expression scores are caused by the small sample size of the GTEx whole blood data set ($N$ = 369).

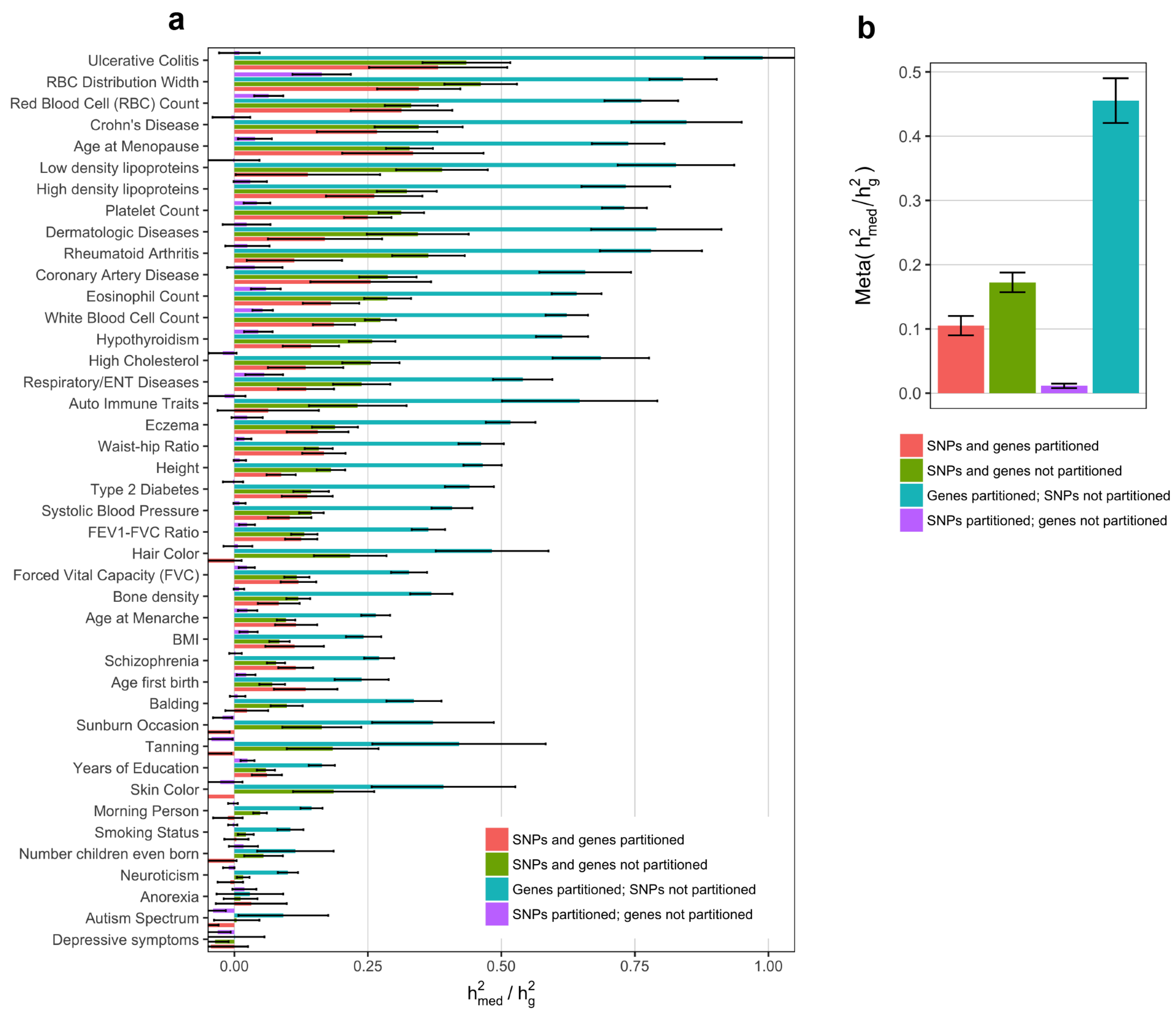


**Supplementary Figure 15.** $h^2_{med}/h^2_g$ **estimates without stratifying genes/SNPs.** All estimates were obtained using all-tissue meta-analyzed expression scores. (**a**) We estimated $h^2_{med}/h^2_g$ for 42 traits without stratifying genes by 5 expression cis-heritability and/or without stratifying SNPs by the baselineLD model. (**b**) Estimates from **a** meta-analyzed across all 42 traits.

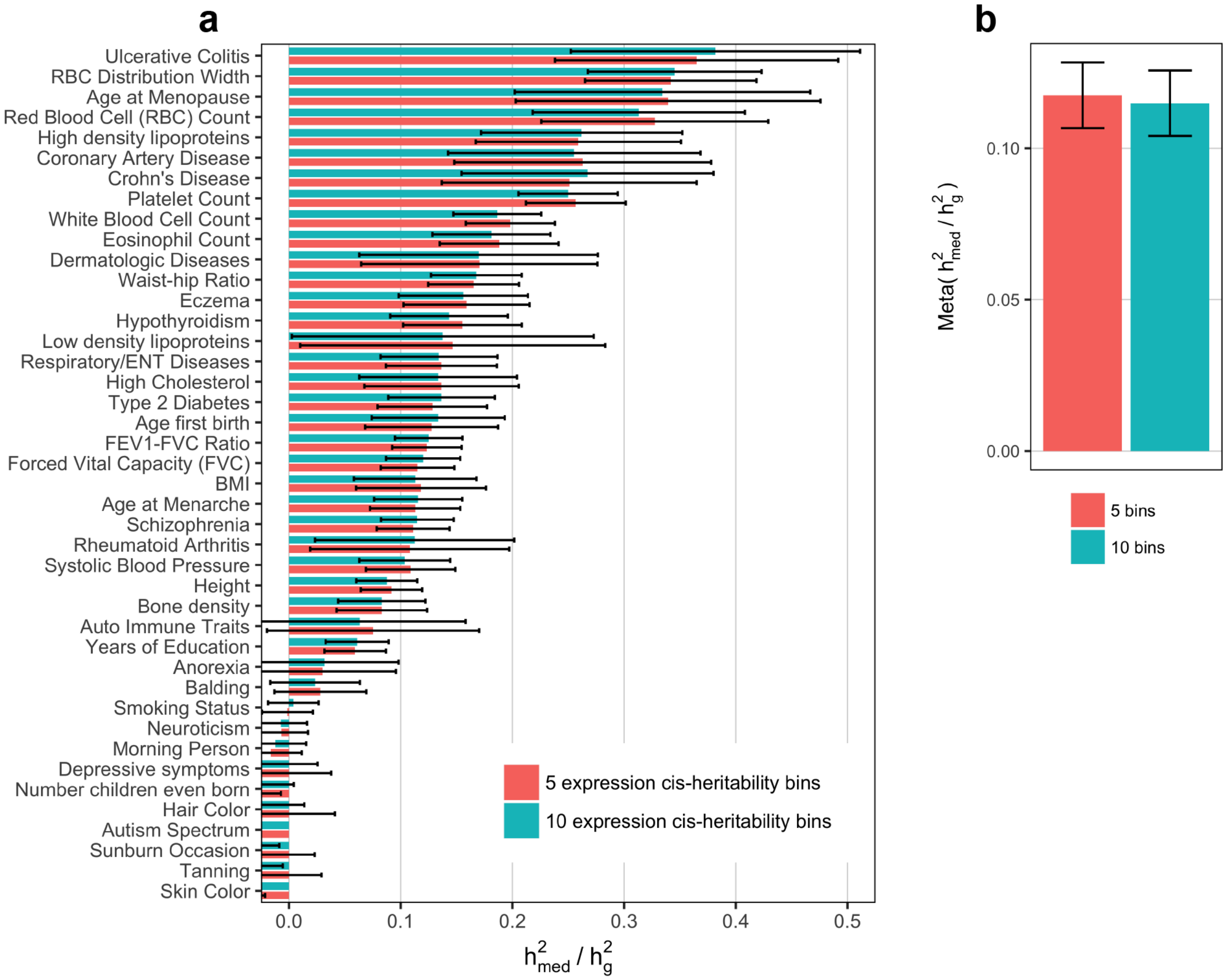


**Supplementary Figure 16.** $h^2_{med}/h^2_g$ **estimates when varying number of expression cis-heritability bins**. All estimates were obtained using all-tissue meta-analyzed expression scores. (**a**) We estimated $h^2_{med}/h^2_g$ for 42 traits while stratifying genes by either 5 or 10 expression cis-heritability bins. (**b**) Estimates from **a** meta-analyzed across all 42 traits.

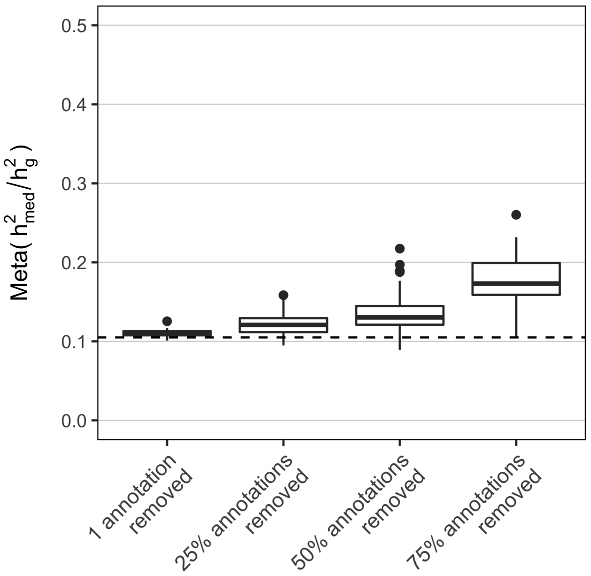


**Supplementary Figure 17.** $h^2_{med}/h^2_g$ **estimates when removing subsets of the baselineLD model.** In total, the baselineLD model v2.0 we used in our main analyses contains 72 SNP annotations. We grouped together related baselineLD annotations (typically consisting of a main annotation and the same annotation with 100-500 bp flanking windows), producing the following 29 categories: Coding, Conserved, CTCF, DGF, DHS, Enhancer, Fetal DHS, H3K27ac, H3K4me1, H3K4me3, H3K9ac, Intron, Promoter, Repressed, Super Enhancer, TFBS, Transcribed, TSS, 3' UTR, 5' UTR, Weak Enhancer, GERP, Allele Age, LLD, Recombination Rate, Nucleotide Diversity, Background Selection Statistic, CpG Content, and ASMC. When removing annotations, we remove all related annotations that fall into one of the 29 categories. Each data point represents an $h^2_{med}/h^2_g$ estimate meta-analyzed over 42 traits and estimated from GTEx all-tissue expression scores. For the boxplot labelled "1 annotation removed," we show $h^2_{med}/h^2_g$ estimates when removing each of the 29 individual categories. For the boxplots labelled "25%/50%/75% annotations removed," we show $h^2_{med}/h^2_g$ estimates from 100 random subsets of the categories corresponding to the percentage of annotations removed. Dotted line indicates the $h^2_{med}/h^2_g$ estimate when using the full baselineLD model.

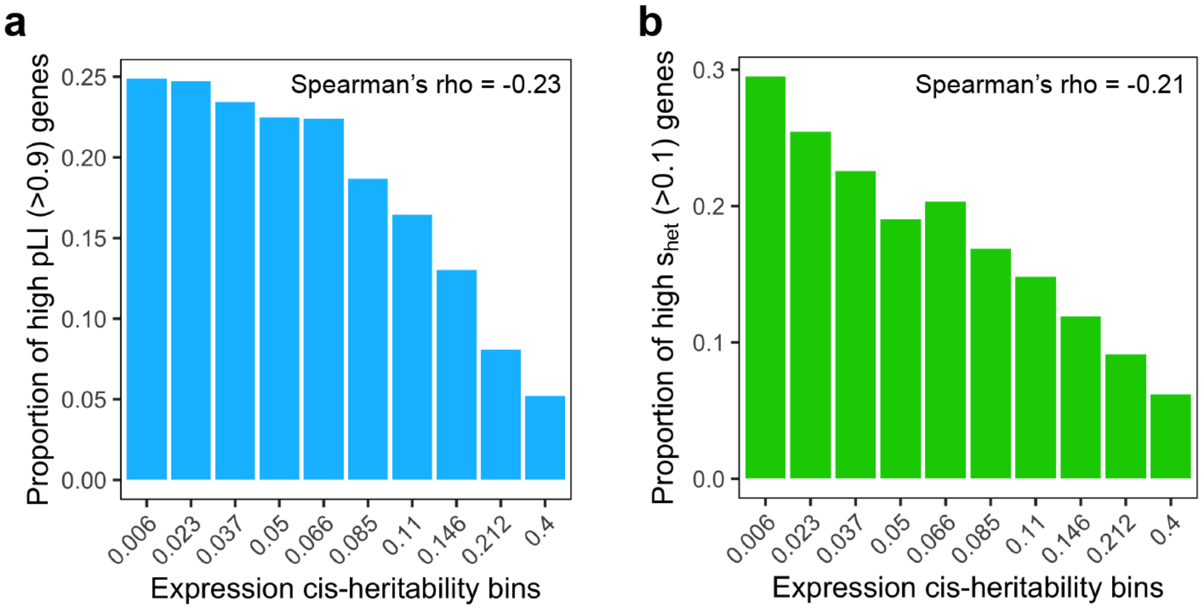


**Supplementary Figure 18. Relationship between expression cis-heritability and metrics of gene essentiality.** For each gene, pLI (probability of loss-of-function intolerance) was obtained from Lek et al. 2016 Nature and $s_{het}$ (selection against protein-truncating variants) was obtained from Cassa et al. 2017 Nature Genetics.

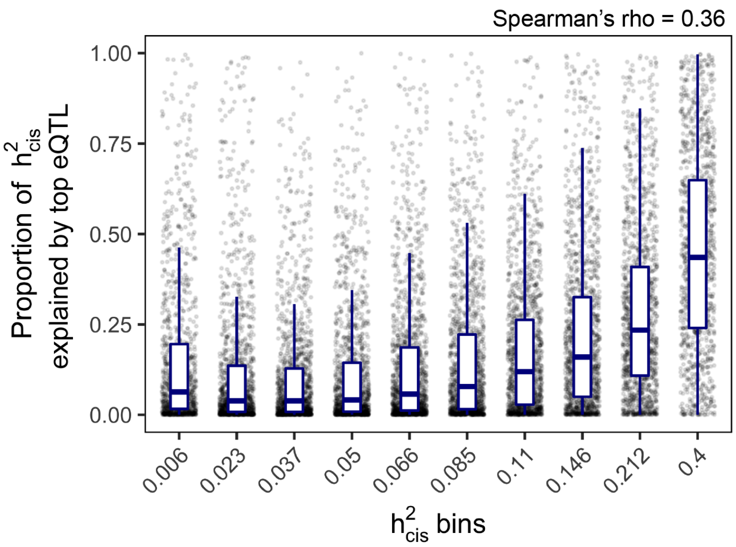


**Supplementary Figure 19. Relationship between expression cis-heritability and proportion of expression cis-heritability explained by top cis-eQTL.** Top cis-eQTL effect size is meta-analyzed using fixed effects meta-analysis across all 48 GTEx tissues.

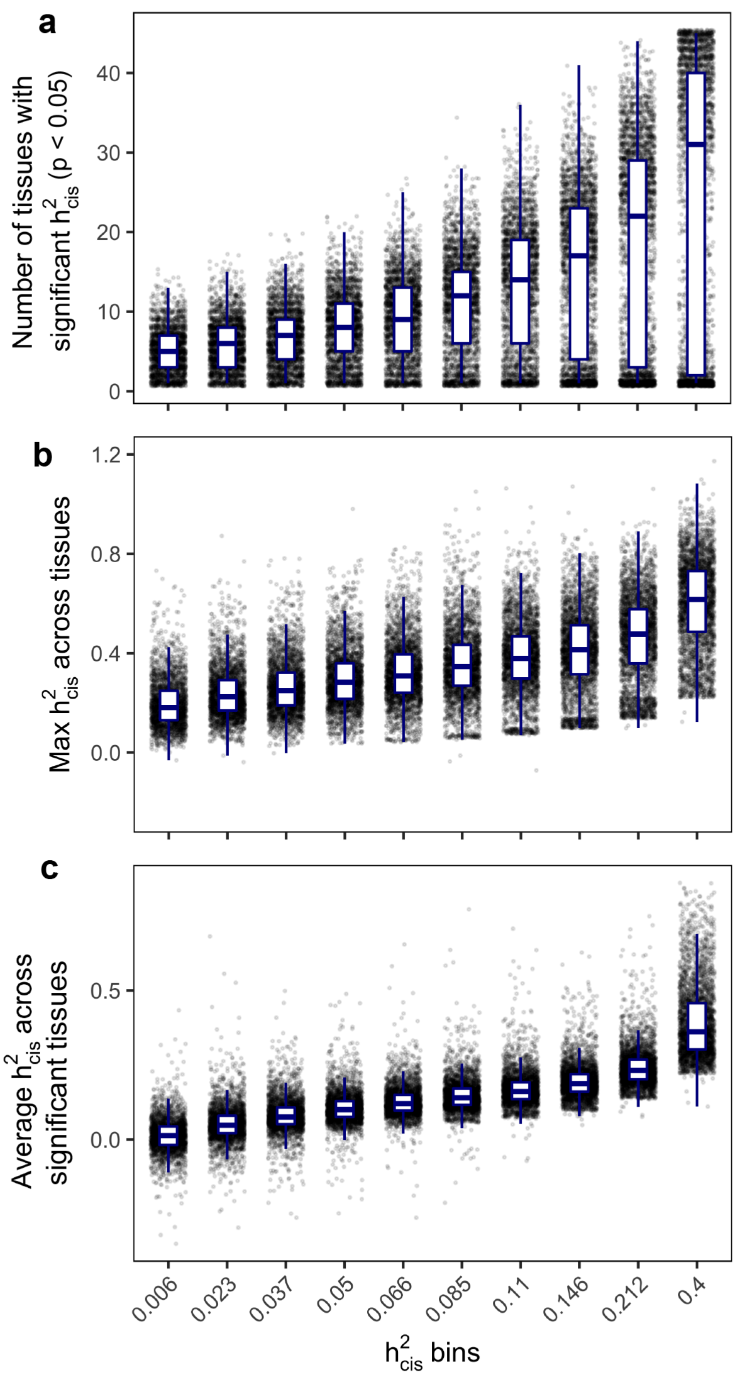


**Supplementary Figure 20. Relationship between expression cis-heritability and metrics of tissue specificity.** Meta-tissue $h^2_{cis}$ (x-axis) is computed for each gene by averaging $h^2_{cis}$ across all tissues. x-axis labels indicate the average meta-tissue $h^2_{cis}$ of genes within each decile. $h^2_{cis}$ (y-axis) refers to estimates within individual tissues. (**a**) Relationship between meta-tissue $h^2_{cis}$ deciles and the number of tissues with significantly nonzero $h^2_{cis}$ ($p < 0.05$) for each gene. (**b**) Relationship between meta-tissue $h^2_{cis}$ deciles and the max $h^2_{cis}$ across tissues for each gene. (**c**) Relationship between meta-tissue $h^2_{cis}$ deciles and the average $h^2_{cis}$ across tissues with significantly nonzero $h^2_{cis}$ ($p < 0.05$) for each gene.

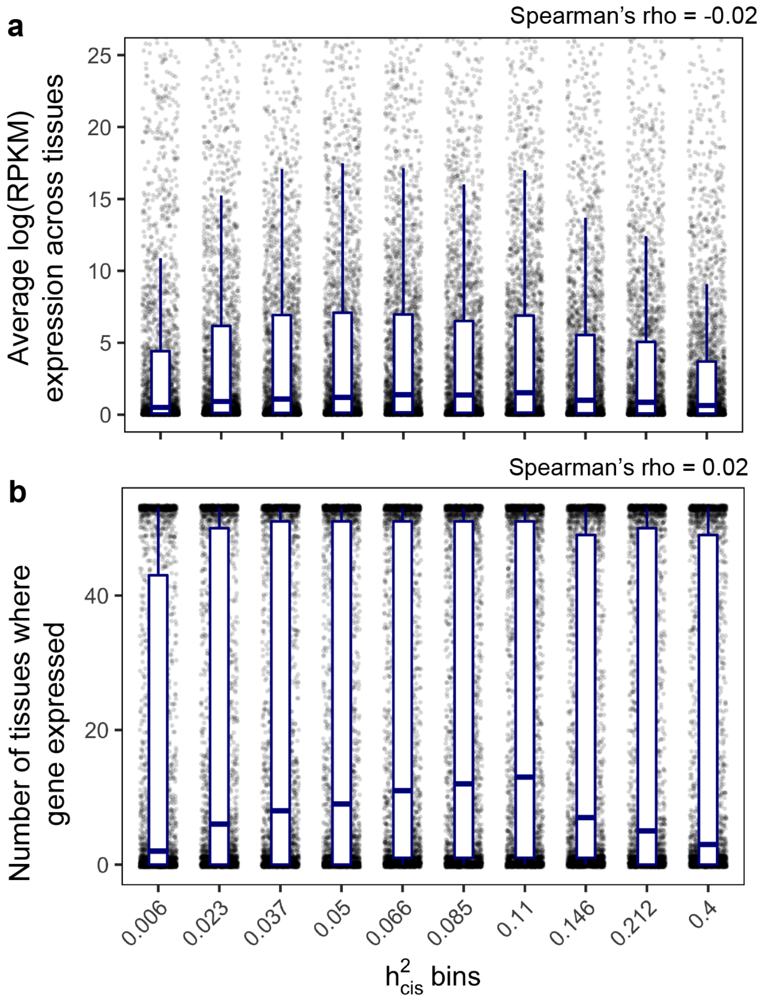


**Supplementary Figure 21. Relationship between expression cis-heritability and expression levels.** (**a**) Expression levels represent the median log(RPKM) expression across individuals, which are then averaged across tissues. (**b**) A gene is expressed in a tissue if RPKM > 0.3 in that tissue.

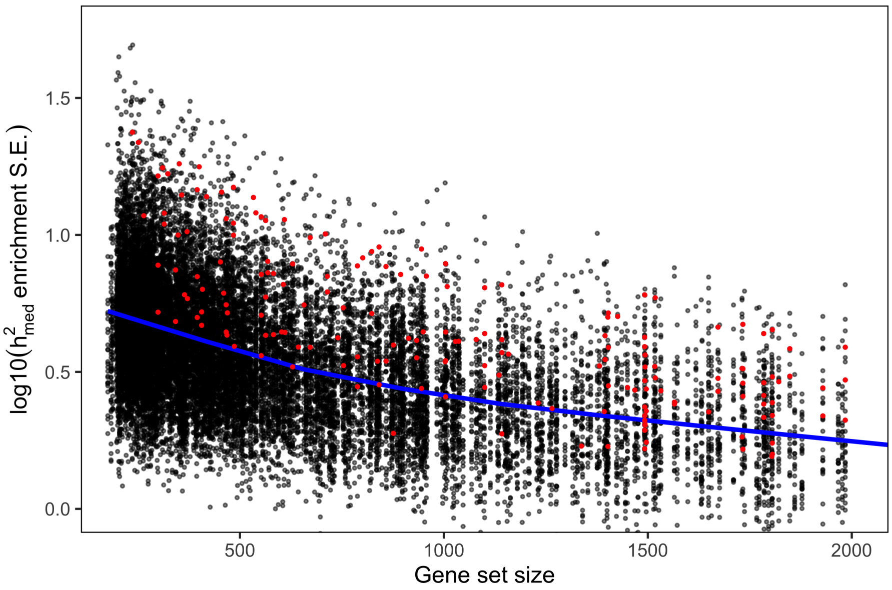


**Supplementary Figure 22. Relationship between gene set size and log10 $h^2_{med}$ enrichment standard error.** Each point represent a gene set-complex trait pair. Points highlighted in red indicate gene set-complex trait pairs with FDR $<$ 0.05 (after accounting for 21,502 hypotheses tested). Blue line indicates the LOESS best fit line.

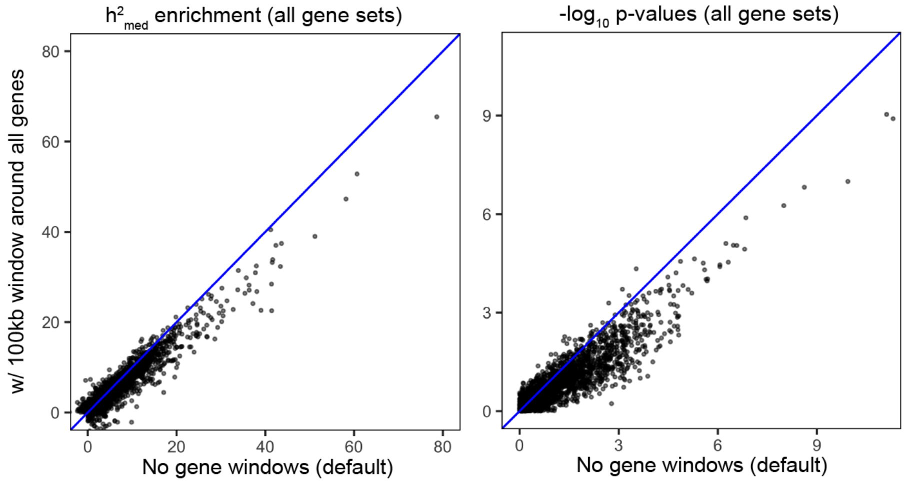


**Supplementary Figure 23. $h^2_{med}$ enrichment estimates with 100 Kb window around genes.** (**Left**) $h^2_{med}$ enrichment estimates for all 21,502 trait-gene sets pairs analyzed in the main text when including a SNP annotation corresponding to 100 Kb windows around each gene in each gene set. (**Right**) Same as left, but showing $h^2_{med}$ enrichment p-values.

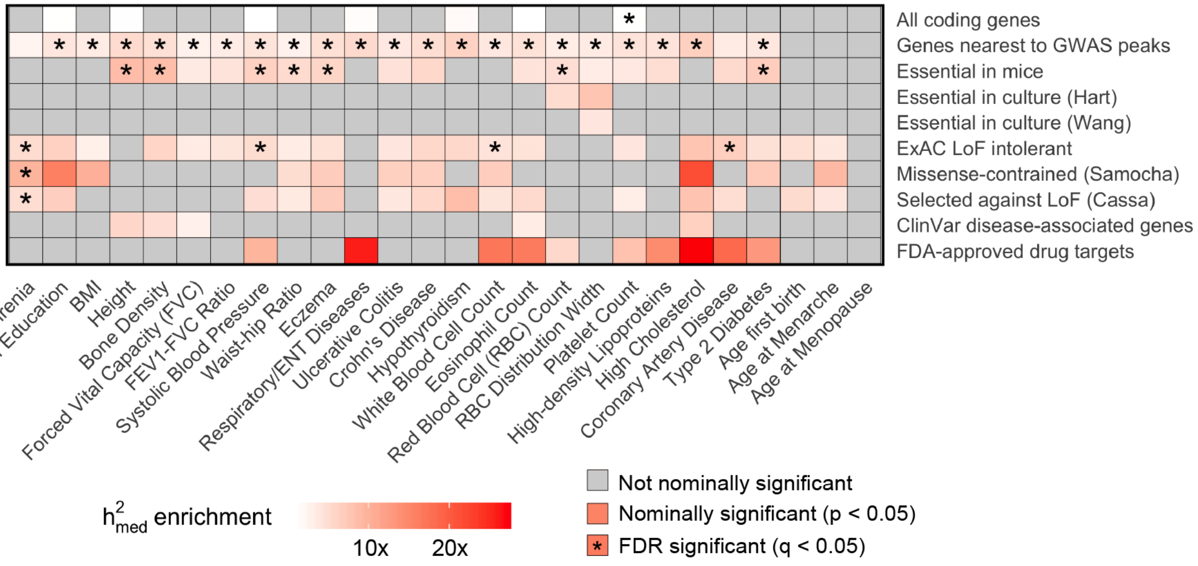


**Supplementary Figure 24.** $h^2_{med}$ **enrichment estimates for all 10 broadly essential gene sets across all 26 complex traits.**

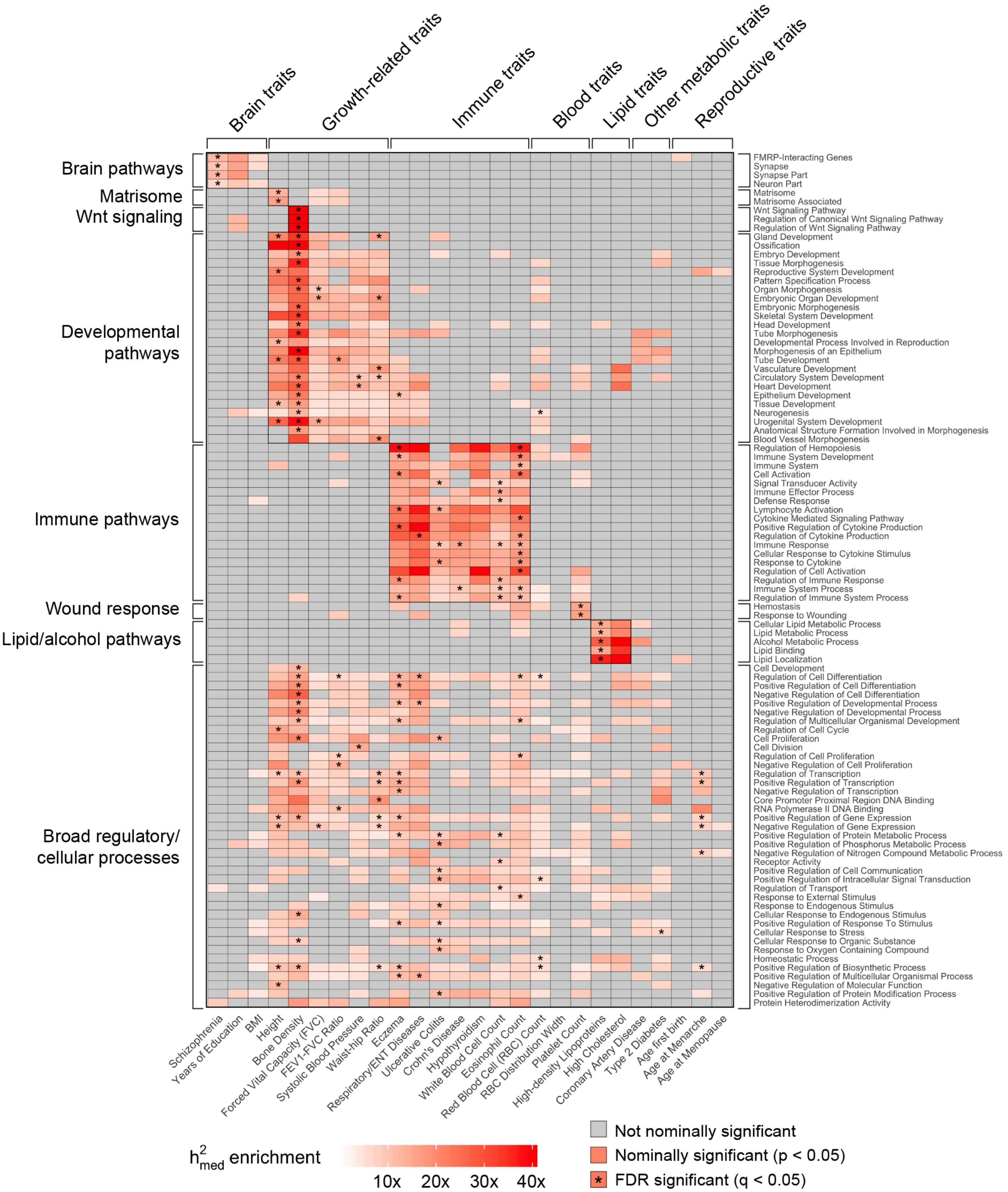


**Supplementary Figure 25. $h^2_{med}$ enrichment estimates for 97 pathway-specific gene sets across all 26 complex traits.** 97 pathway-specific gene sets represent all pathway-specific gene sets (out of 780 total) with FDR-significant $h^2_{med}$ enrichment in at least one of the 26 complex traits. For sake of display, we grouped together related traits and gene sets.

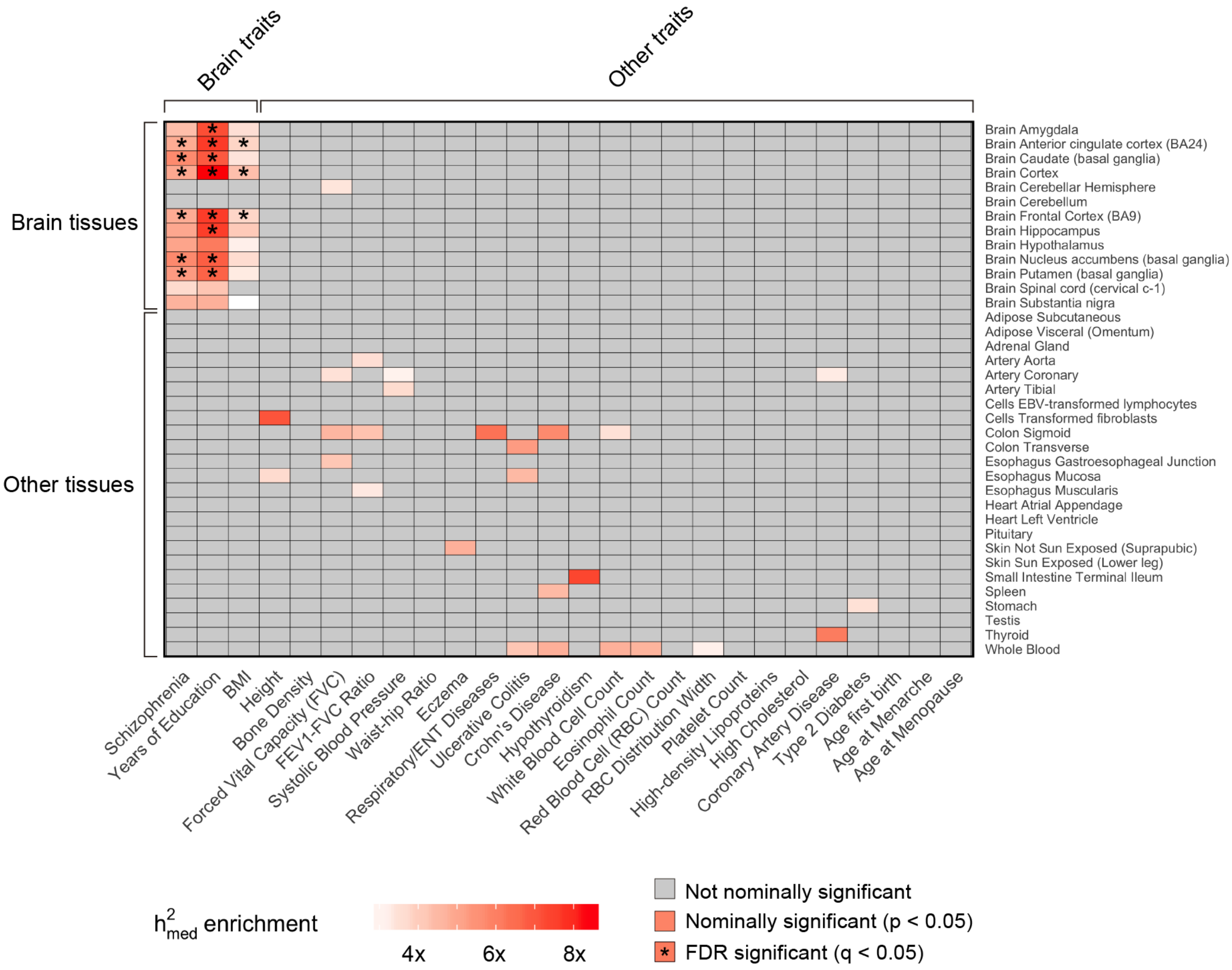


**Supplementary Figure 26.** Same as Figure 7b, but including $h^2_{med}$ enrichment estimates for all 26 traits and with individual GTEx tissues labelled.

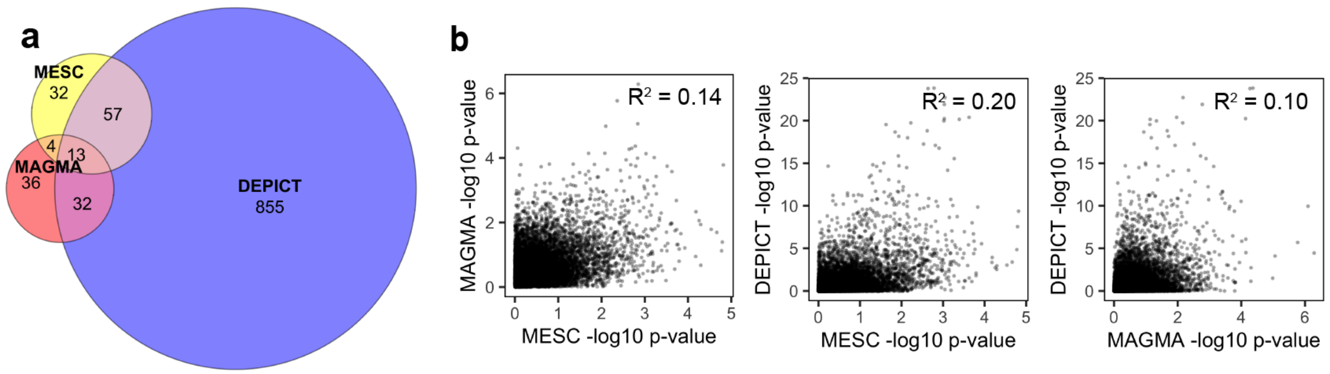


**c**

| Trait | Gene set name | MESC p-value | MAGMA p-value | DEPICT p-value |
|---|---|---|---|---|
| BMI | Regulation of Cellular Component Biogenesis (GO) | 0.0016 | 0.22 | 0.21 |
| Bone Density | Forebrain Development (GO) | 0.00091 | 0.66 | 0.67 |
| | Blood Vessel Morphogenesis (GO) | 0.0016 | 0.051 | 0.11 |
| | Cellular Response to Endogenous Stimulus (GO) | 0.00072 | 0.016 | 0.0049 |
| | Urogenital System Development (GO) | 0.00073 | 0.0059 | 0.061 |
| | Vasculature Development (GO) | 0.0013 | 0.22 | 0.12 |
| | Pattern Specification Process (GO) | 5.1E-05 | 0.012 | 0.0029 |
| | Regionalization (GO) | 0.0011 | 0.019 | 0.0029 |
| | Core Promoter Proximal Region DNA Binding (GO) | 0.0011 | 0.38 | 0.42 |
| Crohn's Disease | Positive Regulation of Phosphorus Metabolic Process (GO) | 0.0011 | 0.03 | 0.0044 |
| Eczema | Epithelium Development (GO) | 0.00068 | 0.65 | 0.17 |
| | Negative Regulation of Cell Differentiation (GO) | 0.0014 | 0.0022 | 0.038 |
| FEV1-FVC Ratio | Negative Regulation of Cell Proliferation (GO) | 0.0002 | 0.28 | 0.078 |
| | Positive Regulation of Cell Differentiation (GO) | 0.0014 | 0.055 | 0.0028 |
| Forced Vital Capacity | Sensory Organ Development (GO) | 0.0011 | 0.21 | 0.087 |
| Height | Cell Cycle (Reactome) | 0.0013 | 0.0044 | 0.028 |
| | Urogenital System Development (GO) | 5.6E-05 | 0.0087 | 0.0019 |
| High-density Lipoproteins | Phospholipid Metabolic Process (GO) | 0.0013 | 0.49 | 0.061 |
| Respiratory/ENT Diseases | Positive Regulation of Multicellular Organismal Process (GO) | 0.00016 | 0.16 | 0.016 |
| | Regulation of Cell Adhesion (GO) | 0.00091 | 0.025 | 0.013 |
| Schizophrenia | Synapse (GO) | 0.00075 | 0.095 | 0.023 |
| | Synapse Part (GO) | 0.00083 | 0.1 | 0.034 |
| Systolic Blood Pressure | Cell Division (GO) | 0.00064 | 0.21 | 0.84 |
| Ulcerative Colitis | Positive Regulation of Phosphorus Metabolic Process (GO) | 0.00016 | 0.067 | 0.0081 |
| | Positive Regulation of Protein Metabolic Process (GO) | 0.00073 | 0.094 | 0.0028 |
| | Positive Regulation of Protein Modification Process (GO) | 0.00029 | 0.045 | 0.066 |
| Waist-hip Ratio | Gland Development (GO) | 0.00042 | 0.22 | 0.077 |
| | Core Promoter Proximal Region DNA Binding (GO) | 0.00032 | 0.63 | 0.41 |
| | Double Stranded DNA Binding (GO) | 0.0014 | 0.42 | 0.11 |
| | Protein Heterodimerization Activity (GO) | 0.00086 | 0.32 | 0.0061 |
| White Blood Cell Count | Regulation of Vesicle Mediated Transport (GO) | 0.0015 | 0.28 | 0.062 |
| | Response to Cytokine (GO) | 0.0012 | 0.12 | 0.0028 |

**Supplementary Figure 27. Comparison between gene set enrichment estimates from MESC, MAGMA, and DEPICT.** (**a**) Venn diagram showing the overlap between significantly enriched trait-gene set pairs (FDR $< 0.05$) identified by the three methods. (**b**) Scatterplots of -log10 enrichment p-values from MESC vs. MAGMA (left), MESC vs. DEPICT (middle), and MAGMA vs. DEPICT (right). Each point represents a trait-gene set pair. (**c**) List of all 32 gene sets-complex traits pairs detected as significant by MESC (FDR q-value $< 0.05$) that are not detected as significant by MAGMA or DEPICT. See Supplementary Table 9 for enrichment estimates for all gene set-complex traits pairs.

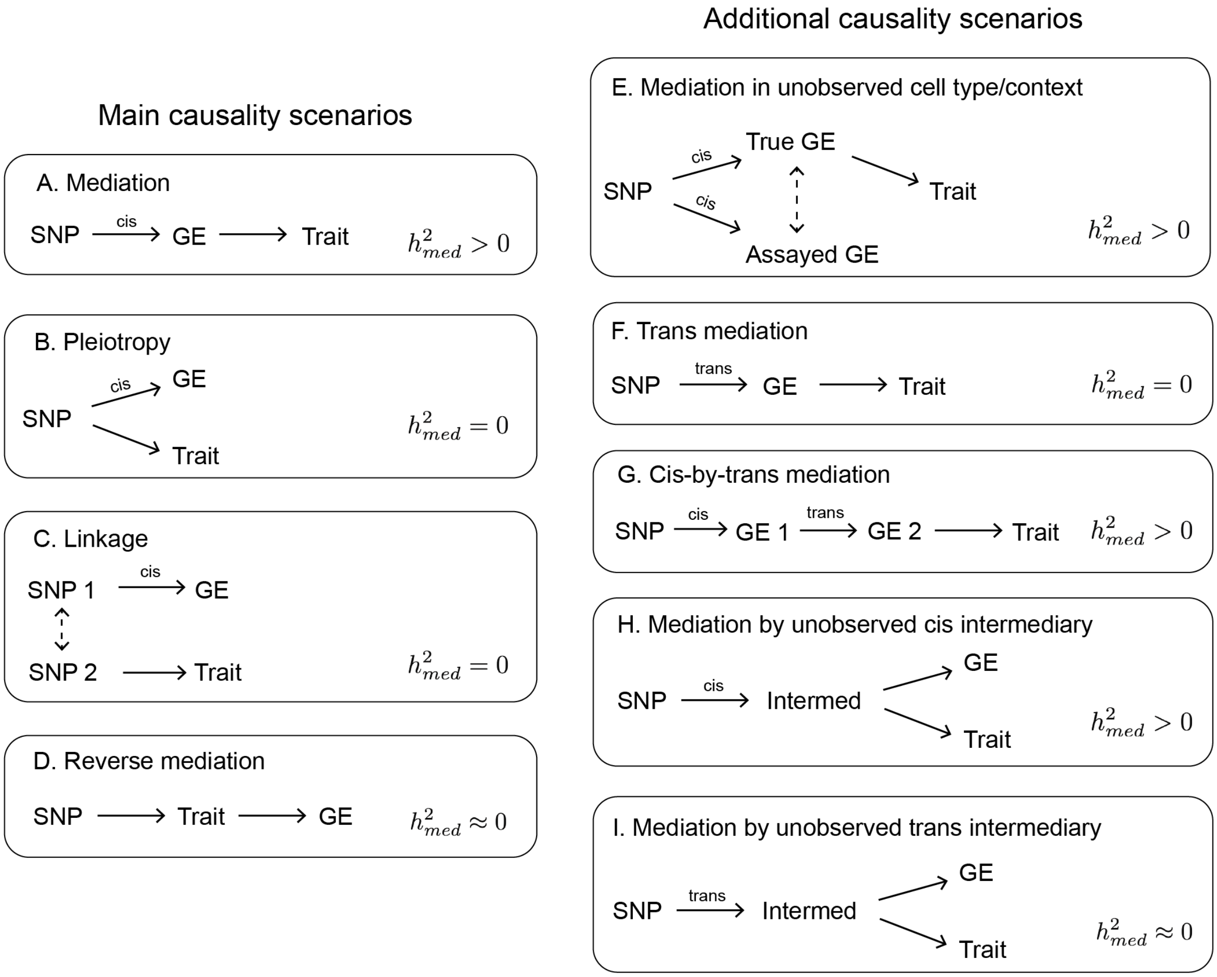


**Supplementary Figure 28. Modes of expression causality.** See "Modes of expression causality" for a description of each scenario and its contribution to estimates of $h^2_{med}$.

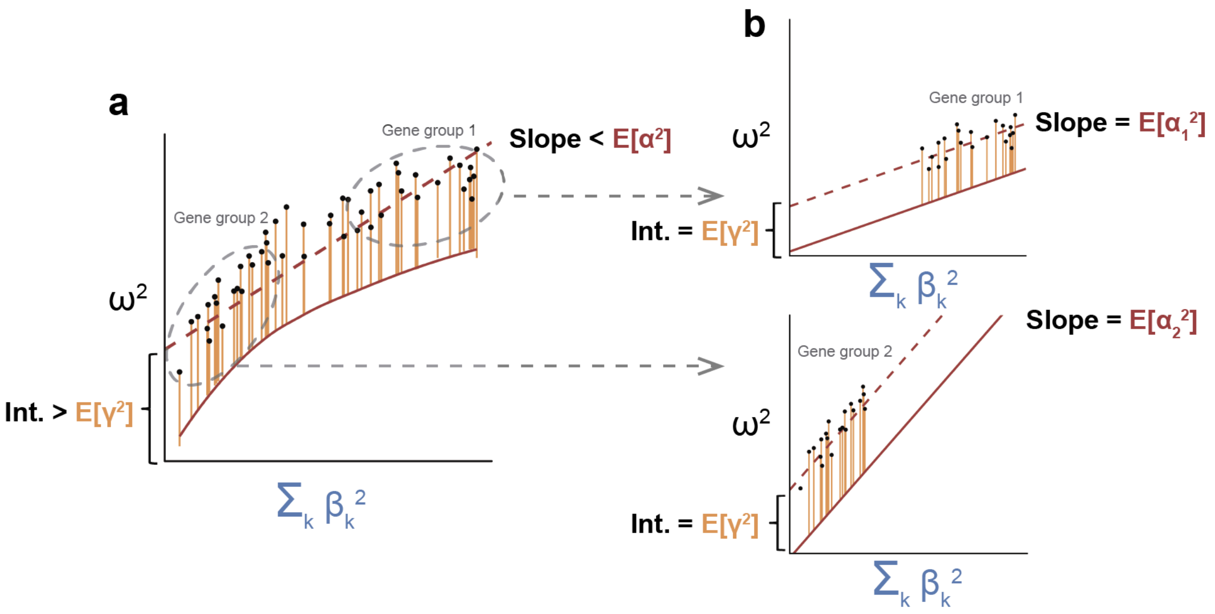


**Supplementary Figure 29. Illustration of impact of violations to gene-eQTL effect size independence on estimates of $E[\alpha^2]$.** See "Model assumptions" for context for this figure. In the figure, we depict a scenario where the magnitude of $\alpha$ is negatively correlated with the magnitude of $\beta$. (**a**) If we perform the regression using all genes, the slope from the regression will be downwardly biased relative to the true $E[\alpha^2]$. (**b**) If we stratify the regression across genes by the magnitude of their expression cis-heritability, we can obtain approximately unbiased estimates of $E[\alpha_D^2]$ for each gene category $D$.

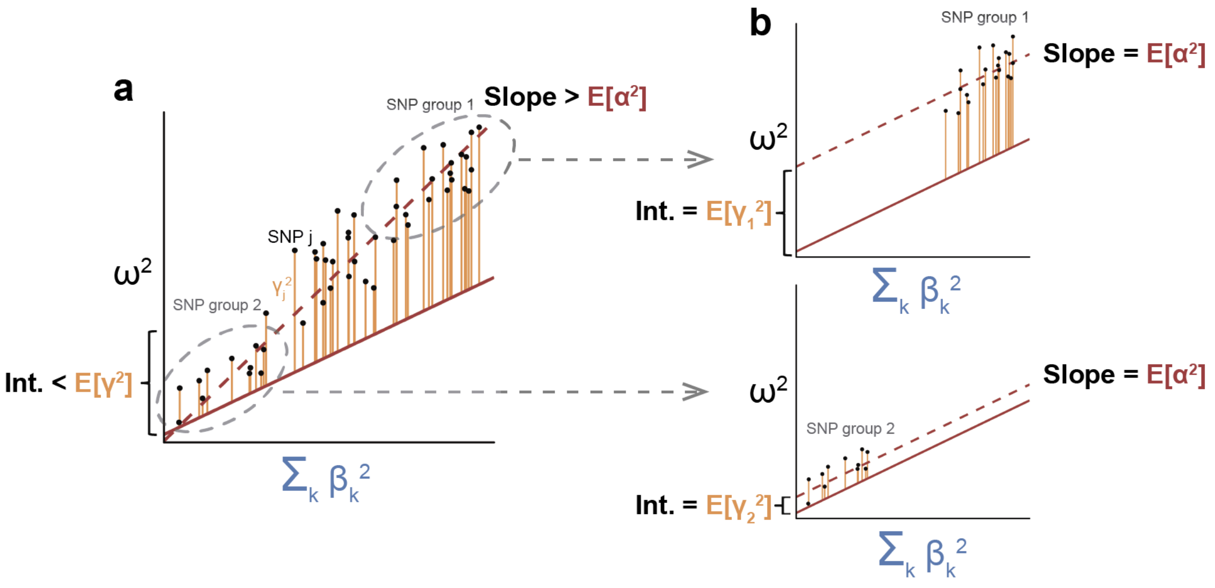


**Supplementary Figure 30. Illustration of impact of violations to pleiotropy-eQTL effect size independence on estimates of $E[\alpha^2]$.** See "Model assumptions" for context for this figure. In the figure, we depict a scenario where the magnitude of $\gamma$ is positively correlated with the magnitude of $\beta$. (**a**) If we perform the regression using all SNPs, the slope from the regression will be upwardly biased relative to the true $E[\alpha^2]$. (**b**) If we stratify the regression across SNPs by the magnitude of their eQTL effect sizes, we can obtain an approximately unbiased estimate of $E[\alpha^2]$.

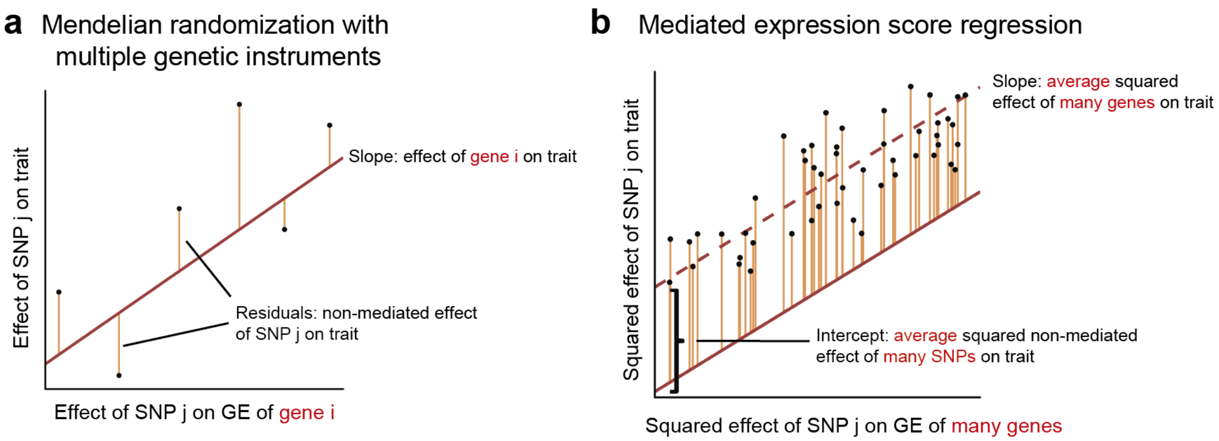


**Supplementary Figure 31. Comparison between Mendelian randomization with multiple genetic variants and MESC.**

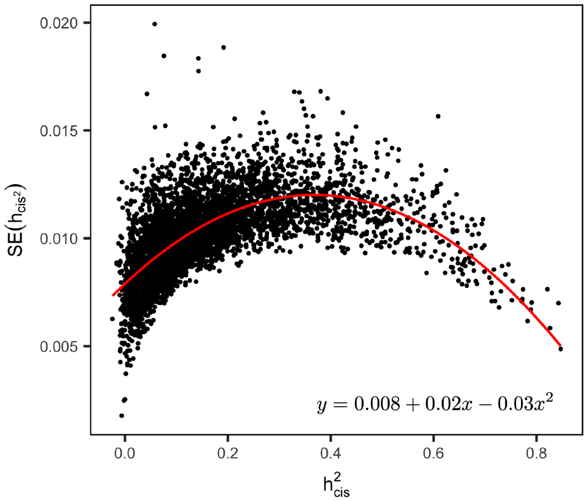


**Supplementary Figure 32. Standard errors of expression cis-heritability estimates across GTEx tissues.** Expression cis-heritability estimates are obtained for each gene in each of 48 GTEx tissues using GCTA. Standard error represents the mean standard error of expression cis-heritability estimates for each gene across all 48 tissues. Red line denotes the best quadratic fit.

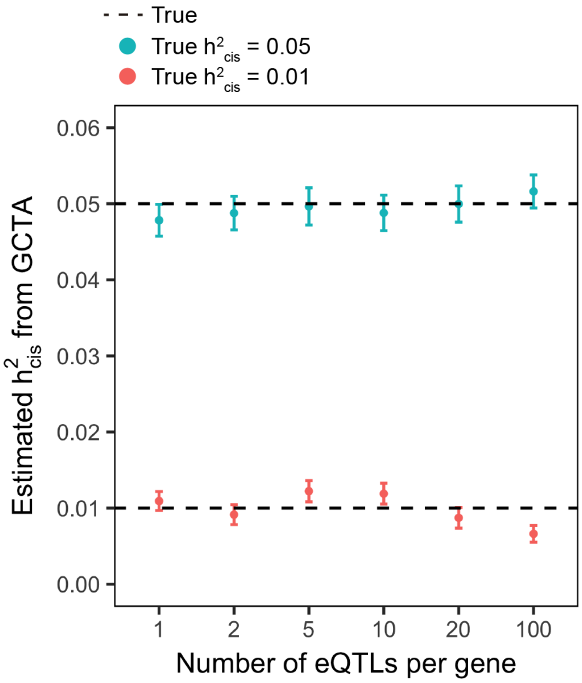


**Supplementary Figure 33. GCTA estimates of heritability under simulated cis-eQTL genetic architectures.** We simulated cis-eQTLs for a gene by selecting $x$ random SNPs within a random 1 Mb window on chromosome 1 (restricting to Hapmap3 SNPs). One cis-eQTL was randomly selected to be the lead eQTL with effect size drawn from $\mathcal{N}(0,\, 0.8h^2_{cis})$. The remaining $x-1$ cis-eQTLs had effect sizes drawn from $\mathcal{N}(0,\, 0.2h^2_{cis}/(x-1))$. Expression phenotypes for the gene were simulated for 1000 individuals (genotypes randomly selected from UK Biobank) with environmental noise drawn from $\mathcal{N}(0,\, 1-h^2_{cis})$. GCTA was used to predict $h^2_{cis}$ from the expression phenotypes and genotypes within the 1 Mb window. Error bars represent mean standard errors across 100 simulations.

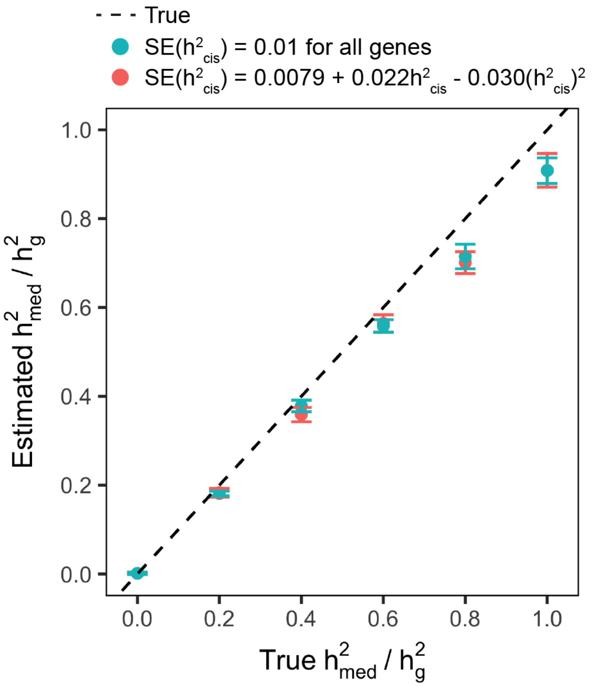


**Supplementary Figure 34.** $h^2_{med}/h^2_g$ **estimates with varying** $SE(h^2_{cis})$ **as a function of** $h^2_{cis}$**.** Simulation was performed in the same manner as in Figure 2a (with expression panel size fixed at 1000). Standard error for $h^2_{cis}$ was simulated from either $\mathcal{N}(0,\, 0.01^2)$ (consistent the mean of empirical $SE(h^2_{cis})$ estimates across all GTEx samples) or from $\mathcal{N}(0,\, (0.0079 + 0.22h^2_{cis} - 0.03(h^2_{cis})^2)^2)$ (consistent with the best quadratic fit line relating empirical $h^2_{cis}$ to $SE(h^2_{cis})$ estimates across all GTEx samples, see Supplementary Figure 32). Error bars represent mean standard errors across 100 simulations.
